# Subcellular and regional localization of mRNA translation in midbrain dopamine neurons

**DOI:** 10.1101/2021.07.30.454065

**Authors:** Benjamin D. Hobson, Linghao Kong, Maria Florencia Angelo, Ori J. Lieberman, Eugene V. Mosharov, Etienne Herzog, David Sulzer, Peter A. Sims

**Author notes:** Department of Neurology, University of California San Francisco School of Medicine, San Francisco, CA. Co-Senior Author (Etienne Herzog, David Sulzer, Peter A. Sims).

## Abstract

Local translation within excitatory and inhibitory neurons is involved in neuronal development and synaptic plasticity. Despite the extensive dendritic and axonal arborizations of central monoaminergic neurons, the subcellular localization of protein synthesis is not well-characterized in these populations. Here, we investigated mRNA localization and translation in midbrain dopaminergic (mDA) neurons, cells with enormous axonal and dendritic projections, both of which exhibit stimulation-evoked dopamine (DA) release. Using highly-sensitive ribosome-bound RNA-sequencing and imaging approaches in mDA axons, we found no evidence for axonal mRNA localization or translation. In contrast, mDA neuronal dendritic projections into the substantia nigra reticulata (SNr) contain ribosomes and mRNAs encoding the major components of DA synthesis, release, and reuptake machinery. Surprisingly, we also observed dendritic localization of mRNAs encoding synaptic vesicle-related proteins, including those involved in vesicular exocytic fusion. Our results are consistent with a role for local translation in the regulation of DA release from dendrites, but not from axons. Our translatome data further defined a molecular signature of the sparse mDA neurons resident in the SNr, including enrichment of *Atp2a3/SERCA3*, an ER calcium pump previously undescribed in mDA neurons.

## Introduction

Midbrain dopaminergic (mDA) neurons play critical roles in reward processing, movement control, and cognitive function. The wide range of neural systems modulated by dopamine (DA) signaling is matched by an elaborate mDA neuronal cytoarchitecture that includes unmyelinated axons that course through the medial forebrain bundle (MFB) to reach multiple basal ganglia and cortical targets (Björklund and Dunnett, 2007). Remarkably, single mDA neurons of the murine substantia nigra pars compacta (SNc) can innervate nearly the entire volume of the dorsal striatum, with axonal arborizations reaching up to 500 mm in total length that possess 10^4^-10^5^ presynaptic varicosities (Matsuda et al., 2009). mDA neurons of the ventral tegmental area (VTA) send axons to the nucleus accumbens (NAcc) and prefrontal cortex (PFC), a relatively late maturational step in development that continues into adolescence (Hoops and Flores, 2017; Manitt et al., 2011).

As originally reported in back to back papers from the Westerink and Iversen laboratories (Geffen et al., 1976; Korf et al., 1976), mDA neurons also release DA within the midbrain (reviewed in Rice and Patel, 2015 and Cheramy et al., 1981), including from large, ventrally-directed dendrites of SNc neurons that can project over 500 μm into the substantia nigra pars reticulata (SNr) (Geffen et al., 1976; Tepper et al., 1987). Although recent work has begun to identify molecular mechanisms that enable and control DA release in the midbrain (Chen and Rice, 2001; Mendez et al., 2011; Robinson et al., 2019; Witkovsky et al., 2009) and striatum (Banerjee et al., 2020; Liu et al., 2018), it is unclear how mDA neurons localize and maintain DA neurotransmission machinery in both dendritic and axonal compartments. The energetic demands placed on SNc neurons due to their extensive cytoarchitecture may contribute to neurodegeneration in Parkinson’s disease (Bolam and Pissadaki, 2012; Pissadaki and Bolam, 2013; Sulzer, 2007), underscoring the importance of proper protein production and distribution in these neurons.

The subcellular proteome of neurons is regulated in part by local translation. Dendritic protein synthesis plays a critical role in various forms of postsynaptic plasticity, including synaptic potentiation (Kang and Schuman, 1996), late long-term potentiation (Bradshaw et al., 2003; Cracco et al., 2005), and long-term depression (LTD) mediated by Group 1 metabotropic glutamate receptors (mGluRs) (Huber et al., 2000; Lüscher and Huber, 2010). Axonal mRNA localization and protein synthesis are well-established in peripheral neurons and in developing axonal growth cones (reviewed in Crispino et al., 2014; Jung et al., 2012). More recent evidence suggests that local translation occurs in mature central nervous system (CNS) axons (Shigeoka et al., 2016), with evidence for presynaptic protein synthesis in hippocampal cannabinoid-induced presynaptic LTD (Younts et al., 2016), neurotransmitter release at the Calyx of Held synapse (Scarnati et al., 2018), responses to neurotrophins (Hafner et al., 2019), and fear-related learning in cortical-amygdalar axons (Ostroff et al., 2019). While accumulating evidence supports a role for dendritic and axonal protein synthesis in excitatory and inhibitory neurons, less is known about axons engaged in modulatory neurotransmission, particularly those that release monoamine neurotransmitters. Intriguingly, the mRNA encoding tyrosine hydroxylase (TH), the rate-limiting enzyme in catecholamine biosynthesis, is localized to axons of sympathetic neurons *in vitro* (Gervasi et al., 2016). Ablation of an axonal localization motif in the 3’ untranslated region (UTR) of *Th* mRNA decreases axonal TH protein levels as well as release of norepinephrine (Aschrafi et al., 2017). These results suggest that local protein synthesis might regulate DA neurotransmission in the central nervous system.

Here, we present a systematic investigation of mRNA localization and translation within mDA neurons in the mouse brain. Using cell-type specific ribosomal capture (RiboTag) and imaging, we find a striking localization of ribosomes and mRNAs encoding DA transmission machinery within dopaminergic dendrites. In contrast, RiboTag studies provided no evidence of translating mRNAs in mDA striatal axons. Fluorescence-activated synaptosome sorting (FASS) of dopaminergic synaptosomes of provided very limited evidence of mRNA localization in dopaminergic axonal varicosities. Surprisingly, we show that mRNAs encoding canonical presynaptic proteins involved in synaptic vesicle neurotransmission are localized within dopaminergic dendrites. Our results reveal the subcellular organization of protein synthesis in mDA neurons, with implications for the regulation of DA neurotransmission in health and disease.

## Results

### DAT^IRES-Cre^:RiboTag mice enable visualization and capture of mDA neuronal ribosomes

To study the subcellular landscape of translation in mDA neurons, we crossed DAT^IRES-Cre^ mice (Bäckman et al., 2006) with RiboTag mice (Sanz et al., 2009) in order to express HA-tagged eukaryotic ribosomal protein L22 (eL22-HA) specifically in mDA neurons (**Figure 1A**). We confirmed high efficiency and specificity of eL22-HA expression in TH^+^ mDA neurons in the SNc and VTA (**Figure S1A**). Anti-HA immunoprecipitation (IP) of ventral midbrain polysome extracts from Cre-positive RiboTag mice (DAT^IRES-Cre/wt^;RiboTag^+/-^) yielded >64-fold enrichment of mDA neuron-specific mRNAs *Th* and *Slc6a3* (DAT), while glial and mRNAs *Gfap*, *Mbp*, and the soluble polyadenylated spike-in standard *ERCC-0096*were depleted based on qRT-PCR (**Figure 1B**). mDA neuron-specific mRNAs were not enriched in control IPs from Cre-negative littermates (DAT^wt/wt^;RiboTag^+/-^).

**Figure 1:**
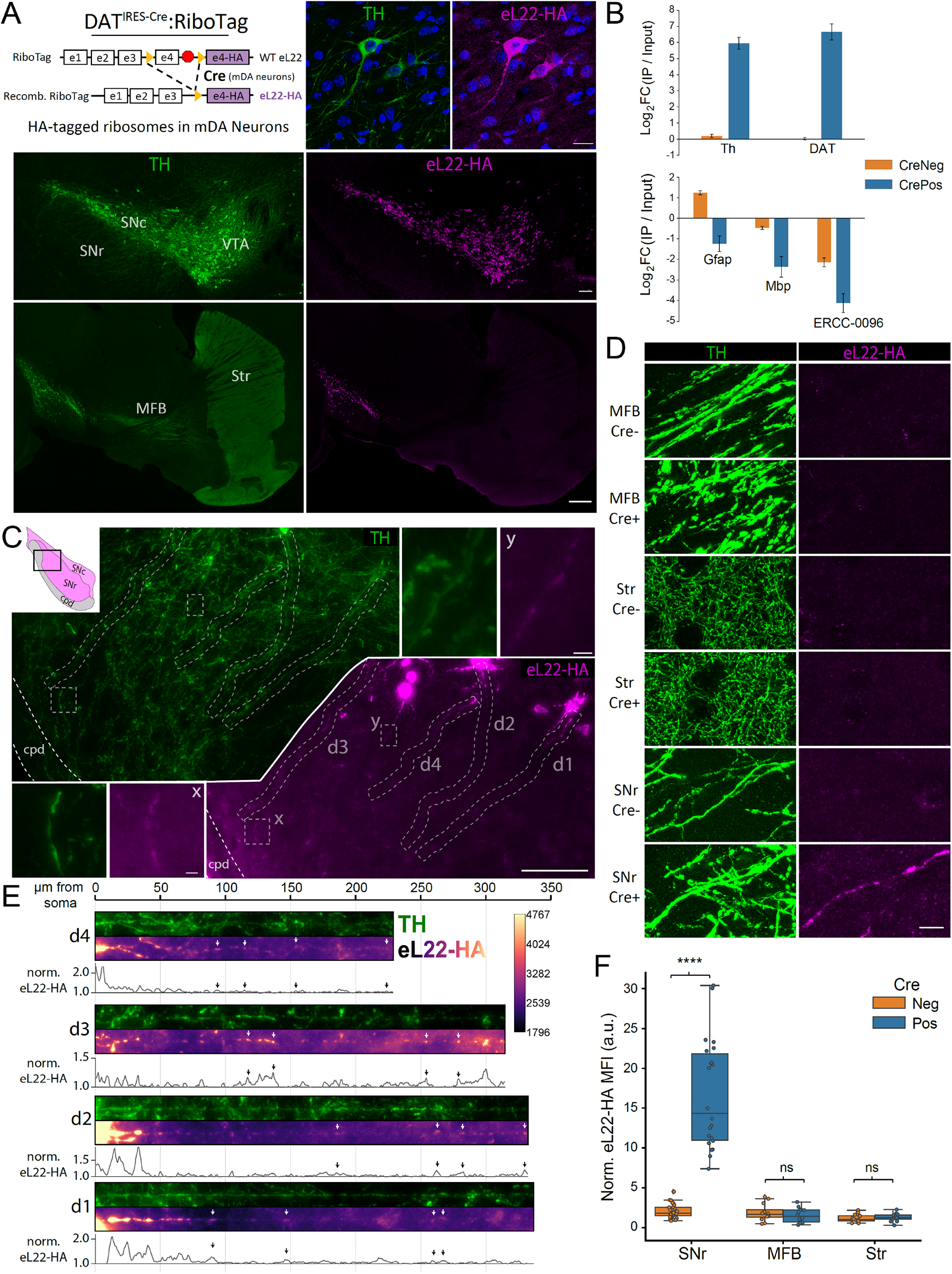
Subcellular distribution of eL22-HA tagged ribosomes in DAT^IRES-Cre^:RiboTag mice. Data in Figure 1 are from mature adult mice (10-14 mo.) **(A)** DAT^IRES-Cre^:RiboTag mice genetics *<u>Upper left</u>:* and immunostaining for tyrosine hydroxylase (TH; green) and eL22-HA (magenta). *<u>Upper right</u>*: mDA neurons in the SNc, DAPI in blue, scale bar: 20 µm. *<u>Middle</u>*: Coronal midbrain section, scale bar: 100 µm. *<u>Lower</u>*: Sagittal section, scale bar: 500 µm. **(B)** qRT-PCR of RiboTag IP and input samples from ventral midbrain of DAT^IRES-Cre^:RiboTag mice (Cre+, n=7) or Cre negative littermates (Cre-, n=5). Cq values were normalized to β-Actin within each input or IP sample. Mean delta-delta Cq (log2 fold changes) +/-SEM are plotted. **(C)** Coronal midbrain section of ventrolateral SNc and SNr stained for TH and eL22-HA (location shown in upper left). Dashed white lines indicate insets labeled x and y or dendrites d1-d4 which are displayed below in panel E. Main image scale bar: 100 µm, ‘x’ inset scale bar: 5 µm, ‘y’ inset scale bar: 5 µm. cpd: cerebral peduncle. **(D)** Representative immunofluorescence of TH and eL22-HA staining in the medial forebrain bundle, striatum, and SNr of Cre-/Cre+ RiboTag mice. Scale bar: 10 µm. **(E)** Straightened dendritic segments d1-d4 shown in panel (C). For each straightened dendrite, eL22-HA fluorescence intensity was normalized to local background and plotted below the images. Arrows indicate ‘hotspots’ of eL22-HA fluorescence along the dendrites. **(F)** Quantification of enhanced eL22-HA immunofluorescence within TH+ neurites in the medial forebrain bundle (axons), striatum (axons), and SNr (dendrites) of Cre- and Cre+ mice. Box and whiskers plots depict the background-normalized eL22-HA mean fluorescence intensity of TH+ pixels within a field of the indicated region (n = 6-10 fields, n = 4 sections, n = 3 mice per each genotype/region).

Consistent with previous studies of recombination in DAT-Cre lines (Bäckman et al., 2006; Lammel et al., 2015; Mingote et al., 2017; Turiault et al., 2007), no expression of eL22-HA was observed in the dorsal striatum, nucleus accumbens, or cortex (**Figure 1C**). DAT-driven expression of tagged ribosomal protein L10a has been previously observed in A12 (Brichta et al., 2015), but this hypothalamic nucleus was readily dissected from the ventral midbrain. Recent work showed that another DAT-Cre line (Ekstrand et al., 2007) drove off-target expression in multiple regions outside the midbrain, including the bed nucleus of stria terminalis (BNST) and the lateral septum (LS) (Papathanou et al., 2019). We found no evidence of eL22-HA expression in the BNST or LS in DAT^IRES-Cre/wt^;RiboTag^+/-^ mice (**Figure S1B**). Similar results were obtained when crossing DAT^IRES-Cre/wt^ mice with Ai9 tdTomato reporter mice (Madisen et al., 2010), with the exception that scattered TH^-^/tdTom^+^ cells were present in the LS (**Figure S1B**). Although these cells were eL22-HA^-^ in DAT^IRES-Cre/wt^;RiboTag^+/-^ mice, nonetheless, we removed all tissue medial to the lateral ventricles, including the LS, BNST, and NAcc shell in striatal dissections (see **Methods**). Thus, in midbrain and striatal dissections of DAT^IRES-Cre/wt^;RiboTag^+/-^ mice, eL22-HA was exclusively derived from mDA neurons.

We leveraged the mDA neuron-specific eL22-HA expression to study the subcellular distribution of ribosomes via immunohistochemistry. Using fluorophore-conjugated secondary antibodies, the majority of eL22-HA labeling was present in soma and proximal dendrites (**Figure 1A**, **Figure S1C**), with no apparent labeling of axons in the MFB or striatum (**Figure 1A**). We used tyramide signal amplification (Adams, 1992; Bobrow et al., 1992) to enhance the anti-HA signal, which produced more intense labeling of dendrites and provided ∼8-fold brighter fluorescence in the soma (**Figure S1D-E**). SNc mDA neurons typically possess three to six long, mostly unbranched dendrites; one or two of these are directed ventrolaterally into the SNr and exhibit the largest diameters and overall length (Juraska et al., 1977; Prensa and Parent, 2001; Tepper et al., 1987). With signal amplification, eL22-HA labeling was apparent within such dendrites in the SNr (**Figure 1C** & **1E**), even at the distal edge near the cerebral peduncles (cpd; **Figure S1F**, *lower*). Co-localization with TH staining confirmed that eL22-HA clusters were scattered throughout the ventral-directed dendrites of SNc mDA neurons (**Figure 1C** & **1E**), which could be distinguished from a few mDA neuronal soma present in the SNr (**Figure S1F**, *upper*). We confirmed the specificity of immunolabeling by quantifying eL22-HA fluorescence in TH^+^ processes in the SNr of Cre-positive and Cre-negative RiboTag mice (**Figure 1D** & **1F**). A previous study found an absence of eGFP-tagged ribosomal protein L10a in dopaminergic axons using conventional immunostaining and low magnification (Brichta et al., 2015). In agreement with that report, even with strong signal amplification and high-resolution confocal imaging, we observed no specific eL22-HA labeling in the MFB or striatum (**Figure 1D** & **1F**). We conclude that the majority of mDA neuronal ribosomes are located in the soma and proximal dendrites, with lower abundance in distal dendrites and exceedingly low levels in axons.

### Sensitive, quantitative capture of dopaminergic ribosomes from regional dissections

To identify translating mRNAs in distinct subcellular compartments of mDA neurons, we conducted RiboTag IP on dissections of four regions (**Figure 2A**): 1) the entire striatal complex, including dorsal and ventral striatum, containing mDA axons, 2) the VTA, which contains mDA neuronal soma and dendrites, 3) the SNc, which contains mDA neuronal soma and horizontally oriented dendrites (Tepper et al., 1987), and 4) the SNr, which contains a few mDA neuronal soma amidst a high density of ventral-projecting dendrites of SNc mDA neurons (**Figure S1G**). As described previously, the exceedingly low yields associated with axonal RiboTag IP necessitate the use of Cre-negative (DAT^wt/wt^;RiboTag^+/-^) mice to control for non-specific binding (Shigeoka et al., 2016). Compared to previous protocols using Protein G Dynabeads, we found that biotinylated anti-HA IgG and streptavidin T1 Dynabeads enabled rapid binding with higher specificity (**Figure S1H**). The total RNA yield from Cre-negative IPs was typically tens to hundreds of picograms and could be accurately estimated using qRT-PCR for beta-actin (*Actb*) (**Figure S1I**).

**Figure 2:**
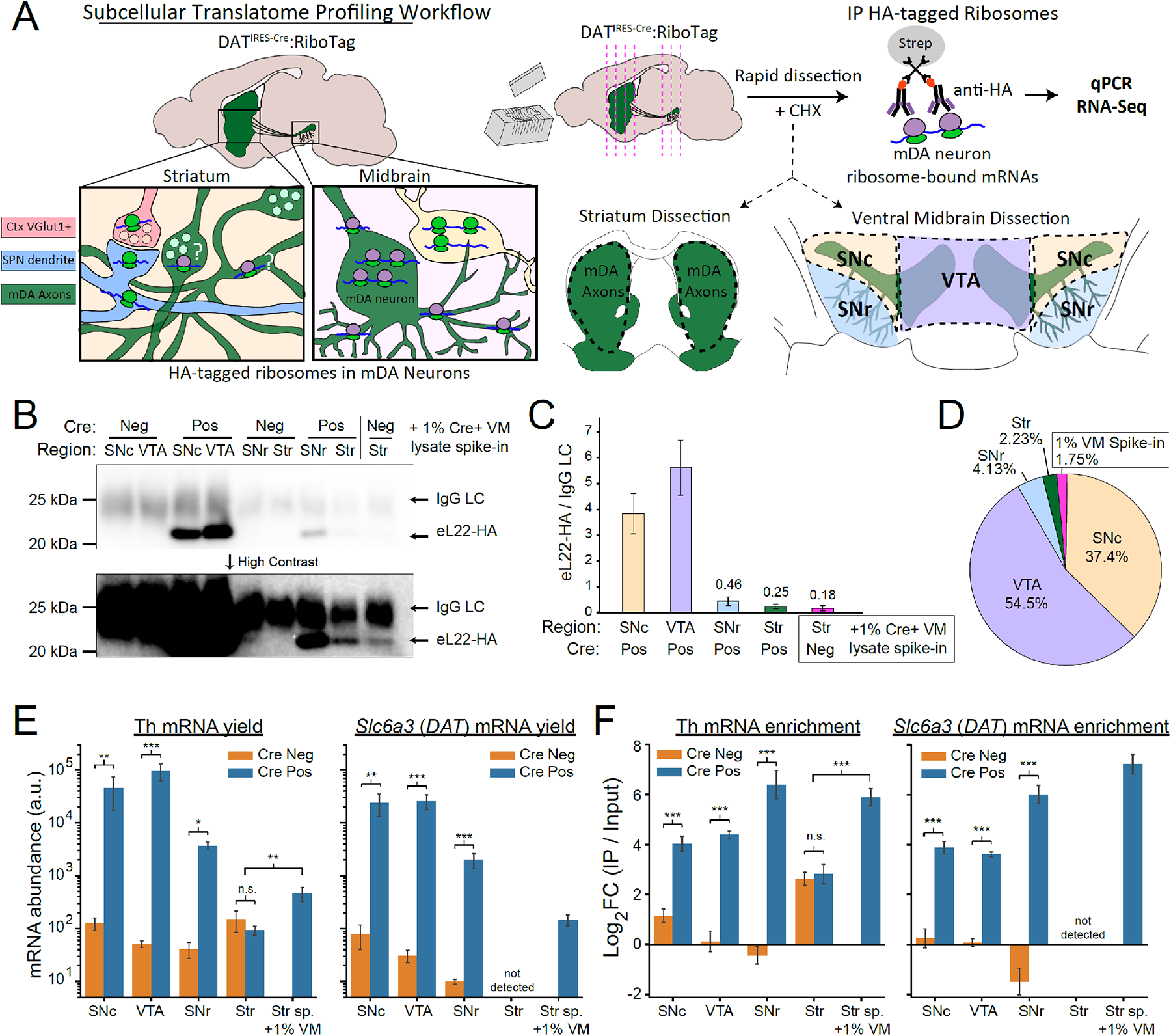
Regional distribution of eL22-HA protein and dopaminergic mRNAs captured by RiboTag IP. Data in Figure 2 are from mature adult mice (10-14 mo.) **(A)** Subcellular translatome profiling workflow schematic. DAT^IRES-Cre^:RiboTag mice (and Cre negative controls) are quickly sliced using an ice-cold brain matrix and razor blades, followed by rapid dissection of the striatum, VTA, SNc, and SNr. Tissues are lysed and subjected to RiboTag IP for capture of mDA neuronal ribosome-bound mRNA. **(B)** Western blot of captured eL22-HA from RiboTag IPs. eL22-HA (23 kDa) is detected just below IgG light chain (LC, ∼25 kDa). Bands in striatal samples and 1% VM spike-in are seen only at high contrast (*lower*). Blot is representative of n=3-4 samples for each condition. **(C)** Quantification of western blot eL22-HA signal intensity, normalized to IgG light chain intensity. Mean +/-SEM are plotted for the indicated regions and genotypes: (SNc Cre+) n=3, (VTA Cre+) n=4, (SNr Cre+) n=3, (Str Cre+) n=3, (Str Cre-/1% VM Cre+ spike-in), n=3. **(D)** Fractional abundance of eL22-HA captured in each region, calculated using normalized eL22-HA intensity from panel C. **(E-F)** qRT-PCR of Th and Slc6a3 (DAT) mRNA in RiboTag IPs from the indicated regions and genotypes (n=3-4 each for region/genotype). * p < 0.05, ** p < 0.01, *** p < 0.001, Welch’s unequal variances t-test. **(E)** mRNA abundance in arbitrary units (2^40 – Cq^). Mean a.u. +/-SEM are plotted. **(F)** RiboTag IP enrichment relative to Input. Cq values were normalized to β-Actin within each input or IP sample. Mean delta-delta Cq (log2 fold changes) +/-SEM are plotted.

Since we were unable to visualize eL22-HA immunofluorescence in the MFB or striatum, we employed western blotting of eL22-HA IPs to estimate eL22-HA abundance in each dissected region. Although prior studies have employed axonal ribosome IP (Ostroff et al., 2019; Shigeoka et al., 2016), these did not include a diluted soma spike-in control to assess the relative abundance of captured ribosomes. To estimate the sensitivity of our IP in the striatum, we included control samples consisting of Cre-negative striatal lysates spiked with 1% of ventral midbrain (VM) lysates from Cre-positive mice. Western blotting of captured eL22-HA revealed prominent bands in VTA and SNc IPs, while faint bands were only visible at high contrast in IPs from the striatum and 1% VM spike-in control (**Figure 2B-C**). Quantification revealed that of all eL22-HA captured, approximately 37.4% was from the SNc, 54.5% from the VTA, 4.13% from the SNr, 2.23% from the striatum, and 1.75% from our 1% VM spike-in control (**Figure 2D**). The striatal eL22-HA abundance was not significantly different from the 1% VM spike-in control. eL22-HA abundance measured by western blot correlates remarkably well with our spike-in control (1% of VM lysate vs. estimated 1.75% eL22-HA) as well as the reported distribution of mDA neurons in C57BL/6J mice (Nelson et al., 1996): ∼8,000 in the SNc (38% of mDA neurons vs. 37.4% of eL22-HA) and ∼10,000 in the VTA (47.6% of mDA neurons vs. estimated 54.5% of eL22-HA). The eL22-HA western blot (**Figure 2B-D**) and histology (**Figure 1**) data correlate well with typical ribosome distribution in neurons and are consistent with very low levels or absence of ribosomes in striatal mDA axons. However, these assays do not establish if the eL22-HA in a given subcellular compartment is derived from translating ribosomes, and so we next analyzed dopaminergic mRNA capture in the same eL22-HA IPs.

qRT-PCR of RiboTag IPs revealed significant Cre-dependent increases in both yield and enrichment of *Th* and *Slc6a3 (DAT)* mRNA in IPs from the SNc and VTA (**Figure 2E-F**). We also found a significant Cre-dependent yield increase for *Th* and *Slc6a3 (DAT)* mRNAs in SNr IPs (**Figure 2E**), and an enrichment of *Th* and *Slc6a3 (DAT)* from the SNr that was higher than in the SNc and VTA (**Figure 2F**). When comparing striatal IPs from Cre-positive and Cre-negative mice, we found no significant differences in *Th* mRNA yield or enrichment, and *Slc6a3 (DAT)* was undetectable in all samples (**Figure 2E-F**). Similar to the results in the SNr, we found >64-fold Cre-dependent enrichment of both *Th* and *Slc6a3 (DAT)* in our 1% VM spike-in control despite a low yield (**Figure 2E-F**). The yield and enrichment of *Th* mRNA were both significantly higher in our 1% VM spike-in controls compared to striatal IPs from Cre-negative and Cre-positive mice (**Figure 2E-F**). These results suggest that *Th* and *Slc6a3 (DAT)* mRNAs are translated in dopaminergic dendrites in the SNr, but not in striatal mDA axons. A complete statistical summary and yield distribution of eL22-HA, *Th* mRNA, and *Slc6a3 (DAT)* mRNA captured by RiboTag IP across all regions are shown in **Table 1**. These data demonstrate sensitive, quantitative RiboTag IP of small quantities (∼1%) of mDA neuronal ribosomes from distinct subcellular compartments.

**Table 1:**
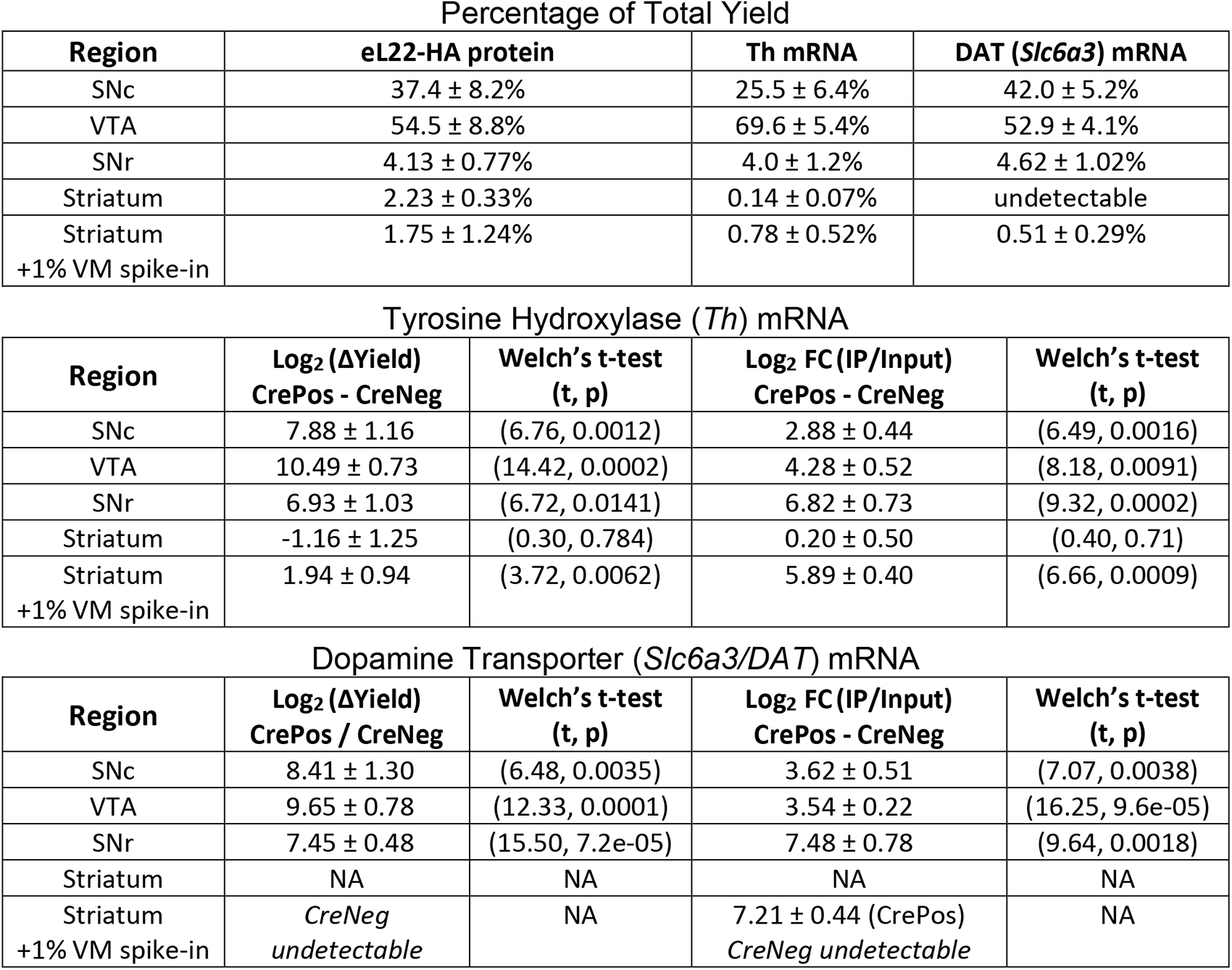
Yield and statistical summary of RiboTag IP capture for eL22-HA, Th mRNA, and *Slc6a3/DAT* mRNA. *<u>Upper:</u>* Mean percentage of total yield ± standard error for each region across all Cre-positive RiboTag IPs for the indicated protein or RNA. Data are derived from **Figure 2B-D** (eL22-HA protein, n=3 for each region) and **Figure 2E-2F** (*Th* and *Slc6a3* mRNA, n=3-4 for each genotype/region). *<u>Lower:</u>* Mean log2 differences in yield (*left*) or enrichment (*right*) ± standard error between Cre-positive and Cre-negative RiboTag IPs for each region/mRNA. Data are derived from **Figure 2E-2F** (*Th* and *Slc6a3* mRNA, n=3-4 for each genotype/region) along with corresponding Welch’s t-test t-statistic and p values.

Percentage of Total Yield
| Region | eL22-HA protein | Th mRNA | DAT (*Slc6a3*) mRNA |
| --- | --- | --- | --- |
| SNC | 37.4 ± 8.2% | 25.5 ± 6.4% | 42.0 ± 5.2% |
| VTA | 54.5 ± 8.8% | 69.6 ± 5.4% | 52.9 ± 4.1% |
| SNr | 4.13 ± 0.77% | 4.0 ± 1.2% | 4.62 ± 1.02% |
| Striatum | 2.23 ± 0.33% | 0.14 ± 0.07% | undetectable |
| Striatum<br>+1% VM spike-in | 1.75 ± 1.24% | 0.78 ± 0.52% | 0.51 ± 0.29% |

Tyrosine Hydroxylase (*Th*) mRNA
| Region | Log<sub>2</sub> (ΔYield)<br>CrePos - CreNeg | Welch's t-test<br>(t, p) | Log<sub>2</sub> FC (IP/Input)<br>CrePos - CreNeg | Welch's t-test<br>(t, p) |
| --- | --- | --- | --- | --- |
| SNC | 7.88 ± 1.16 | (6.76, 0.0012) | 2.88 ± 0.44 | (6.49, 0.0016) |
| VTA | 10.49 ± 0.73 | (14.42, 0.0002) | 4.28 ± 0.52 | (8.18, 0.0091) |
| SNr | 6.93 ± 1.03 | (6.72, 0.0141) | 6.82 ± 0.73 | (9.32, 0.0002) |
| Striatum | -1.16 ± 1.25 | (0.30, 0.784) | 0.20 ± 0.50 | (0.40, 0.71) |
| Striatum<br>+1% VM spike-in | 1.94 ± 0.94 | (3.72, 0.0062) | 5.89 ± 0.40 | (6.66, 0.0009) |

**Table 1: Yield and statistical summary of RiboTag IP capture for eL22-HA, *Th* mRNA, and *Slc6a3/DAT* mRNA**
| Region | Log<sub>2</sub> (ΔYield)<br>CrePos / CreNeg | Welch's t-test<br>(t, p) | Log<sub>2</sub> FC (IP/Input)<br>CrePos - CreNeg | Welch's t-test<br>(t, p) |
| --- | --- | --- | --- | --- |
| SNC | 8.41 ± 1.30 | (6.48, 0.0035) | 3.62 ± 0.51 | (7.07, 0.0038) |
| VTA | 9.65 ± 0.78 | (12.33, 0.0001) | 3.54 ± 0.22 | (16.25, 9.6e-05) |
| SNr | 7.45 ± 0.48 | (15.50, 7.2e-05) | 7.48 ± 0.78 | (9.64, 0.0018) |
| Striatum | NA | NA | NA | NA |
| Striatum<br>+1% VM spike-in | CreNeg<br>undetectable | NA | 7.21 ± 0.44 (CrePos)<br>CreNeg undetectable | NA |

### Lack of Evidence for Axonal Translation in Striatal RiboTag IPs

To identify any mRNAs bound to putative axonal ribosomes, we analyzed the content of striatal RiboTag IPs using RNA-Sequencing (RNA-Seq). Since axonal RiboTag IP yields are exceedingly low, analysis of IPs derived from Cre-negative and Cre-positive samples is critical to control for non-specific binding (Shigeoka et al., 2016). To accommodate picogram samples, we used an low input, pooled library construction strategy that is normally used for single-cell RNA-Seq (scRNA-Seq) in 96-well plates (Snyder et al., 2019) to conduct RNA-Seq of striatal input and IP samples from Cre-negative and Cre-positive mice. This 3’ end RNA-Seq workflow incorporates unique molecular identifier (UMI) barcodes during reverse transcription to mitigate PCR bias. UMI counts provide an estimate of the number of captured mRNA transcripts associated with a specific gene. It is generally appreciated that developing axons have a higher translational capacity than mature axons, which may reflect downregulation of axonal ribosomes following synaptogenesis (Costa et al., 2019). Thus, in addition to middle-aged adult mice (10-14 months, **Figure 2**), we also conducted RiboTag IPs from the striatum of Cre-positive and Cre-negative mice at postnatal ages P0, P7, P14, P21, P31, and P90 (69 mice total, n=2-7 each for Cre-negative and Cre-positive mice at each age). We employed a generalized linear model (GLM) within *DESeq2* (Love et al., 2014) to test whether any mRNAs were significantly enriched in IP vs. Input samples only in Cre-positive mice. Specifically, we used the likelihood ratio test (LRT) to identify genes whose deviance was significantly affected by removal of each term from the full model: ∼*age + genotype + fraction + genotype:fraction* (see **Methods**). In this case, omission of the *genotype:fraction* interaction term (reduced model: ∼*age + genotype + fraction*) tests for a differential effect of *fraction* (IP vs. Input) samples across *genotype* (i.e., ‘Cre-dependent’ enrichment and depletion) (**Figure S2A**).

When analyzing the entire dataset, we found no significant effect of *genotype* for any genes (**Figure 3A-B**), indicating that the single DAT^IRES-Cre^ allele does not alter the striatal transcriptome consistently across *age* and *fraction*. However, there was a statistically significant effect of *fraction* for over 5000 genes (**Figure 3A-B**), demonstrating that IP vs. Input sample differences in these genes are conserved across *genotype* and *age*. This result reflects *age*- and *genotype*-independent bias in the non-specific binding of striatal polysome lysates during RiboTag IP. We also found a statistically significant effect of *age* for over 7000 genes (**Figure 3B**), consistent with major developmental regulation of the striatal transcriptome that is conserved across *fraction* and *genotype*. Critically, we found no significant effect of the *genotype:fraction* interaction for any genes (**Figure 3A-B**), and thus found no statistical evidence for *age*-conserved, Cre-dependent axonal RiboTag IP enrichment of any genes.

**Figure 3:**
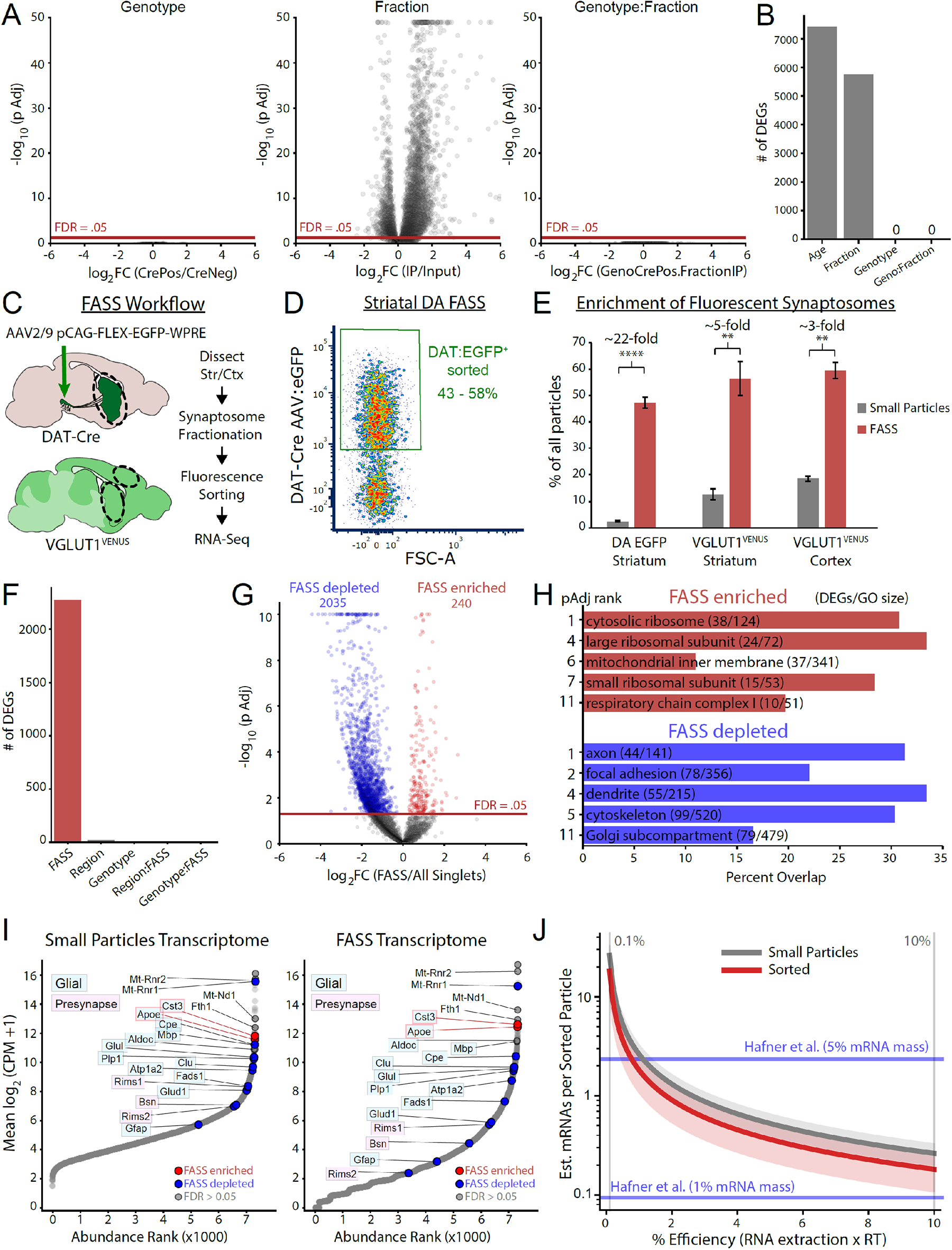
Striatal RiboTag IP and dopaminergic synaptosome sorting provide no evidence of translation in dopamine axons. Data in **Figure 3A-B** are from mice spanning postnatal day P0 to mature adult (10-14 mo.) as specified below. All other data pertaining to synaptosome sorting are from adult mice (3-4 mo.). **(A)** RNA-Seq analysis of striatal RiboTag IP and Input samples at the following ages: (P0, Cre-) n= 6, (P0, Cre+) n=6, (P7, Cre-) n=6, (P7, Cre+) n=6, (P14, Cre-) n=6, (P14, Cre+) n=6, (P21, Cre-) n=6, (P21, Cre+) n=7, (P31, Cre-) n=2, (P31, Cre+), n=2, (P90, Cre-) n=2, (P90, Cre+), n=3, (10-14 mo., Cre-) n=6, (10-14 mo., Cre+) n=4, where each n indicates an IP and corresponding Input sample. Volcano plots are derived from the *DESeq2* LRT, with the indicated terms removed from the following three-factor GLM: *∼Age + Genotype + Fraction + Genotype:Fraction*. Note that a volcano plot from the *Age* LRT (i.e., reduced model omitting the *Age* term: ∼*Genotype + Fraction + Genotype:Fraction*) is not shown since there are many possible *Age* contrasts (i.e., P7 vs. P0, P21 vs. P14, etc.). *<u>Left</u>*: *Genotype* effect across levels of *Fraction* (IP and Input). *<u>Middle</u>*: *Fraction* effect across levels of *Genotype* (Cre-positive and Cre-negative). *<u>Right</u>*: *Genotype:Fraction* effect, which tests for an interaction between *Fraction*and *Genotype*. Log2(GenoCrePos.FractionIP) represents the difference in the *Fraction* effect between genotypes: { Cre-positive log2FC(IP/Input) – Cre-negative log2FC(IP/Input) }. See **Figure S2A** for schematic representation. **(B)** Number of differentially expressed genes (DEGs, BH FDR < 0.05) from the *DESeq2* LRT test for the indicated factors, related to panel A. See **Supplementary File 5** for complete summary of *DESeq2* testing. **(C)** Fluorescence-activated synaptosome sorting (FASS) schematic. Fluorescent synaptosomes are sorted from cortex and/or striatum dissected from mice expressing VGLUT1^VENUS^ or DAT-Cre driven EGFP. **(D)** Representative density plot of EGFP-sorted striatal synaptosomes from DAT-Cre mice expressing EGFP in mDA neurons. Sorted samples are 43-58% EGFP^+^ on re-analysis, which represents a ∼22-fold enrichment from unsorted samples (see panel E). **(E)** Comparison of fluorescent synaptosomes in unsorted and sorted (re-analysis) material for the indicated target sorts. Mean +/-SEM for the %EGFP^+^/VENUS^+^ out of all particles are plotted for the indicated samples: DAT:EGFP Striatum (n=6), VGLUT1^VENUS^ striatum (n=3), and VGLUT1^VENUS^ cortex (n=3). Each sample (n) comprises dissected tissue from 1-3 mice, and both unsorted and sorted (re-analysis) synaptosomes were measured from each sample. ** indicates p < 0.01, **** indicates p < 0.0001, paired t-test. **(F)** Number of differentially expressed genes (DEGs, BH FDR < 0.05) from the *DESeq2*LRT test corresponding to panel E, with the indicated terms removed from the following three-factor generalized linear model: *∼FASS + Region + Genotype + Region:FASS + Genotype:FASS*. See **Supplementary File 6** for complete summary of *DESeq2* testing. **(G)** Volcano plot for *FASS* LRT, comparing sorted samples to all small particles (enriched/depleted genes, BH FDR < 0.05). **(H)** Gene Ontology analysis (*Enrichr*) of FASS-Enriched and FASS-Depleted DEGs, corresponding to panel G. Enriched ontologies were predominantly related to ribosomal subunits and mitochondrial proteins. See **Supplementary File 7** for complete summary. **(I)** Abundance vs. rank plots for all small particles (*left*) and FASS-sorted (*right*) samples. FASS-enriched and -depleted mRNAs are shown in red and blue, respectively. Labeled mRNAs encoding proteins with presynaptic function (pink boxes) were not enriched and were substantially lower in abundance compared to glial-derived mRNAs (light blue boxes). **(J)** Estimated mRNAs per sorted particle as a function of FASS RNA-Seq efficiency, which includes RNA extraction and reverse transcription. Corresponding estimates for whole forebrain VGLUT1^VENUS^ sorted particles from Hafner et al. (2019) are shown in blue, based on 1-5% of total RNA mass corresponding to mRNA. See **Methods** for details.

We also conducted the same analyses at each individual age, again finding no significant effect of *genotype* or *genotype:fraction* interaction, but a significant effect of *fraction* for >1,000 genes at most ages (**Figure S2B**). These findings are consistent with undetectable Cre-dependent mRNA capture in all striatal RiboTag IPs, regardless of age. In further support of these findings, we found no significant Cre-dependent differences in *Th* or *Actb* yield (**Figure S2C**) or in total mRNA yield as determined by total UMIs per sample (**Figure S2D**). Thus, despite our improved RiboTag IP conditions and RNA-Seq protocol with single-cell sensitivity, we found no evidence of ribosome-bound mRNAs derived from DA axons in the striatum. The yield from Cre-negative striatal IPs in our conditions was on the order of 1-50 cells (10-500 pg total RNA estimated via qRT-PCR and 20,000-200,000 UMIs via RNA-Seq), and thus the yield of any axonally-derived ribosomes likely falls well below these background levels. We conclude that translating ribosomes in striatal DA axons are extremely sparse and not amenable to detection using striatal RiboTag IP.

### Lack of dopaminergic mRNA signature in sorted striatal synaptosomes

Another approach to study the presynaptic transcriptome involves fluorescence-activated synaptosome sorting (FASS), which enables enrichment of resealed nerve terminals containing a genetically encoded fluorescent reporter (Biesemann et al., 2014; Hafner et al., 2019; Luquet et al., 2017). Notably, a previous study found that VGLUT1^VENUS^ FASS from whole forebrain enriched mRNAs encoding active zone proteins RIM1 and Bassoon (Hafner et al., 2019). Since these active zone proteins are critically involved in striatal DA release (Liu et al., 2018), we sought to directly interrogate the transcriptome of dopaminergic synaptosomes. We employed our recently developed DA FASS protocol (Pfeffer et al., 2020), where mDA neurons are labeled by injection of AAV expressing Cre-dependent eGFP into DAT-Cre mice (**Figure 3C**). After gating on small particles to avoid synaptosomal aggregates (Biesemann et al., 2014; Hobson and Sims, 2019), we used synaptosomes from WT control mice to establish a background fluorescence threshold for sorting of DAT:EGFP^+^ particles (**Figure S2E**). Reanalysis of DAT:EGFP-sorted synaptosome samples revealed 43-58% EGFP^+^ particles in the sorted material (**Figure 3D**). Compared to 2-6% EGFP^+^ particles in unsorted striatal synaptosomes, sorting thus achieved >20-fold enrichment of DAT:EGFP^+^ synaptosomes (**Figure 3E**, **Figure S2E**). However, as we previously reported, resealed axonal varicosities of mDA neurons can remain stably bound to VGLUT1^+^ presynaptic boutons and other synaptic elements in striatal synaptosome preparations (Pfeffer et al., 2020). To control for mRNA derived from co-enrichment of these other synaptic elements, we also sorted VGLUT1^+^ synaptosomes from the striatum and cortex of VGLUT1^VENUS^ mice (**Figure 3C**). Amongst all striatal synaptosomes, VGLUT1^VENUS+^ particles are more abundant than DAT:EGFP^+^ particles, resulting in a slightly higher sorted purity (50-60%), but substantially lower enrichment (∼5-fold) of VGLUT1^VENUS+^ synaptosomes compared to DAT-EGFP+ synaptosomes (**Figure 3E, Figure S2E**). In addition to DAT:EGFP^+^/VGLUT1^VENUS+^ sorted FASS samples, we also sorted all small particles from each sample to control for bias due to the small particle gating and sorting procedure.

We used the same RNA-Seq protocol as above to characterize the transcriptome of FASS and small particle sorted samples. For each sample of 1.5 - 7.5 million particles, we obtained between 10,000-110,000 UMIs (**Figure S2F**). For striatal samples, DAT:EGFP and VGLUT1^VENUS^ FASS samples yielded significantly fewer UMIs per sorted particle compared to all small particles (**Figure S2G**), although the yield from both sample types was very low (1 UMI per 50-200 sorted particles). Principal components analysis (PCA) revealed that PC1 captured 44% of the variance and clearly separated FASS from small particle sorted samples, but striatal DAT:EGFP FASS samples were not separated from striatal or cortical VGLUT1^VENUS^ FASS samples (**Figure S2H-I**). As above, we used the *DESeq2* LRT to identify genes for which the statistical deviance was significantly affected by omission of terms from the following GLM: ∼*Region + FASS + Genotype + Region:FASS + Genotype:FASS*. We found a significant effect of *FASS* (DAT:EGFP/VGLUT1^VENUS^ sorted vs. all small particles) for >2,000 genes, but no significant effect of *Region:FASS* or *Genotype:FASS* interactions for any genes (**Figure 3F**). These data demonstrate that the FASS transcriptome is distinct from all small particles, but that the VGLUT1^VENUS^ and DAT:EGFP sorted samples are largely indistinguishable, regardless of region. Given the paucity of DA axons in the cortex compared to the striatum, this result argues against any detectable contribution of DA synaptosome-specific mRNAs to the striatal FASS transcriptome. Consistent with this interpretation, most of the 240 genes enriched in FASS samples were enriched by <2-fold (**Figure 3G**), substantially lower than the >20-fold enrichment of DAT-EGFP+ particles in FASS sorted samples. No canonical dopaminergic mRNAs (*Th*, *Slc6a3/DAT*, *Ddc*, and *Slc18a2/VMAT2*) or other DA neuron-specific mRNAs were detected in any FASS samples. Gene ontology (GO) analysis revealed that mRNAs encoding ribosomal proteins and nuclear-encoded mitochondrial proteins are overrepresented amongst FASS-enriched mRNAs (**Figure 3H**), while mRNAs encoding axonal, dendritic, and cytoskeletal proteins were overrepresented amongst FASS-depleted mRNAs.

Nerve terminals are often bound to other particles in synaptosome preparations, which can represent non-specific aggregation (Hobson and Sims, 2019) or native adherence of synaptosomes to postsynaptic densities or other nerve terminals (Pfeffer et al., 2020). Astrocytic processes containing mRNA and ribosomes are also present in synaptosome preparations (Chicurel et al., 1993; Mazaré et al., 2020; Sakers et al., 2017) and are a likely source of mRNA in our FASS samples. We found that despite being depleted relative to all small particles, astrocyte-enriched mRNAs such as *Gfap*, *Glul*, and *Fads1-2* were abundant in our striatal FASS samples (**Figure 3I**). Similarly, we found many oligodendrocyte-enriched mRNAs are still highly abundant in FASS samples, and the microglia- and astrocyte-enriched mRNAs *Cst3* and *Apoe* are enriched in FASS samples (**Figure 3I**). We found no evidence for local translation of active zone proteins in dopaminergic synaptosomes: *Rims1* and *Bsn* mRNAs were very low abundance in small particle sorted samples and were further depleted from FASS samples (**Figure 3I**). The most abundant RNA species in all samples were derived from mitochondrial-encoded genes such as *Mt-Rnr1*, *Mt-Rnr2*, and *Mt-Nd1* (**Figure 3I**). Although mRNAs encoding ribosomal proteins and nuclear-encoded mitochondrial proteins have been observed in axons (Aschrafi et al., 2016; Briese et al., 2016; Gumy et al., 2011; Shigeoka et al., 2016, 2019; Taylor et al., 2009), these mRNAs are also present in dendrites (Fusco et al., 2021; Perez et al., 2021), the synaptic neuropil (Biever et al., 2020; Cajigas et al., 2012), and perisynaptic astrocyte processes (Mazaré et al., 2020). Given the ambiguous cellular and subcellular contribution to the mRNA content of our FASS samples, it is unclear whether any FASS-enriched mRNAs are derived from DA synaptosomes. Since we had to sort millions of synaptosomal particles in order to obtain UMI counts similar to a single cell (**Figure S2F**), we estimated the mRNA content of sorted synaptosomal particles as a function of RNA extraction and reverse transcription efficiency (see **Methods**). Based on reasonable estimates ranging from 1-10% efficiency, we estimate that there are between 0.2 – 2 mRNAs per sorted synaptosomal particle (**Figure 3J**).

Similarly, based on mRNA comprising 1-5% of the total RNA yield from whole forebrain VGLUT1^VENUS^ FASS samples (Hafner et al., 2019), we estimate 0.1 – 2.2 mRNAs per sorted synaptosomal particle (**Figure 3J**). Thus, in addition to the lack of a DAT:EGFP-specific signature and the major contribution of glial and mitochondrial-derived RNAs to the striatal FASS transcriptome, it is possible that many striatal synaptosomes contain no mRNA. Collectively, these data provide no evidence for mRNA localization in dopaminergic synaptosomes.

In a final effort to enrich DA neuron-specific axonal mRNAs, we conducted RiboTag IP and RNA-Seq on striatal synaptosome lysates from Cre-negative and Cre-positive DAT^IRES-Cre^:RiboTag mice. We reasoned that RiboTag IP on striatal synaptosome lysates would provide a lower background than bulk striatal lysates, and could capture some of the FASS-enriched mRNAs bound to putative axonal ribosomes. Similar to whole striatal IPs, we observed no significant effect for *Genotype* or for *Genotype:Fraction* interaction, while hundreds of genes were significantly affected by *Fraction* (**Figure S2J**). We found no significant difference in the estimated total mRNA yield via UMIs (**Figure S2K**), and there was no Cre-dependent RiboTag IP bias for either FASS-enriched or FASS-depleted genes (**Figure S2L-M**). Collectively, these data strongly suggest that mRNAs enriched in DAT:EGFP FASS samples are not derived from ribosomes in dopaminergic varicosities.

### mRNAs encoding dopamine transmission machinery are robustly localized to dopaminergic dendrites, but not axons

The eL22-HA staining in SNr dendrites (**Figure 1**) and the striking enrichment of *Th* and *Slc6a3 (DAT)* mRNAs in SNr RiboTag IPs (**Figure 2**) suggest local translation of these mRNAs in dopaminergic dendrites. Given the presence of occasional mDA neurons in the SNr, we sought to confirm the dendritic localization of *Th* and *Slc6a3 (DAT)* mRNAs using FISH (RNAscope). In addition to dense staining of mDA neuronal soma in the SNc, we observed dispersed *Th* and *Slc6a3/DAT* mRNA puncta throughout the SNr (**Figure S3A**). This staining pattern was not observed for *Ppib* mRNA, a positive control which encodes a ubiquitously expressed endoplasmic reticulum protein and labels all cell bodies (**Figure S3B**). No puncta were observed in sections stained for the bacterial mRNA *DapB*, a negative control (**Figure S3B**).

We optimized the FISH procedure to enable simultaneous immunostaining for TH, which revealed a striking density of *Th* and *Slc6a3 (DAT)* mRNA puncta co-localized with TH^+^ dopaminergic dendrites in the SNr (**Figure 4A**). We found that *Slc18a2* (*vesicular monoamine transporter 2*; *VMAT2*) and *Ddc* (*aromatic l-amino acid decarboxylase*) mRNAs were similarly localized in dopaminergic SNr dendrites (**Figure 4B**). Among these four dopaminergic mRNAs, *Ddc* is the least specific to mDA neurons and is expressed by a variety of cells in the midbrain (**Figure S3C**). Nonetheless, Z-stack confocal imaging enabled us to clearly distinguish *Ddc* puncta within TH^+^ SNr dendrites from those within neighboring soma (**Figure S3D-E**). All four mRNAs were observed within dopaminergic dendrites deep in the SNr, hundreds of microns from SNc soma near the cerebral peduncle (**Figure 4A**, **Figure S3F**).

**Figure 4:**
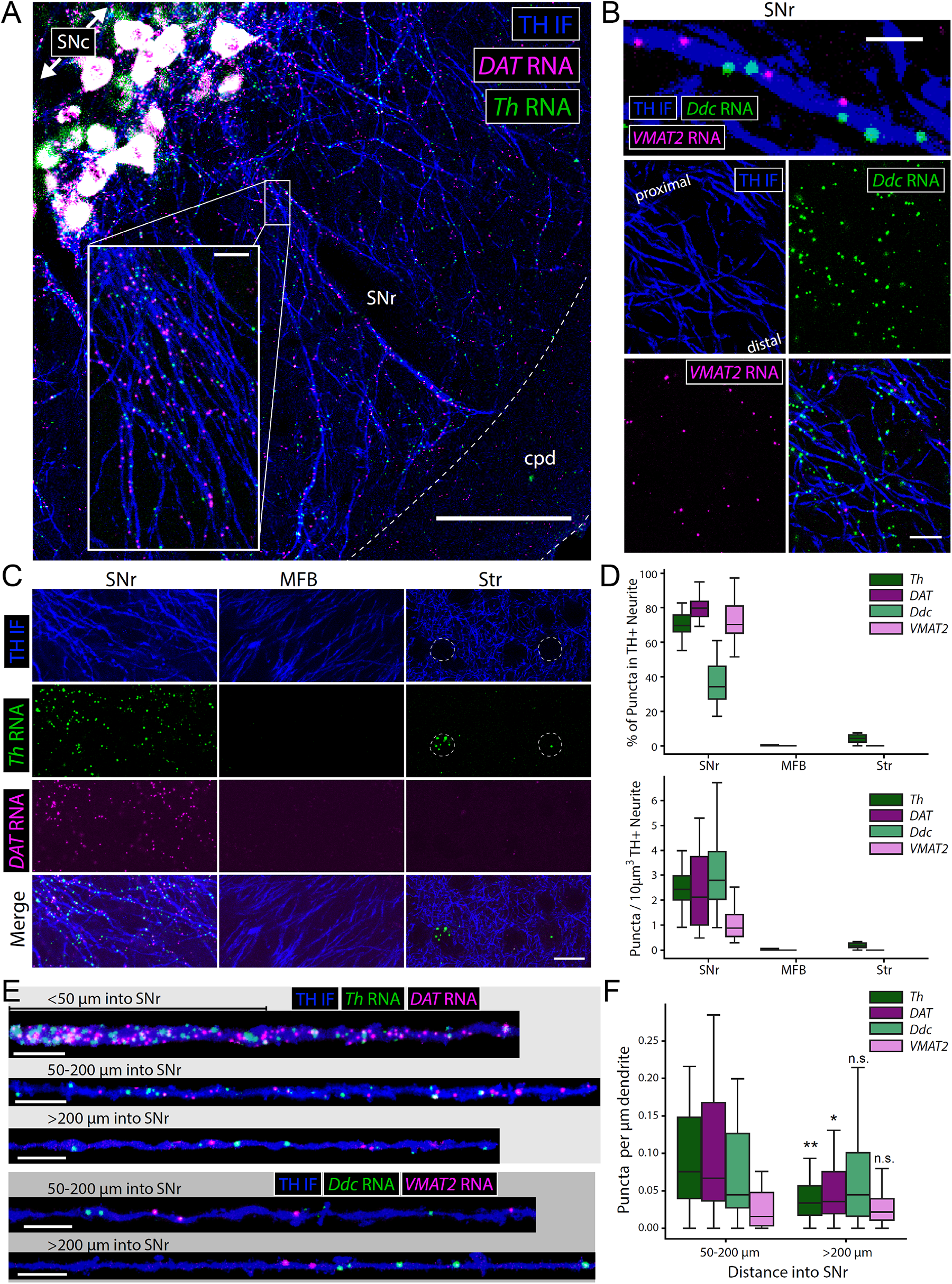
Fluorescence *in situ* hybridization reveals dendritic, but not axonal, localization of dopaminergic mRNAs in the mouse brain. Data in **Figure 4** are from mature adult mice (10-14 mo.) **(A)** Immunostaining for TH and FISH (RNAscope) for *Th* and *Slc6a3* (*DAT*) mRNA in the substantia nigra, cpd: cerebral peduncle. Main image scale bar: 100 µm, inset scale bar: 10 µm. **(B)** Immunostaining for TH and FISH (RNAscope) for *Ddc* and *Slc18a2* (*VMAT2*) mRNA in the murine substantia nigra pars reticulata. Upper image scale bar: 5 µm, lower images scale bar: 15 µm. **(C)** Representative immunostaining for TH and FISH (RNAscope) for *Th* and *Slc6a3* (*DAT*) mRNA in the indicated regions. Scale bar: 15 µm. **(D)** Quantification of RNA puncta within TH^+^ neurites in SNr (dendrites), MFB (axons), and striatum (axons) of mice, related to panel C. Box and whiskers plots represent the specified metric for RNA puncta within a given field in each region: (SNr, *Th*) n=19 fields from 6 sections from 6 mice, (SNr, *Slc6a3*) n=19 fields from 6 sections from 6 mice, (SNr, *Ddc*) n=21 fields from 5 sections from 5 mice, (SNr, *Slc18a2*) n=21 fields from 5 sections from 5 mice, (MFB, *Th*) n=8 fields from 6 sections from 6 mice, (MFB, *Slc6a3*) n=8 fields from 6 sections from 6 mice, (Str, *Th*) n=8 fields from 6 sections from 6 mice, (Str, *Slc6a3*) n=8 fields from 6 sections from 6 mice. *Upper*: % of puncta within TH^+^ neurite. *Lower*: puncta per 10 µm^3^ of TH^+^ neurite. **(E)** Representative segments of TH^+^ dendrite and indicated RNAs at various distances into the SNr. Scale bars: 10 µm. **(F)** Quantification of RNA puncta within TH^+^ dendrites at various distances into the SNr, related to panel E. Box and whiskers plots represent puncta per µm for each segmented dendrite in each region: (50-200µm, *Th*) n=32 dendrites from 6 sections from 6 mice, (>200µm, *Th*) n=32 dendrites from 6 sections from 6 mice, (50-200µm, *Slc6a3*) n=32 dendrites from 6 sections from 6 mice, (>200µm, *Slc6a3*) n=32 dendrites from 6 sections from 6 mice, (50-200µm, *Ddc*) n=31 dendrites from 5 sections from 5 mice, (>200µm, *Ddc*) n=38 dendrites from 5 sections from 5 mice, (50-200µm, *Slc18a2*) n=31 dendrites from 5 sections from 5 mice, (>200µm, *Slc18a2*) n=38 dendrites from 5 sections from 5 mice. * indicates p < 0.05, ** indicates p < 0.01 for Mann-Whitney U-Test comparing >200µm to 50-200µm for each individual RNA.

In contrast to cultured sympathetic neurons (Aschrafi et al., 2017; Gervasi et al., 2016) and to dopaminergic SNr dendrites, we found no *Th* mRNA puncta in MFB mDA axons (**Figure 4C**). Based on the absence of catecholamine-synthesizing neurons in the striatum, previous work proposed mDA axons as the source of striatal *Th* mRNA *in vivo* (Melia et al., 1994). In the striatum, we found dense clusters of *Th* mRNA puncta in soma-sized areas devoid of TH-immunopositive axons (**Figure 4C**). Given that *Th* mRNA is expressed within a subset of striatal neurons (Saunders et al., 2018), these clusters likely represent *Th* mRNA^+^ interneurons that release GABA, not DA (Xenias et al., 2015) (**Figure S4A**). Indeed, these *Th* mRNA^+^ neurons occasionally expressed detectable TH immunolabel, although in our hands this was uncommon (**Figure S4B**). To further establish that striatal interneurons are the source of striatal *Th* mRNA, we measured dopaminergic mRNA levels by qRT-PCR in wild-type and Pitx3-KO mice, which display virtually no dopaminergic innervation of the dorsal striatum due to developmental cell death of SNc mDA neurons (Lieberman et al., 2018; Nunes et al., 2003). Despite a >4-fold decrease in *Th*, *Slc6a3/DAT*, and *Slc18a2/VMAT2* mRNA in the ventral midbrain, where VTA mDA neurons are still intact, we found no significant difference in any of these dopaminergic mRNAs in dorsal striatum tissue (**Figure S4C**). Collectively, these data show that *Th* mRNA^+^ striatal interneurons, and not dopaminergic axons, are responsible for the changes in total striatal *Th* mRNA observed following ablation of mDA striatal axons (Melia et al., 1994).

Using stringent criteria to quantify mRNA puncta within TH^+^ neurites (see **Methods**), we found that >70% of all *Th, Slc6a3/DAT*, and *Slc18a2*/*VMAT2* mRNA puncta in the SNr were clearly localized in dopaminergic dendrites (**Figure 4D**). Since *Ddc* is expressed in other midbrain cells, only about a third of *Ddc* mRNA puncta met the co-localization criteria. Meanwhile, <5% of *Th* and *Slc6a3/DAT* mRNA puncta were co-localized within dopaminergic axons in the MFB and striatum (**Figure 4D**). Despite being the least mDA neuron-specific, *Ddc* mRNA was most abundant in dopaminergic SNr dendrites, followed closely by *Th* and *Slc6a3/DAT* (**Figure 4D**). *Slc18a2/VMAT2* mRNA puncta were least abundant in dopaminergic SNr dendrites (**Figure 4D**). In addition to three-dimensional quantification per dendritic volume, we also quantified dopaminergic mRNA abundance along segmented dendrites. We analyzed single dendritic segments in the proximal SNr (50-200 µm from the SNc) or distal SNr (>200 µm from the SNc), since the mRNA abundance within the first 50 µm was often too high for proper puncta quantification (**Figure 4E**). Similar to 3D quantification, the abundance of *Th* and *Slc6a3/DAT* mRNA in proximal dendritic segments was notably higher than *Slc18a2/VMAT2* mRNA (**Figure 4F**). However, *Th* and *Slc6a3/DAT* mRNA abundance significantly declined in distal dendritic segments, while *Ddc* and *Slc18a2/VMAT2* mRNA did not, such that the abundance of all four mRNAs was comparable within distal dendritic segments (**Figure 4F**). We thus conclude that mRNAs encoding dopamine synthesis, release, and reuptake machinery are present throughout mDA dendritic projections into the SNr.

### Regional translatome profiling reveals *Aldh1a1*^+^/*Sox6*^+^ molecular profile of SNr mDA neurons

Although we identified dendritic ribosomes and mRNAs within the SNr, a few mDA neuronal soma are present in this region. To characterize the mDA neuronal translatome in our SNr dissections (**Figure 2**), we conducted full-length total RNA-Seq (see **Methods** for details) of Input and RiboTag IP samples from the SNc, VTA, and SNr. Principal component analysis (PCA) clearly separated IP vs. Input samples along PC1 and Regions along PC2 (**Figure S5A**). Differential expression of IP vs. Input samples from Cre-positive mice using *DESeq2* revealed a core enrichment signature of dopaminergic gene expression in all three regions (**Figure 5A**), including *Th*, *Ddc, Slc18a2/VMAT2*, and *Slc6a3/DAT*, as well as other canonical mDA marker genes such as *Pitx3*, *En1*, *Ret, Gch1, and Dlk1* (**Figure 5A** **& 5C)**. Thousands of genes were enriched or depleted from IP samples from each region (**Figure 5B**), including canonical glial and GABAergic genes (**Figure 5C**). The enrichment of virtually all dopaminergic genes was strikingly higher in SNr IPs than in VTA and SNc IPs (**Figure 5C**). Together with the substantially lower yield from SNr samples (**Figure 2**), these results suggested the possibility that eL22-HA tagged ribosomes derived from a very small number of SNr mDA neurons dominated the RNA content of our SNr RiboTag IPs. To further analyze the anatomical specificity of our dissections and explore this hypothesis, we leveraged recent datasets analyzing RNA expression at the single mDA neuron level. We mapped our data onto the classification scheme recently introduced by Poulin et al. (2020) (**Figure S5B**), which synthesizes single-cell expression data from six studies (Hook et al., 2018; Kramer et al., 2018; La Manno et al., 2016; Poulin et al., 2014; Saunders et al., 2018; Tiklová et al., 2019). Using only 20 marker genes spanning the mDA neuronal clusters proposed by Poulin et al. (2020), we were able to accurately cluster SNc, VTA, and SNr RiboTag IP samples (**Figure S5C**). VTA-enriched marker genes included *Lpl, Tacr3, Calb1*, *Calb2*, *Cck*, *Neurod6*, and *Grp*, which mostly correspond to the *Otx2*^+^/*Aldh1a1*^+^ population. Meanwhile, SNr-enriched marker genes included *Aldh1a1, Sox6, Aldh1a7*, and *Anxa1*, which correspond to the defining markers of the *Aldh1a1*^+^/*Sox6*^+^ ventral-tier SNc mDA neuronal population. Given the enhanced vulnerability of this population in models of Parkinson’s disease (Cai et al., 2014; Liu et al., 2014; Poulin et al., 2014), we investigated the translatome of our SNr RiboTag IP in greater detail.

**Figure 5:**
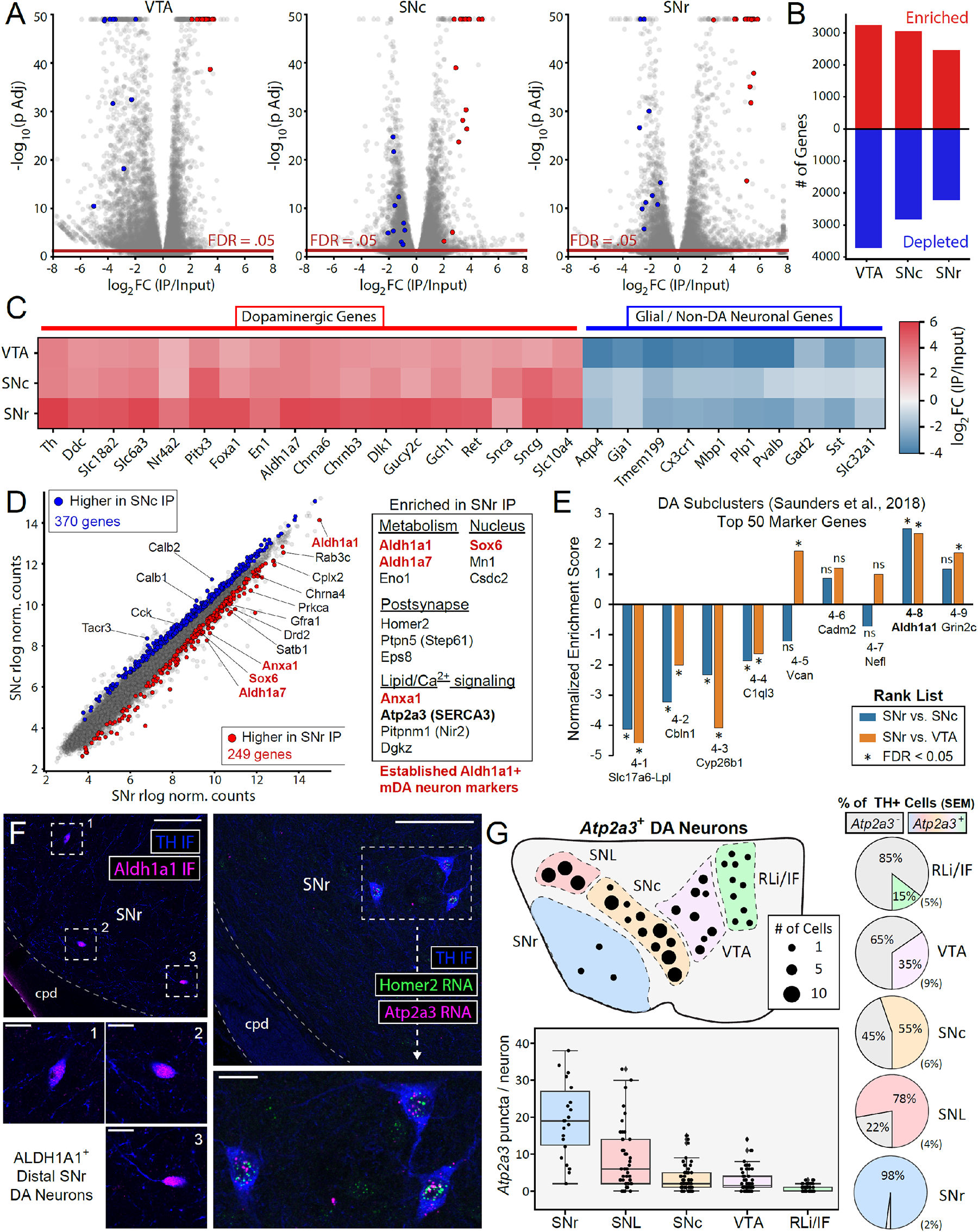
SNr RiboTag translatome reveals *Aldh1a1*^+^ molecular signature and *Atp2a3* (SERCA3) expression in SNr mDA neurons. Data in **Figure 5** are from mature adult mice (10-14 mo.) **(A)** Volcano plots for RiboTag IP vs. Input comparisons (*DESeq2*) in VTA, SNc, and SNr samples (n=4 each). Points colored red or blue correspond to the genes shown in panel C, which are specific to mDA neurons or specific to glia and other neurons, respectively. **(B)** Number of RiboTag IP enriched or depleted genes (FDR < 0.05) from *DESeq2* comparison of IP vs. Input samples in the indicated regions, related to panel A. See **Supplementary File 9** for complete summary of *DESeq2* testing. **(C)** Heatmap of average RiboTag IP enrichment for VTA, SNc, and SNr RiboTag IP samples (n=4 each). Genes specific to mDA neurons (*left*) or glia other neurons (*right*) correspond to red and blue labeled points in the above volcano plots from panel A. **(D)** Average *DESeq2* rlog normalized counts for SNr and SNc RiboTag IP samples (n=4 each). Red and blue genes indicate differential expression between SNr and SNc IP samples, with select genes and cellular function labeled. Genes indicated in red text are canonical markers of Aldh1a1^+^/Sox6^+^ ventral-tier SNc mDA neurons. **(E)** Summary of pre-ranked Gene Set Enrichment Analysis using the top 50 marker genes of each mDA neuronal cluster from Saunders et al. (2018) as gene sets and the rank lists from SNr vs. SNc or SNr vs. VTA RiboTag IPs as indicated. * indicates FDR < 0.05 for the given rank list/gene set combination. **(F)** *<u>Left</u>*: Immunostaining for TH and ALD1A1 reveals distal SNr mDA neurons are ALDH1A1^+^. Scale bar: 100 µm. Inset scale bars: 20 µm. *<u>Right</u>*: FISH for *Atp2a3* (*SERCA3*) and *Homer2* in the SNr reveals the presence of these mRNAs in distal SNr mDA neurons, not in dopaminergic SNr dendrites. Scale bar: 100 µm. Inset scale bar: 20 µm. **(G)** Quantification of *Atp2a3* mRNA expression in mDA neurons using RNA scope and TH immunofluorescence. *<u>Upper left</u>*: Anatomical representation of TH^+^/Atp2a3^+^ neurons in a single hemi-section (15 μm thickness) containing the indicated ventral midbrain regions (approximately -3.2 mm posterior to Bregma). Each dot represents 1, 5, or 10 mDA neurons as indicated, approximating the average of 5 hemi-sections from 4 mice: (RLi/IF) 9.8, (VTA) 31.6, (SNc) 62.8, (SNL) 28.8, and (SNr) 2.8. *<u>Right</u>*: Pie charts indicating the percentage of mDA neurons positive or negative for *Atp2a3* mRNA (TH^+^/*Atp2a3*^-^ or TH^+^/*Atp2a3*^+^) and SEM as indicated, corresponding to the anatomical representation in *Upper left*. Total cell counts for each region (TH+ neurons / TH^+^*Atp2a3*^+^ neurons): RLi/IF (49/338), VTA (158/503), SNc (314/555), SNL (144/187), SNr (28/29). *<u>Lower left</u>*: Box plot of *Atp2a3* mRNA puncta per TH+ neuron in the indicated regions. Data in each region are derived from 5 sections from 4 mice: (SNr) 22 cells, (SNL) 47 cells, (SNc) 64 cells, (VTA) 58 cells, (RLi/IF) 58 cells.

The yield of RiboTag IPs from Cre-positive VTA, SNc, and SNr samples was significantly higher than Cre-negative samples (**Figure S5D**), and the yield of Cre-negative midbrain IPs was generally below the 500 pg required for full length RNA-Seq (**Figure S5E**). Using the ultra-low input, UMI-based RNA-Seq protocol, we identified mRNAs significantly enriched in Cre-negative IP samples relative to Cre-positive IP samples and removed these non-specific binders from subsequent analyses (**Supplementary File 9**). We used *DESeq2* to identify mRNAs differentially expressed between SNr and SNc IPs (**Figure 5D**), with further filtering to select only genes enriched in RiboTag IPs and remove non-specific binders (see **Methods**). We identified 249 genes with higher abundance in SNr compared to SNc IPs, including the ventral-tier SNc mDA neuronal markers noted above (*Aldh1a1, Sox6, Aldh1a7, Anxa1*) (**Figure 5D**) (see **Supplementary File 9** for complete summary). Other SNr-enriched mRNAs encoded proteins involved in lipid/calcium signaling, metabolism, and postsynaptic function (**Figure 5D**). Many of these genes were also higher in SNr compared to VTA IPs (**Figure S5F-H**), suggesting a relative enrichment of *Aldh1a1*^+^/*Sox6*^+^ mDA neurons in our SNr dissection. To further test this interpretation, we conducted pre-ranked gene set enrichment analysis (GSEA) (Subramanian et al., 2005) on the SNr vs. SNc IP and SNr vs. VTA IP differential expression rank lists using the top 50 marker genes of each cluster reported in Saunders et al. (2018). The top 50 markers for *Aldh1a1*^+^ cluster #4-8 were significantly enriched in both SNr vs. SNc and SNr vs. VTA comparisons (**Figure 5E**). Using immunostaining, we found ALDH1A1 expression in all of the few TH^+^ mDA neurons within the proximal SNr (**Figure S6A**) and distal SNr (**Figure 5F**). Finally, we used FISH to study the distribution of several SNr-enriched mRNAs that were not previously described as markers of *Aldh1a1*^+^/*Sox6*^+^ mDA neurons (*Atp2a3*, *Homer2*, *Dgkz*, and *Prkca*). Although we could readily identify FISH puncta within mDA neuronal soma in the proximal and distal SNr (**Figure 5F** and **Figure S6B-D**), we found no evidence of dendritic localization for these mRNAs. Thus, our RiboTag IP predominantly reflects the translatome of mDA neurons within the SNr, demonstrating a molecular signature that significantly overlaps with *Aldh1a1*^+^/*Sox6*^+^ mDA neurons in the ventral-tier SNc (Poulin et al., 2020; Saunders et al., 2018).

Given the importance of autonomous pacemaking activity and cytosolic Ca^2+^ oscillations to mDA neuronal physiology (Zampese and Surmeier, 2020), the SNr enrichment of *Atp2a3* mRNA, which encodes the sarco/endoplasmic reticulum Ca^2+^-ATPase isoform 3 (SERCA3), is of particular interest. SERCA3 is predominantly expressed in hematopoietic and endothelial cells (Bobe et al., 1994; Burk et al., 1989; Wuytack et al., 1994), with cerebellar Purkinje neurons being the only neuronal population previously demonstrated to express SERCA3 (Baba-Aïssa et al., 1996). Our RiboTag data indicate translation of *Atp2a3/SERCA3* in the SNr, SNc, and VTA, although relative abundance was highest in the SNr (**Figure 5E**). We used TH immunostaining and FISH to further define the anatomical distribution of *Atp2a3/SERCA3*-expressing mDA neurons. In addition to the SNr (**Figure 5F**), we found *Atp2a3/SERCA3*^+^ mDA neurons in the substantia nigra pars lateralis (SNL), SNc, VTA, and midline nuclei (rostral linear nucleus and interfascicular nucleus, RLi/IF) (**Figure S6E-H**), although *Atp2a3/SERCA3* expression was substantially lower in the VTA and midline nuclei. The anatomical distribution of *Atp2a3/SERCA3*^+^ mDA neurons in a typical midbrain hemi-section is shown in **Figure 5G**, along with quantification of mRNA puncta per neuron and the percentage of TH^+^ cells expressing *Atp2a3/SERCA3* within each region. While few in number, SNr mDA neurons were virtually all *Atp2a3/SERCA3*^+^ and expressed the highest levels of mRNA per neuron (**Figure 5G**). mDA neurons in the SNL also express high levels of *Atp2a3/SERCA3* and are nearly 80% *Atp2a3/SERCA3*^+^, while *Atp2a3/SERCA3* expression is extremely sparse in the midline nuclei (**Figure 5G**). Future studies will be required to investigate the physiological function of SERCA3 in mDA neurons and explore the consequences of its heterogeneous expression profile.

### Midbrain synaptosome RiboTag IP reveals dendritic localization of mRNAs encoding vesicular release proteins

In addition to regional dissection, another approach to identify translating mRNAs in dendrites is to combine cell type-specific ribosome IP with subcellular fractionation (Ouwenga et al., 2017, 2018). SNc mDA neurons possess dendritic spines, although at lower densities than classical spiny neurons (Hage and Khaliq, 2015; Jang et al., 2015). Critically, resealed dendritic elements within midbrain synaptosome preparations exhibit capacity to uptake and release DA (Hefti and Lichtensteiger, 1978a, 1978b; Silbergeld and Walters, 1979). As an orthogonal approach to SNr dissection, we conducted RiboTag IP on synaptosomes prepared from ventral midbrain tissue (**Figure 6A**). qRT-PCR revealed significantly greater yield of *Th*, *Slc6a3/DAT*, and *Actb* mRNA in Cre-positive IPs compared to Cre-negative controls (**Figure 6B**). Similarly, *Th* and *Slc6a3/DAT* mRNA were enriched roughly nearly 16-fold greater in IP vs. Input comparisons for Cre-positive mice compared to Cre-negative controls (**Figure 6C**). Given the absence of local axon collaterals from mDA neurons (Omelchenko and Sesack, 2009; Tepper et al., 1987), these data demonstrate mDA neuron-specific ribosome capture from postsynaptic elements.

**Figure 6:**
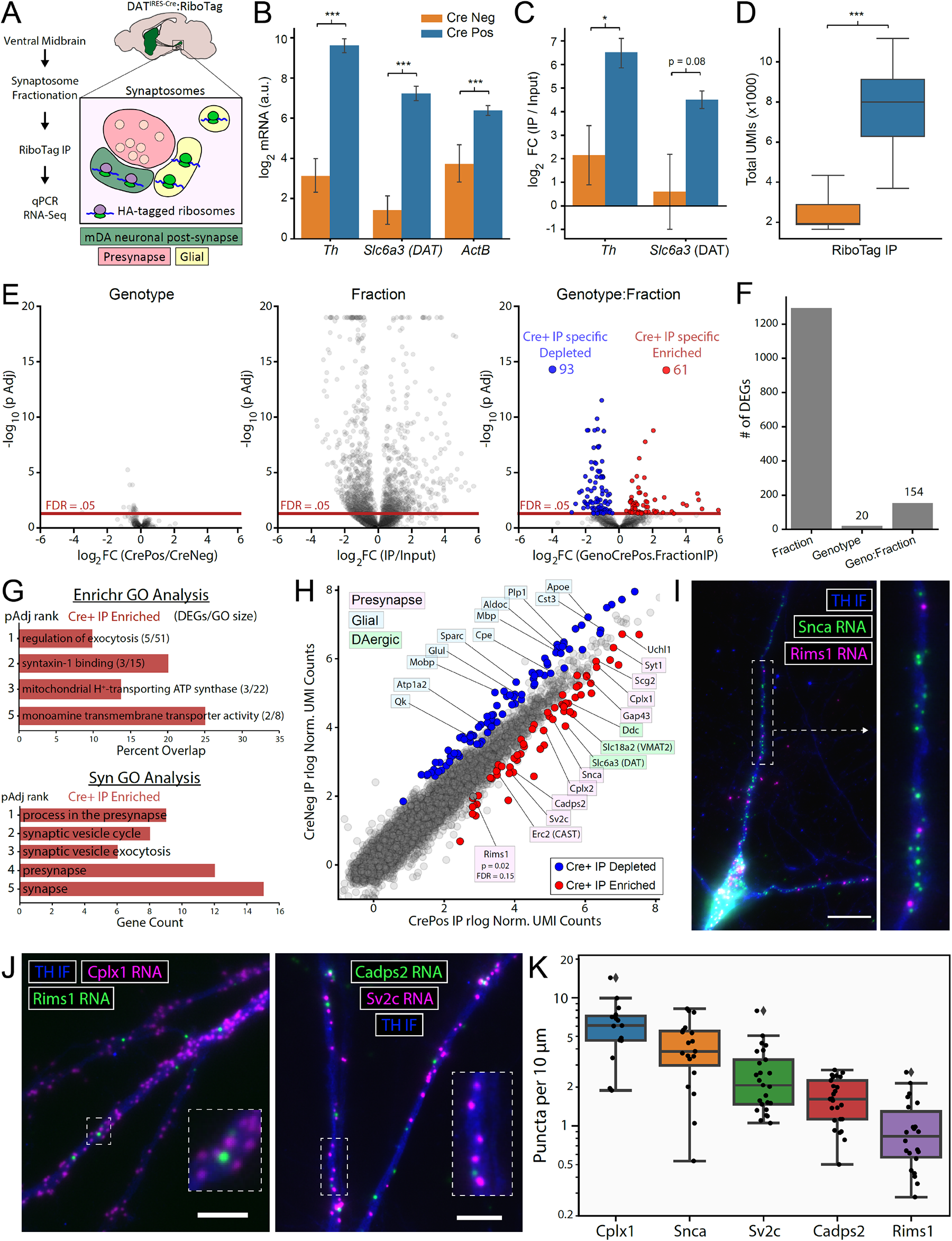
Dendritic localization of mRNAs encoding exocytosis and vesicular release proteins in dopamine neurons. Data in **Figure 6A-H** are from mature adult mice (10-14 mo.), while **Figure 6I-K** are from postnatal mDA neuronal cultures. **(A)** Schematic depicting midbrain synaptosome fractionation and RiboTag IP. **(B)** qRT-PCR measurement of *Th*, *Slc6a3/DAT*, and *Actb* from RiboTag IP of VM synaptosomes from Cre-negative (n=5) and Cre-positive mice (n=6). Log2 mRNA abundance is in arbitrary units (40 – Cq). *** p < 0.001, Welch’s unequal variances t-test. **(C)** qRT-PCR measurement of *Th* and *Slc6a3/DAT* enrichment in RiboTag IP from Cre-negative (n=5) and Cre-positive mice (n=6). Cq values were normalized to β-Actin within each input or IP sample. Mean delta-delta Cq (log2 fold changes) +/-SEM are plotted. * p < 0.05, Welch’s unequal variances t-test. **(D)** UMIs per sample for VM synaptosome RiboTag IPs from Cre-negative (n=6) and Cre-positive mice (n=8). *** p < 0.001, Welch’s unequal variances t-test. **(E)** Volcano plots are derived from the *DESeq2* LRT, with the indicated terms removed from the following two-factor GLM: *∼Genotype + Fraction + Genotype:Fraction*. *<u>Left</u>*: *Genotype* effect across levels of *Fraction* (IP and Input). *<u>Middle</u>*: *Fraction* effect across levels of *Genotype* (Cre-positive and Cre-negative). *<u>Right</u>*: *Genotype:Fraction* effect, which tests for an interaction between *Fraction* and *Genotype*. Log2(GenoCrePos.FractionIP) represents the difference in the *Fraction* effect between genotypes: { Cre-positive log2FC(IP/Input) – Cre-negative log2FC(IP/Input) }. **(F)** Number of differentially expressed genes (DEGs, FDR < 0.05) from the *DESeq2* LRT test for the indicated factors, related to panel E. See **Supplementary File 10** for complete summary of *DESeq2* testing. **(G)** *<u>Upper</u>*: Gene Ontology analysis (*Enrichr*) of Cre-positive IP enriched genes, corresponding to panel E. *<u>Lower</u>*: Gene Ontology analysis (*SynGO*) of Cre-positive IP enriched genes, corresponding to panel E. See **Supplementary File 11** for complete summary of *Enrichr* and *SynGO* analysis. **(H)** Average *DESeq2* rlog normalized UMI counts for Cre-positive (n=8) and Cre-negative (n=6) VM synaptosome IPs. Red and blue genes indicate Cre-positive IP enriched and depleted genes (FDR < 0.05), respectively. Select genes related to dopaminergic (green), glial (blue), and presynaptic function (pink) are labeled. **(I)** Representative immunostaining for TH and FISH (RNAscope) for *Snca* and *Rims1* mRNA in cultured mDA neurons. Dashed white lines indicate the inset shown on the right. Scale bar: 20 µm. **(J)** Representative immunostaining for TH and FISH (RNAscope) for *Cplx1* and *Rims1* mRNA (left) or for *Cadps2* and *Sv2c* mRNA (right) in dendrites of cultured mDA neurons. Dashed white lines indicate the inset shown on the right. Scale bars: 10 µm. **(K)** Quantification of RNA puncta within dendrites of cultured mDA neurons, related to panels I-J. Box and whiskers plots represent RNA puncta per 10µm of dendrite. Data are derived from 2-3 independent cultures, with n dendrites quantified for each mRNA: *Cplx1* (n=16), *Snca* (n=19), *Sv2c* (n=25), *Cadps2* (n=25), *Rims1* (n=35).

We next analyzed the mDA neuronal translatome of these midbrain synaptosome samples using 3’ UMI-based RNA-Seq. Consistent with the Cre-dependent increase in mRNA yield measured by qRT-PCR, Cre-positive IP samples had significantly more UMIs compared to Cre-negative controls (**Figure 6D**). As above, we used the *DESeq2* LRT to identify genes whose statistical deviance was significantly affected by omission of terms from the following GLM: ∼*genotype + fraction + genotype:fraction*. Consistent with a major influence of non-specific binding in all ultra-low input ribosome IP experiments, we found a significant effect of *fraction* for >1,000 genes (**Figure 6E-F**). However, in contrast to bulk striatal and striatal synaptosome RiboTag IP experiments (**Figure 3**), we observed a significant effect of *genotype:fraction* interaction for 154 genes (**Figure 6E-F**). These genes are thus significantly depleted (93) or enriched (61) in IP compared to Input samples *only* in Cre-positive mice (see **Supplementary File 10** for complete summary). Similar to striatal synaptosomes (**Figure 3**), glial mRNAs such as *Apoe*, *Cst3*, *Cpe*, *Glul*, *Mbp*, and *Plp1* were abundant in midbrain synaptosomes; however, these glial mRNAs were uniformly depleted from Cre-positive IP samples (**Figure 6H**). Strikingly, GO analysis of the Cre-positive IP-enriched genes revealed significant over-representation of terms such as ‘regulation of exocytosis’, ‘process in the presynapse’, and ‘synaptic vesicle exocytosis’ (**Figure 6G**). Thus, in addition to canonical dopaminergic mRNAs, we found Cre-dependent enrichment of mRNAs encoding a wide range of proteins with presynaptic function (**Figure 6H**). These included mRNAs encoding proteins involved in synaptic vesicle fusion and recycling (*Erc2/CAST*, *Cplx1*, *Cplx2*, *Syt1*, *Sv2c*, and *Snca*) as well as release from dense core vesicles (*Cadps2/CAPS2* and *Scg2*) (**Figure 6H**). We also observed near-significant enrichment of *Rims1* mRNA, which encodes the active zone protein RIM1 that was recently shown to be critically involved in both axonal and somatodendritic DA release (Liu et al., 2018; Robinson et al., 2019). Many of these mRNAs have also been identified in the dendrites of both glutamatergic and GABAergic hippocampal neurons (Perez et al., 2021) (**Figure S7A**).

Because most mRNAs identified in the midbrain synaptosome IP are not specific to mDA neurons, we validated the dendritic localization of several mRNAs encoding release proteins in postnatally-derived mDA neuron cultures (Rayport et al., 1992), which allowed unambiguous localization of these mRNAs in dopaminergic dendrites using FISH. We first confirmed that cultured mDA neurons recapitulate the dendritic localization of *Th*, *Ddc*, *Slc6a3/DAT*, and *Slc18a2/VMAT2* mRNAs observed in the ventral midbrain (**Figure S7B-E**). Although α-synuclein is abundant in presynaptic terminals, we found dense localization of *Snca* mRNA in dendrites (**Figure 6I**). Similarly, we found a striking density of *Cplx1* (Complexin 1) mRNA in dendrites, along with scattered *Rims1* mRNA (**Figure 6J**). Although Ca^2+^-dependent activator protein of secretion 2 (CAPS2) is involved in catecholamine loading into dense core vesicles in neuroendocrine cells (Brunk et al., 2009; Ratai et al., 2019), CAPS2 has to our knowledge not been characterized in mDA neurons. We found *Cadps2/CAPS2* mRNA within dopaminergic dendrites, along with *Sv2c* mRNA (**Figure 6J**). Synaptic vesicle glycoprotein 2C (SV2C) is involved in axonal DA release (Dunn et al., 2017) and thus may also play a role in somatodendritic DA release. In accordance with the abundance of these mRNAs in the midbrain synaptosome IP (**Figure 6H**), quantification of mRNA puncta in dendrites revealed that *Cplx1* and *Snca* were the most abundant, followed by *Sv2c* and *Cadps2*, and finally *Rims1* (**Figure 6K**). Collectively, these data suggest that local translation of vesicular release proteins may regulate dendritic DA release in mDA neurons.

## Discussion

### Distribution of translation machinery in mDA neurons

We have used multiple immunofluorescence and biochemical approaches to characterize the subcellular distribution of tagged ribosomes in DAT^IRES-Cre^:RiboTag mice, each of which identified the soma as the major site of protein synthesis in mDA neurons. Even with extensive signal amplification, eL22-HA immunoreactivity outside of the mDA neuronal soma was observed only in dendrites, and not in axons (**Figure 1**). Using western blotting, we found that a minor fraction of eL22-HA was present in striatal dissections (**Figure 2D**), but no evidence that this eL22-HA was associated with functional axonal ribosomes. eL22-HA has been found in non-ribosomal fractions of embryonic stem cells (Simsek et al., 2017) and eL22 is known to have extra-ribosomal functions (Bhavsar et al., 2010; Houmani et al., 2009; Ni et al., 2006). We note, however, that visualization of eL22-HA tagged ribosomes in retinal axons required immunoelectron microscopy (Shigeoka et al., 2016), and so we cannot definitively exclude the possibility that extremely low levels of translating ribosomes are present below the limit of detection in our studies.

Many mDA axonal varicosities lack active zone scaffolding proteins and do not release DA upon stimulation (Liu et al., 2018; Pereira et al., 2016), and we initially hypothesized that local translation could be responsible for the determination of active and silent presynaptic sites. However, we found no evidence of axonal mRNA localization, including for presynaptic scaffolding proteins such as *Rims1* or *Bsn*. The paucity of eL22-HA in mDA axons is surprising given their massive axonal arborization. Strikingly, while the striatal axons of SNc mDA neurons likely comprise >90-95% of their cellular volume (e.g., Matsuda et al., 2009) and ∼90% of their cellular protein (Hobson et al., 2021), we found only ∼1% of eL22-HA in the striatum (**Figure 2D**). Even if all of this eL22-HA were presumed to be present in functional ribosomes, the ribosome/protein ratio would be 3-4 orders of magnitude lower in striatal axons compared to mDA neuronal perikarya. Our results are consistent with evidence that a combination of slow and fast axonal transport (Maday et al., 2014; Roy, 2014) of somatically synthesized proteins provides the major source of axonal protein in mDA neurons. Indeed, the massive bioenergetic burden placed on axonal transport systems in mDA neurons may contribute to their demise in Parkinson’s disease (Chu et al., 2012; Sulzer, 2007).

### Lack of dopaminergic mRNA translation in axons

Using a variety of highly sensitive sequencing and imaging approaches, our data indicate that dopaminergic mRNAs (*Th*, *Slc6a3/DAT*, *Ddc*, and *Slc18a2/VMAT2*) are not locally translated in mDA neuronal axons in the mouse brain. One question raised by these results is the apparent discrepancy in axonal localization of *Th* mRNA in mDA neurons compared to cultured sympathetic neurons, where a motif in the 3’ untranslated region (UTR) of *Th* mRNA confers axonal localization and enhances catecholamine release (Aschrafi et al., 2017; Gervasi et al., 2016). Why is *Th* mRNA localization not observed in mDA axons *in vivo*? Recent evidence suggests that *Th* and *Dbh* (dopamine beta-hydroxylase) mRNAs may be transported into sympathetic axons via a shared ribonucleoprotein complex, and that their axonal trafficking is functionally regulated by angiotensin II signaling (Aschrafi et al., 2019). mDA neurons express angiotensin II receptors (Grammatopoulos et al., 2007), but not *Dbh* mRNA, and may lack RNA-binding or trafficking proteins expressed by sympathetic neurons. As the axon of most SNc mDA neurons arises from a dendrite rather than the soma (Juraska et al., 1977; Tepper et al., 1987), we were surprised to find that *Th* mRNA was absent from MFB axons despite extensive localization throughout SNr dendrites (**Figure 3**). Further research is needed to characterize the molecular mechanisms that control *Th* mRNA trafficking in central and peripheral catecholamine neurons.

### Regional heterogeneity in mDA neuronal molecular profile

Integration of single-cell expression studies enables identification of major mDA neuronal subpopulations (Poulin et al., 2020), but most studies have focused on embryonic or early postnatal timepoints (Hook et al., 2018; Kramer et al., 2018; La Manno et al., 2016; Poulin et al., 2014; Tiklová et al., 2019). It has been unclear if the mRNA levels that distinguish mDA neuronal subsets are conserved at the level of translation and at mature ages. Our translational profiling data from middle-aged adult (10-14 mo.) mice recapitulate the regional enrichment of mDA neuronal subset markers (**Figure S5B-C**). Further, we demonstrate that similar to ventral-tier SNc mDA neurons, SNr mDA neurons exhibit an *Aldh1a1*^+^/*Sox6*^+^ molecular signature (**Figure 5**). These data are in accordance with early studies that regarded SNr mDA neurons as displaced SNc mDA neurons (Guyenet and Crane, 1981; van der Kooy, 1979) and later studies demonstrating similar electrophysiological properties of SNc and SNr mDA neurons (Brown et al., 2009; Richards et al., 1997). However, we observed significant differences in expression of genes related to calcium homeostasis in this population. In addition to an established depletion of the Ca^2+^-binding protein calbindin D28k (*Calb1*) (Yamada et al., 1990), we demonstrate the enrichment of *Atp2a3*/*SERCA3* in SNr mDA neurons (**Figure 5D-G**). However, *Atp2a3/SERCA3* expression was also observed in mDA neurons within the SNL, SNc, and VTA. Intriguingly, the only other neurons reported to express SERCA3 are cerebellar Purkinje neurons (Baba-Aïssa et al., 1996), which also exhibit pacemaker firing (Raman and Bean, 1999). SERCA3 has a nearly 5-fold reduced affinity for Ca^2+^ compared to the ubiquitous SERCA2 isoform (Lytton et al., 1992), and may be important for the regulation of cytosolic Ca^2+^ dynamics in these neurons.

### mRNA localization and translation in dopaminergic dendrites

In addition to their massive, complex axonal arborizations in the striatum, SNc mDA neurons must supply SNr projection dendrites with machinery for DA synthesis, release, and reuptake. Immunoelectron microscopy studies of SNc neurons revealed that plasma membrane-associated DAT is sparse in proximal dendrites, with increasing density in distal dendrites, perhaps reflecting specific transport to functional dendritic sites (Hersch et al., 1997; Nirenberg et al., 1996a). Here, we show that *Th*, *Ddc*, *Slc6a3*/*DAT*, and *Slc18a2/VMAT2* mRNAs are localized throughout mDA neuronal dendrites in the SNr (**Figure 4**) and are bound to dopaminergic ribosomes in midbrain synaptosomes (**Figure 6**). In conjunction with vesicular sorting mechanisms (Li et al., 2005), dendritic translation could serve as a means to rapidly modify the local abundance of dopamine transmission machinery. In addition to plasma membranes and postsynaptic densities, DAT is also localized on vesicular and tubular membrane elements within dendrites (Hersch et al., 1997; Nirenberg et al., 1996a). VMAT2 is also present on these structures, termed ‘tubulovesicles’, which are suspected to consist of smooth endoplasmic reticulum (ER) and may represent the site of dendritic DA storage and release (Cheramy et al., 1981; Nirenberg et al., 1996b). The local synthesis of DAT and VMAT2 would require the presence of a form of dendritic ER, and suggests that translation of these DA transport and reuptake proteins may occur directly on ER-related storage and release structures within mDA dendrites. Although the ‘tubulovesicle’ structures in dendrites have been classically described as smooth ER, a large fraction of mRNAs in neurites may be occupied by a single ribosome (Biever et al., 2020), which can elude detection in ultrastructural studies.

Beyond the core dopaminergic machinery, how do mDA neurons manage simultaneous axonal and dendritic localization of vesicular release proteins? For proteins involved in both axonal and somatodendritic DA release, such as RIM1 (Liu et al., 2018; Robinson et al., 2019), our data show that post-translational protein trafficking supplies the vast majority of protein to striatal mDA axons. On the other hand, local translation of RIM1 and other release proteins such as complexins may be important for establishing exocytic release sites in dopaminergic dendrites (**Figure 6**). Postsynaptic complexins are known to regulate AMPA receptor exocytic events during long-term potentiation (Ahmad et al., 2012), although these receptors are recycled on dendritic endosomes (Bredt and Nicoll, 2003) which are not known to be involved in DA release.

Importantly, while evoked release of DA from dendrites was reported nearly a half century ago, the molecular characteristics of the organelles and fusion mechanisms that mediate somatodendritic DA release remain unclear (Rice and Patel, 2015). When expressed in hippocampal neurons, VMAT2 colocalizes with brain-derived neurotrophic factor (BDNF) on vesicles that undergo regulated exocytosis in dendrites (Li et al., 2005). Given that CAPS2 binds to and regulates the depolarization-induced release of neurotrophin-containing vesicles in cerebellar granule cells (Sadakata et al., 2004), the dendritic translation of *Cadps2/CAPS2* mRNA (**Figure 6**) raises the intriguing possibility of CAPS2 involvement in dendritic DA release. More broadly, it is possible that trafficking of synaptic vesicle release proteins in mDA neurons has been optimized to shuttled them into striatal axons, and that such a polarization is incompatible with simultaneous trafficking into dendrites. Local translation in dopaminergic dendrites may provide an alternative mechanism of localization for these proteins, enabling dynamic regulation of proteins at the precise intracellular sites of dendritic exocytosis.

## Materials and Methods

### Animals

All animals were housed in a 12h/12h light/dark cycle with *ad libitum* access to food and water. DAT^IRES-Cre^ mice (JAX #006660, RRID: IMSR_JAX:006660) (Bäckman et al., 2006), Ai9 mice (JAX #007909, RRID: IMSR_JAX:007909) (Madisen et al., 2010) and RiboTag mice (JAX #029977, RRID: IMSR_JAX:029977) (Sanz et al., 2009) were obtained from Jackson Laboratories (Bar Harbor, ME). DAT-Cre mice (MGI:3770172, RRID: MGI:3770172) (Turiault et al., 2007) used in FASS studies were a kind gift from Dr François Tronche. VGLUT1^VENUS^ mice (Slc17a7^tm1.1Ehzg^, RRID: 5297706) used in FASS studies have been previously described (Biesemann et al., 2014; Herzog et al., 2011).

Middle aged adult mice (10-14 months) of both sexes were used in most experiments unless otherwise noted, except for DA FASS studies which used mature adult mice (3-6 months) of both sexes. For RiboTag experiments involving early postnatal ages (P0-P31), mice of both sexes were used and the exact ages are indicated in the text and figure captions. DAT^IRES-Cre^:RiboTag experimental litters were bred by crossing homozygous RiboTag mice (RiboTag^+/+^) with heterozygous DAT^IRES-Cre^ (DAT^IRES-Cre/wt^) mice, yielding litters of DAT^IRES-Cre/wt^;RiboTag^+/-^ (‘Cre positive’) and DAT^wt/wt^;RiboTag^+/-^ (‘Cre negative’) mice. Experimenters were blind to the genotype of mice in these litters throughout animal sacrifice and tissue dissection. Genotyping for the DAT^IRES-Cre^ allele was conducted prior to biochemical experiments using established protocols (Bäckman et al., 2006). All experimental procedures were conducted according to NIH guidelines and were approved by the Institutional Animal Care and Use Committees of Columbia University and the New York State Psychiatric Institute, or according to the European guide for the care and use of laboratory animals and approved by the ethics committee of Bordeaux Universities (CE50) under the APAFIS #21132-2019061314534378v4 (CNRS, France).

### Viral Injections

As previously described (Pfeffer et al., 2020), Stereotaxic injections were performed in heterozygous DAT-*Cre*^+^ mice of either sex at 8 to 9 weeks of age. An Adeno-Associated Virus (AAV1) pCAG-FLEX-EGFP-WPRE from the University of Pennsylvania core facility (Oh et al., 2014) was injected into DAT-*Cre*^+^ mice. Saline injected littermates were used as auto-fluorescence controls. The stereotaxic injections were performed in Isoflurane-anesthetized mice using a 30μl glass micropipette. Injection coordinates for the Substantia Nigra pars compacta (SNc) were anterior/posterior (A/P) - 3.6 mm, lateral (L) +/-1.3mm, dorsal/ventral (D/V) - 4.2mm. Injection coordinates for the Ventral Tegmental (VTA) were A/P - 3.16mm, L +/- 0.6mm; D/V - 4.2mm. A/P and L coordinates are with respect to the *bregma*, whereas D/V coordinates are given with respect to the brain surface. The animals were euthanized after 28 days at the maximal viral EGFP expression. For fluorescence activated synaptosome sorting (FASS) experiments, four to six DAT-*Cre*^+^ mice and one WT mouse were used.

### Antibodies and Reagents

| Name | Manufacturer | Catalog | RRID | Usage |
| --- | --- | --- | --- | --- |
| Rabbit anti-HA | Abcam | ab9110 | RRID:AB_307019 | IHC, 1:1000 |
| Biotinylated Rabbit anti-HA | Abcam | ab26228 | RRID:AB_449023 | IP (NA) |
| Chicken anti-TH | Millipore | AB9702 | RRID:AB_570923 | IHC, 1:500<br>ICC, 1:1000 |
| Rabbit anti-RFP | Rockland | 600-401-379 | RRID:AB_2209751 | IHC, 1:500 |
| Rabbit anti-ALDH1A1 | Abcam | ab52492 | RRID:AB_867566 | IHC, 1:500 |
| Goat anti-Chicken IgY (H+L), Alexa Fluor Plus 488 | ThermoFisher | A-32931TR | RRID:AB_2866499 | IHC/ICC, 1:1000 |
| Goat anti-Rabbit IgG (H+L), Alexa Fluor 647 | ThermoFisher | A32733 | RRID:AB_2633282 | IHC/ICC, 1:1000 |
| Goat anti-Rabbit IgG HRP | Vector Laboratories | PI-1000 | RRID:AB_2336198 | IHC, 3:10,000 |
| Mm-Slc6a3-C1 | ACD Biotechne | 315441 | NA | FISH (NA) |
| Mm-Th-C2 | ACD Biotechne | 317621-C2 | NA | FISH (NA) |
| Mm-Ddc-C3 | ACD Biotechne | 318681-C3 | NA | FISH (NA) |
| Mm-Slc18a2-C1 | ACD Biotechne | 425331 | NA | FISH (NA) |
| Mm-Dgkz-C1 | ACD Biotechne | 534861 | NA | FISH (NA) |
| Mm-Prkca-C2 | ACD Biotechne | 432261-C2 | NA | FISH (NA) |
| Mm-Homer2-O1 | ACD Biotechne | 581231 | NA | FISH (NA) |
| Mm-Atp2a3-C2 | ACD Biotechne | 1049861-C2 | NA | FISH (NA) |
| Mm-Cplx1-C3 | ACD Biotechne | 482531-C3 | NA | FISH (NA) |
| Mm-Snca-C1 | ACD Biotechne | 313281 | NA | FISH (NA) |
| Mm-Cadps2-C3 | ACD Biotechne | 529361-C3 | NA | FISH (NA) |
| Mm-Sv2c-C1 | ACD Biotechne | 545001 | NA | FISH (NA) |
| Mm-Rims1-C2 | ACD Biotechne | 539601-C2 | NA | FISH (NA) |
| TSA Cy3 | Perkin Elmer | NEL744001KT | NA | FISH, 1:1500 |
| TSA Cy5 | Perkin Elmer | NEL745001KT | NA | FISH, 1:1500<br>IHC, 1:7500 |
| Mouse anti-HA | Cell Signaling | 2367S | RRID:AB_10691311 | WB, 1:1000 |
| Goat anti-Mouse HRP | Jackson ImmunoResearch | 115-005-003 | RRID:AB_2338447 | WB, 1:5000 |

IHC: Staining of acute brain slices or sections, ICC: Staining of cultured neurons, WB: western blotting, FISH: Fluorescence *in situ* hybridization.

### Neuronal Cultures

Ventral mesencephalic cultures containing dopaminergic neurons were prepared according to established procedures (Rayport et al., 1992). The ventral midbrain (SN and VTA) from postnatal day 0–2 mice of either sex was dissected, dissociated, and plated on a monolayer of rat cortical astrocytes at the plating density of ∼100,000 cells/cm^2^. Experiments were conducted 14–21 d after plating.

### Immunohistochemistry (IHC)

Mice were anesthetized with euthasol and transcardially perfused with ∼15 mL of 0.9% saline followed by 40-50 mL of ice-cold 4% paraformaldehyde (PFA) in 0.1 M phosphate buffer (PB), pH 7.4. Brains were post-fixed in 4% PFA in 0.1M PB for 6-12 hours at 4°C, washed three times in phosphate buffered saline (PBS), and sectioned at 50 µm on a Leica VT1000S vibratome. Sections were placed in cryoprotectant solution (30% ethylene glycol, 30% glycerol, 0.1M PB, pH 7.4) and stored at −20°C until further use.

Sections were removed from cryoprotectant solution and washed three times in tris-buffered saline (TBS) at room temperature. Sections were then permeabilized in TBS + 0.2% Triton-X 100 for one hour at room temperature, followed by blocking in TBS + 10% normal goat serum (NGS) and 0.3% Triton-X 100 for 1.5 hours at room temperature. Sections were then directly transferred to a pre-chilled solution containing primary antibodies in TBS + 2% NGS + 0.1% Triton-X 100 and incubated for ∼40 hours at 4°C. Sections were washed in TBS + 0.05% Tween 20 (TBS+T) five times over an hour at room temperature. Sections undergoing tyramide signal amplification were treated with 3% hydrogen peroxide in TBS+T for 15 minutes at room temperature, followed by another two washes in TBS+T. Sections were incubated in a solution containing secondary antibodies in TBS + 2% NGS + 0.1% Triton-X 100 at room temperature for 1.5 hours, followed by four washes in TBS+T over 45 minutes at room temperature. Sections undergoing tyramide signal amplification were then incubated in TSA-Cy5 (Perkin Elmer, 1:7500) in the manufacturer’s diluent buffer for 1 hour at room temperature. Following four additional washes in TBS, sections were slide mounted and coverslipped with Fluoromount G (Southern Biotech). See *Antibodies and Reagents*for a complete list of antibodies and concentrations used in this study.

### Tissue Dissection for RiboTag IP

Mice were sacrificed by cervical dislocation and brains were rapidly extracted and submerged in ice-cold 0.32 M sucrose buffer with 5 mM HEPES pH 7.4, 10 mM MgCl2, and 100 µg/mL cycloheximide (CHX). Brains were placed on an ice-cold brain matrix (Zivic Instruments) and separated into 0.5-1.0 mm sections using ice cold razor blades. Striatum was dissected from slices between approximately -0.5 mm to 1.5 mm AP to Bregma. To avoid potential DAT^IRES-Cre^ recombined cells in the lateral septum, a single vertical cut was made descending from the lateral ventricle on each side, and all medial tissue (including lateral septum and nucleus accumbens shell) was discarded. The corpus callosum, cortex, and ventral olfactory tubercle were removed. The remaining dorsal and ventral striatum tissue was flash frozen on liquid nitrogen and stored at -80°C.

Ventral midbrain tissue was dissected from slices between approximately -2.5 mm to -3.75 mm AP to Bregma. First, the cortex, hippocampi, and any hypothalamus or white matter ventral to the midbrain were removed. For whole ventral midbrain tissue dissections, a single horizontal cut was made just dorsal to the rostral linear nucleus and all dorsal tissue was discarded. The remaining tissue containing the SN/VTA was flash frozen on liquid nitrogen and stored at -80°C. For regional dissections, the SNr was first dissected away from the midbrain using a conservative semilunar cut halfway from the edge of the cerebral peduncle to the SNc (see **Figure 2A**). The remaining SNc tissue on either side was separated from the VTA by a vertical cut at the lateral edge of the VTA. All tissues were flash frozen and stored at -80°C.

### Synaptosome Preparation for RiboTag IP

Ventral midbrain or striatal dissections were homogenized in 1 mL of ice-cold 0.32 M sucrose with 5 mM HEPES pH 7.4, 10 mM MgCl2, 100 µg/mL cycloheximide (CHX), 1x EDTA-free protease inhibitors (Roche), and 100 U/ml SUPERaseIN. Nuclei and large debris were cleared at 2,000 xg for 10 minutes at 4°C. The supernatant (S1) was further centrifuged at 7,000 xg for 15 minutes at 4°C to yield the P2 pellet. The supernatant (S2) (cytoplasm and light membranes) was removed from the P2 pellet, which was washed by resuspension in 1 mL of ice-cold 0.32 M sucrose buffer (HEPES, MgCl2, CHX, and inhibitors as above) and re-centrifuged at 10,000xg at 4°C prior to lysis. P2 pellets were lysed in 1 ml of lysis buffer (5 mM HEPES pH 7.4, 150 mM KCl, 10 mM MgCl2, 1% Igepal CA-620, 100 µg/ml cycloheximide, 1x EDTA-free protease inhibitors (Roche), and 100 U/ml SUPERaseIN). After resuspension, samples were incubated at 4°C on a rotor for 15 minutes. The resulting synaptosome lysate was subjected to RiboTag IP as described below.

### Fluorescence-Activated Synaptosome Sorting (FASS)

Synaptosomes were prepared from the striatum or cortex of VGLUT1^venus^ or DAT-Cre eGFP-expressing mice by homogenization in 1ml of ice-cold isosmolar buffer (0.32M sucrose, 4 mM HEPES pH7.4, protease inhibitor cocktail Set 3 EDTA-free (EMD Millipore Corp.)), using a 2 ml-glass-Teflon homogenizer with 12 strokes at 900 rpm. The homogenizer was rinsed with 250 μL of isosmolar buffer and 3 manual strokes and then, the pestle was rinsed with additional 250 μL of isosmolar buffer. The final 1.5ml of homogenate (H) was centrifuged at 1000xg for 5min at 4°C in a benchtop microcentrifuge. The supernatant (S1) was separated from the pellet (P1) and centrifuged at 12,600xg for 8 min at 4°C. The supernatant (S2) was discarded and the synaptosomes-enriched pellet (P2) was resuspended in 0.5 ml of isosmolar buffer and layered on a two-step Ficoll density gradient (900 µL of 7.5% and 900 µL of 13% Ficoll, 4 mM HEPES). The gradient was centrifuged at 50,000xg for 21min at 4°C (Beckman Coulter Optima MAX XP ultracentrifuge with a TL-55 rotor). Sucrose synaptosomes were recovered at the 7.5/13% Ficoll interface using a 0.5 mL syringe.

Ficoll gradient-purified synaptosomes were diluted in PBS containing 1 µg /ml FM4-64 and stored on ice throughout FASS procedures. The FACSAria-II (BD Biosciences) was operated with the following settings: 70 μm Nozzle, sample shaking 300 rpm at 4°C, FSC neutral density (ND) filter 1.0, 488 nm laser on, area scaling 1.18, window extension 0.5, sort precision 0-16-0, FSC (340 V), SSC (488/10 nm, 365V), FITC (EGFP) (530/30 nm, 700 V), PerCP (FM4-64) (675/20 nm, 700 V). Thresholding on FM4-64 was set with a detection threshold at 800. Samples were analyzed and sorted at rates of 15,000-20,000 events/s and flow rate of 3. Data was acquired using BD FACS DIVA 6. Cytometry plots were generated using FCS Express 7 (De Novo Software).

### Fluorescence in situ hybridization (FISH)

For mouse brain tissue and neuronal cultures, FISH was performed using the highly sensitive RNAScope® Multiplex Fluorescent v2 assay (ACD Bio). See *Antibodies and Reagents* for complete list of probes and reagents used in this study. Although most single FISH puncta using this assay are likely single mRNA molecules (Wang et al., 2012), this cannot be definitively determined due to the enzymatic signal amplification and non-diffraction-limited size of the mRNA puncta.

Mouse brain sections were prepared as above, removed from cryoprotectant solution, and washed three times in tris-buffered saline (TBS) at room temperature. Sections were incubated with hydrogen peroxide (ACD) for 15 minutes at room temperature, washed several times in TBS, and then mounted to Superfrost slides (Fisher). Sections were allowed to dry for 10 minutes and a hydrophobic barrier (PAP pen, Vector Labs) was created around the tissue. Tissue was incubated in 50% EtOH, then 70% EtOH, then 100% EtOH for 5 minutes each. Sections were rehydrated in TBS for several minutes, digested with Protease IV (ACD) for 25 minutes at room temperature, and rinsed twice with TBS before proceeding to the RNA Scope Multiplex Fluorescent v2 assay (ACD).

Neuronal cultures were fixed in 4% paraformaldehyde in 0.1M phosphate buffer + 4% sucrose for 10 minutes at room temperature. After several washes in TBS, the dish was filled with methanol pre-chilled to -20°C. Cultures were stored at -20°C for up to 4 weeks prior to FISH. After allowing cultures to room temperature, methanol was replaced with 70% EtOH at room temperature for 2 minutes, then with 50% EtOH for 2 minutes, and then cultures were washed for 10 minutes in TBS. Cultures were treated with hydrogen peroxide (ACD) for 10 minutes at room temperature, followed by Protease III (ACD) diluted 1:15 in TBS for 10 minutes at room temperature, followed by two rinses in TBS before proceeding to the RNA Scope Multiplex Fluorescent v2 assay.

The RNA Scope Multiplex Fluorescent v2 assay was conducted according to the manufacturer’s instructions, with all incubations taking place in a humidified chamber at 40°C. Two 5-minute washes in excess RNA Scope Wash Buffer (ACD) took place between each incubation in sequential order: probes (2-hours), AMP1 (30 minutes), AMP2 (30 minutes), AMP3 (15 minutes), HRP-C1/2/3 (15 minutes), TSA Cy3 (30 minutes), HRP blocker (30 minutes), HRP-C1/2/3 (15 minutes), and TSA Cy5 (30 minutes). Samples were washed twice more in RNA Scope Wash Buffer, then twice more in TBS. Samples were then blocked and immunostained for Tyrosine Hydroxylase as described above. After immunostaining, samples were mounted in Fluoromount G and stored at 4°C for up to 1 week before imaging.

### RiboTag Ribosome Immunoprecipitation

Frozen tissues were thawed on ice in a glass-glass dounce homogenizer with 1-1.5 mL of ice-cold lysis buffer (20 mM HEPES pH 7.4, 150 mM KCl, 10 mM MgCl2, 0.5 mM DTT, 100 µg/mL cycloheximide (CHX), 1x EDTA-free protease inhibitors (Roche), and 100 U/ml SUPERaseIN). Tissues were lysed on ice using 30 strokes each with A and B pestles. Lysates were transferred to pre-chilled Eppendorf tubes and centrifuged at 1,000 xg 4°C for 10 minutes, after which the supernatant was transferred to a new tube. 1/9^th^ the volume of 10% Igepal CA-630 was added to the lysates (final concentration 1%) and they were rotated at 4°C for 15 minutes. Lysates were clarified by centrifuging at 20,000 xg 4°C for 10 minutes and transferred to a new tube. 5% of the lysate was reserved as Input and frozen at -80°C.

1.5 µg (for striatal samples) or 6 µg (for midbrain samples) of biotinylated rabbit anti-HA was then added and the lysates were rotated overnight at 4°C. Streptavidin T1 Dynabeads (ThermoFisher, catalog #65601) were then added to the lysates (5 µL per µg of biotinylated antibody) and rotated for 30 minutes at 4°C. Beads were captured on a magnetic rack and the lysate was discarded. Beads were resuspended in 500 µL of ice-cold high salt buffer (20 mM HEPES pH 7.4, 350 mM KCl, 10 mM MgCl2, 1% Igepal CA-630, 0.5 mM DTT, 100 µg/mL cycloheximide (CHX), 1x EDTA-free protease inhibitors (Roche), and 100 U/mL SUPERaseIN) and transferred to a new tube. Beads were rotated for 30 minutes at 4°C, then captured on a magnetic rack and washed again 3 more times with ice-cold high salt buffer (four washes total over two hours). After the last wash, beads were resuspended in 100 µL of ribosome release buffer (20 mM HEPES pH 7.4, 50 mM EDTA, 100 U/mL SUPERaseIN) and incubated for 10 minutes at room temperature. Beads were captured on a magnetic rack and the eluate containing released mRNA was transferred to a new tube. Beads (with eL22-HA still bound) were flash frozen on liquid nitrogen and stored at -80°C. The aqueous mRNA eluate was purified using the RNEasy MinElute kit (Qiagen, catalog #74204) according to the manufacturer’s instructions. RNA was eluted in 14 µL of nuclease free water supplemented with 20 U/mL SUPERaseIN and stored at -80°C.

### Quantitative Reverse Transcription PCR (qRT-PCR)

For RiboTag IP samples, an equal fraction of captured RNA was reverse transcribed (1.5 µL of the 14 µL elution from RNEasy MinElute purification). For Input or other tissue samples, 20-50 ng of total RNA was reverse transcribed. RNA was reverse transcribed in a 20 µL reaction with 0.5 U of Maxima H Reverse Transcriptase (ThermoFisher, catalog # EP0753) and random hexamers (5 µM, ThermoFisher catalog #SO142).

Quantitative PCR was run with TaqMan Universal Master Mix (ThermoFisher catalog #4440042) and TaqMan FAM-MGB primer/probe sets spanning exon junctions on a BioRad CFX96. The following primer/probe sets were used (ThermoFisher): Mouse *ActB*: Mm01205647_g1, Mouse *Th*: Mm00447557_m1, Mouse *Slc6a3/DAT*: Mm00438388_m1, Mouse *Slc18a2/VMAT2*: Mm00553058_m1, Mouse *Gfap*: Mm01253033_m1, Mouse *Mbp*: Mm01266402_m1, and *ERCC-0096*: Ac03460023_a1. For RiboTag IP samples, an equal fraction of cDNA was used in each reaction. For Input or other tissue samples, 3-5 ng cDNA was used in each reaction.

### Western Blotting

Frozen Streptavidin T1 Dynabeads were thawed and resuspended in 1x LDS sample buffer supplemented with 20 mM DTT. To elute eL22-HA, beads were boiled at 95°C for 5 minutes and then placed onto a magnetic rack. Samples were loaded into 10% Bis-Tris polyacrylamide gels (Invitrogen, ThermoFisher catalog #NP0303BOX) and transferred to PVDF membranes (Immobilon-P, MilliporeSigma, catalog #IPVH00010). Membranes were initially washed for 15 minutes in TBST (1X TBS + 0.1% Tween 20), blocked for an hour in 5% BSA/TBST, and incubated overnight at 4°C with primary antibody in 5% bovine serum albumin/TBST overnight. After primary incubation, membranes were washed three times in TBST prior to incubation with HRP-conjugated secondary antibody in 5% BSA/TBST for one hour at room temperature. After secondary incubation, membranes were washed three times in TBST. Signal was developed using Immobilon enhanced chemiluminescent substrate (Millipore, catalog #WBKLS0500) and imaged on an Azure Biosystems C600 system.

### eL22-HA Image Analysis

10 µm Z stacks of 60x fields of view from the SNr, MFB, and striatum were acquired and collapsed via maximum projection. A binary mask was used to identify pixels in TH+ dendrites and axons. The mean eL22-HA intensity for TH positive pixels was subtracted from the mean eL22-HA intensity for all pixels within each field and is reported in **Figure 1F** as normalized eL22-HA mean fluorescence intensity (MFI).

### FISH Image Analysis

RNA puncta were analyzed using TrackMate (Tinevez et al., 2017). The Laplacian of Gaussian spot detector with estimated blob diameter of 0.5-1.0 μm. Additional filtering was implemented using a combination of quality, contrast, and total intensity as necessary to suppress background spot detection. For each image, the centroid (X,Y,Z) coordinates, diameter, and other quantitative parameters of each punctum were exported for further analysis.

For % co-localization shown in **Figure 4D**, a binary threshold was set for the TH immunofluorescence signal based on two standard deviations above the image background to generate a binary mask of pixels for TH^+^ neurites. The 23 pixels surrounding the TrackMate centroid coordinate of each punctum were analyzed (3 x 3 x 3 cube of pixels excluding the four corner pixels) for overlap with the TH^+^ neurite pixels. Puncta with >60% overlapping pixels were retained as co-localized within TH^+^ neurites. The number of puncta co-localized within TH^+^ neurites was divided by the total volume of TH^+^ pixels in each field, yielding puncta per volume of TH^+^ neurite shown in **Figure 4D**.

For quantification of puncta per µm of dendrite shown in **Figure 4F**, individual dendrites were segmented using the SimpleNeuriteTracer plugin in ImageJ. Dendrites were filled in three-dimensions and exported as a binary mask, from which the (X,Y,Z) coordinates of all pixels in each dendrite were extracted. TrackMate was run once on each original image file, and the number of puncta within each dendrite was determined using the same co-localization analysis as above. The number of puncta in each dendrite was divided by the path length of each dendrite from SimpleNeuriteTracer.

For quantification of *Atp2a3* puncta per neuron shown in **Figure 5G**, individual mDA neuronal soma were segmented in maximum projections of 10 μm Z-stack images using the ImageJ magic wand tool on thresholded TH pixel intensities. Each soma was saved as an ROI, and the (X,Y) coordinates of each ROI were exported. TrackMate was run once on each original image file, and the puncta within each soma were determined using the same co-localization analysis as above.

For quantification of puncta per 10 µm of dendrite shown in **Figure 6K**, dendrites of cultured neurons were manually segmented using *Selection* – *Straighten* in ImageJ. TrackMate was run on each individual image file, and the number of puncta in each dendrite was divided its length.

### Full-length total RNA Sequencing

Full-length total RNA-Seq was conducted using the SMARTer Stranded Total RNA-Seq Kit v3, Pico Input Mammalian (Takara Bio, catalog no. 634485). 1000 pg of total RNA was used for Input samples. For RiboTag IP samples, an estimated 500-1000 pg (via ActB qPCR, see **Figure S1I**) was used. Libraries were constructed according to the manufacturer’s instructions with the following parameters: 1) 4-minute fragmentation prior to reverse transcription, and 2) 14-15 cycles of PCR following ZapR depletion. Unique dual indexes were assigned to each sample, and libraries were pooled at 1 nM following quantification using Qubit dsDNA HS and Agilent 2100 Bioanalyzer High Sensitivity DNA assays. Pooled libraries were sequenced on a NextSeq 500 with 2x75bp paired end reads (HO 150 kit, Illumina).

The first 15 bp of Read 2 (UMI and TSO sequences) were removed using fastx-trimmer (http://hannonlab.cshl.edu/fastx_toolkit/index.html), and paired-end reads then were depleted of rRNA by alignment to mouse 5S, 5.8S, 18S, and 28S rRNA using bowtie2 (Langmead and Salzberg, 2012). rRNA-depleted paired-end reads were then aligned to the mouse genome (GENCODE M25, GRCm38.p6) using STAR 2.6.7a (Dobin et al., 2013). Uniquely mapped reads were then quantified at the exon level using featureCounts version 1.6 (Liao et al., 2014).

### Low input RNA Sequencing with 96-well plate, pooled library construction

The protocol for plate based, 3’ end unique molecular indicator (UMI)-based RNA sequencing of single cells has been described previously (Snyder et al., 2019) and was further modified to accommodate ultra-low input RiboTag IP samples. See **Supplementary File 1** for sequences of all custom primers and oligonucleotides used in this protocol. Briefly, an estimated 20 – 500 pg of total RNA (based on qPCR, see above) for each sample was loaded into the wells of a 96 well plate in a volume of 6 µL of nuclease-free water containing 1 U/µL SUPERaseIN (ThermoFisher). After adding 1.5 µL of 10 µM barcoded RT primer (Integrated DNA Technologies), primer annealing was performed at 72°C for 3 minutes. Reverse transcription was performed by adding 7.5 µL RT mix to each well (2.81 µL of 40% polyethylene glycol 8000, 1.15 µL of 100 mM dNTPs, 3 µL of 5X Maxima H RT Buffer, 0.2 µL of 200 U/µL Maxima H Reverse Transcriptase (ThermoFisher), 0.2 µL of 20 U/µL SUPERaseIN, and 0.15 µL of 100 µM Template Switching Oligo (Integrated DNA Technologies), and 1 µL of nuclease free water). Reverse transcription was performed at 42°C for 90 minutes, followed by 10 cycles of 50°C for 2 minutes, 42°C for 2 minutes, 75°C for 10 minutes, followed by a 4°C hold. Excess primers were removed by adding 2 µL of Exonuclease I mix (1.875U ExoI in water) to each well and incubating at 37°C for 30 minutes, 85°C for 15 minutes, 75°C for 30 seconds, 4°C hold.

All wells were pooled into a single 15-ml falcon tubes and cDNA was purified and concentrated using Dynabeads^™^ MyOne^™^ Silane beads (ThermoFisher) according to the manufacturer’s instructions. The cDNA was split into duplicate reactions containing 25µl cDNA, 25µl 2x HIFI HotStart Ready Mix (Kapa Biosystems), and 0.2M SMART PCR Primer. PCR was run as follows: 37°C for 30 minutes, 85°C for 15 minutes, 75°C for 30 seconds, 4°C hold. Duplicate reactions were combined and purified using 0.7 volumes AMPure XP beads (Beckman Coulter). The amplified cDNA was visualized on an Agilent 2100 Bioanalyzer and quantified using a Qubit II fluorometer (ThermoFisher).

Sequencing libraries were constructed using Nextera XT (Illumina) with modifications. A custom i5 primer was used (NexteraPCR) with 0.6ng input cDNA and 10 cycles of amplification was performed. Unique i7 indexes were used for each plate. After amplification, the library was purified with two rounds of AMPure XP beads, visualized on the Agilent 2100 Bioanalyzer and quantified using the Qubit II fluorometer. Libraries were sequenced on an Illumina NextSeq 500 using the 75 cycle High Output kit (read lengths 26(R1) x 8(i) x 58(R2)). Custom sequencing primers were used for Read 1 (SMRT_R1seq and ILMN_R1seq, see *Antibodies and Reagents*). With each plate we targeted ∼400M reads. Library pools were loaded at 1.8 pM with 20% PhiX (Illumina).

Reads were aligned to the mouse reference genome GRCm38 and transcriptome annotation (Gencode vM10) using the STAR aligner with parameters *–sjdbOverhang 65 –twopassMode Basic* after trimming poly(A)-tails from the 3’-ends. The aligned reads were demultiplexed using the well-identifying barcodes, correcting all single-nucleotide errors. All reads with the same well-identifying barcode, UMI, and gene mapping were collapsed to represent an individual transcript. To correct for sequencing errors in UMIs, we further collapsed UMIs that were within Hamming distance one of another UMI with the same well-identifying barcode and gene. For each 96-well plate, after generating a final list of individual transcripts with unique combinations of well-identifying barcodes, UMIs, and gene mapping, we produced a molecular count matrix for downstream analysis.

### Synaptosome mRNA Content Estimation

For UMI-based estimation of mRNAs per particle shown in **Figure 3J**, the extent to which the Total UMIs per sorted particle underestimates the number of mRNAs per sorted particle was modeled as a function of the efficiency of RNA extraction and reverse transcription:

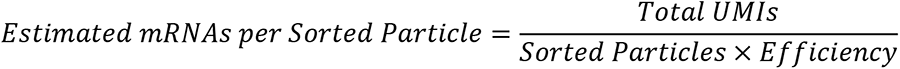

Where *Efficiency* is in decimal form (i.e., 1% efficiency = 0.01, such that the Estimated mRNAs per Sorted Particle is 100-fold more than for 100 % efficiency = 1).

For the estimation of mRNAs per particle based on total RNA measurement of forebrain VGLUT1^venus^ FASS samples (Hafner et al., 2019) shown in **Figure 3J**, the total RNA yield (1-5 ng) from 100 million sorted particles was converted to mRNA estimates as follows:

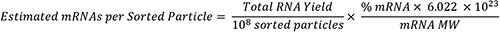

Where *% mRNA* specifies the estimated mass % of mRNA amongst total RNA in decimal form (typically 1-5% or 0.01-0.05), and *mRNA MW* is the average molecular weight of a eukaryotic mRNA in g/mol = (2000 nt x 320.5 g/mol) + 159. In **Figure 3J**, the upper bound corresponds to *Total RNA Yield* = 5 ng and *% mRNA* = 0.05, while the lower bound corresponds to *Total RNA Yield* = 1 ng and *% mRNA* = 0.01.

### RNA-Seq Differential Expression Analysis

Analysis of RiboTag IP and Input sample UMI count matrices shown in **Figure 3**, **Figure S2**, and **Figure 6** (from *Low input RNA Sequencing with 96-well plate, pooled library construction*) was conducted using a generalized linear model (GLM) in *DESeq2* (Love et al., 2014). The likelihood ratio test (LRT) was used to identify genes for which a given term contributed significantly to the likelihood of the full GLM compared to a GLM lacking the given term. In other words, the LRT identifies genes for which a given term adds significant explanatory power to the GLM. The *DESeq2 dds* object was constructed with two- or three-factor models and their interaction terms as specified in the **Results** text. For example, in comparing the full model: ∼*genotype* + *fraction* + *genotype:fraction* vs. the reduced model: ∼*genotype + fraction*, the p-values report on whether the increased likelihood of the full model is greater than by chance if the *genotype:fraction* term truly has no explanatory power. For most LRTs, the log2 Fold Changes specify the contrast between the two levels of the factor (e.g., IP vs. Input for *fraction*, Cre-negative vs. Cre-positive for *genotype*, etc.). For interaction terms, the contrast specifies the difference in log2 Fold Change for the effect of one factor between the levels of the other factor (i.e., for *genotype:fraction*, the difference in IP vs. Input between Cre-negative and Cre-positive samples). Note that the *age* factor in **Figure 3A-B** has multiple levels, and so the p-values do not relate specifically to any single contrast. The log2 Fold Changes specified for the *age* LRT in **Supplementary File 5** are for the contrast P90 vs. P0.

Complete *DESeq2* summary for analyses related to the following figures is found in the corresponding supplementary files:

- **Supplementary File 5** contains *DESeq2* results for bulk striatal RiboTag IP analysis related to **Figure 3**
- **Supplementary File 8** contains *DESeq2* results for striatal synaptosome RiboTag IP analysis related to **Figure S2**
- **Supplementary File 6** contains *DESeq2* results for FASS analysis related to **Figure 3**
- **Supplementary File 10** contains *DESeq2* results for Midbrain synaptosome RiboTag IP analysis related to **Figure 6**

Analysis of midbrain RiboTag IP and Input samples in **Figure 5** and **Figure S5** (from *Full-length total RNA Sequencing*) was conducted in a generalized linear model (GLM) in *DESeq2*. The Wald test was used to make direct comparisons between specific IP samples (e.g., SNr IP vs. VTA IP) or between the IP and Input samples within each region (e.g., SNr IP vs. SNr Input). Downstream filtering of differentially expressed genes (DEGs, FDR < 0.05) to remove non-specific mRNAs is summarized in **Figure S5F**. After identification of DEGs in comparisons of SNr IP vs. VTA or SNc RiboTag IPs, the intersection of SNr-enriched or SNr-depleted genes (relative to SNc/VTA) from these two DEG lists is retained. Next, only genes enriched in SNr IP vs. Input or SNc/VTA IP vs. Input comparisons were retained. Fourth, genes that are significantly higher in Cre-negative IP samples compared to Cre-positive IP samples are removed (non-specific binders). Complete *DESeq2* summary for **Figure 5** and **Figure S5** are found in **Supplementary File 9**.

### Gene Set Enrichment Analysis (GSEA) and Gene Ontology (GO) Analysis

For all GO analyses, a single list of unique genes was used (i.e., differentially expressed genes from *DESeq2* analysis). The GO analyses shown in **Figure 3H** and **Figure 6G** were conducted using web-based Enrichr (Xie et al., 2021) with 2018 GO Terms for Cellular Component, Biological Process, and Molecular Function (Ashburner et al., 2000; Gene Ontology Consortium, 2021). The synaptic gene ontology analysis shown in **Figure 6G** was conducted using SynGO (Koopmans et al., 2019). The results shown in **Figure 5E** employed GSEA (Subramanian et al., 2005) in pre-ranked mode, with the SNr vs. SNc IP or SNr vs. VTA IP *DESeq2* log2 fold change as the rank list and the top 50 markers of each cluster as the gene sets.

### Statistical Analysis

Unless otherwise noted, all statistical analysis and data visualization was conducted in Python using *SciPy*, *Matplotlib*, and *Seaborn* packages. Statistical comparisons were conducted using Welch’s unequal variance t-tests or Mann-Whitney U tests, with number of replicates and other statistical testing information indicated in the figure captions.

### Data and Code Availability

The RNA-Seq data generated in this study are publicly available on the NIH Gene Expression Omnibus (GEO) database as GSE180913. Raw count matrices and differential expression analysis output are provided as supplementary material.

Python and Shell code used for processing of RNA-seq data is accessible at: https://github.com/simslab/DropSeqPipeline8, and Python code for FISH analysis is accessible at: https://github.com/simslab/Neurite_FISH_Quant.

## Supplementary Information

- Supplementary Figures S1-S7
- Supplementary File 1: Custom oligonucleotide and barcoded primer sequences for plate-based, pooled library construction, UMI-based RNA-Seq
- Supplementary File 2: Sample Maps for plate-based UMI RNA-Seq and Clontech v3 RNA-Seq count matrices
- Supplementary File 3: UMI count matrix for plate-based pooled RNA-Seq samples
- Supplementary File 4: Count matrix for full-length RNA-Seq samples (Clontech)
- Supplementary File 5: DESeq2 results related to Figure 3A-B (Bulk Striatum RiboTag IP)
- Supplementary File 6: DESeq2 results related to Figure 3C-I (FASS RNA-Seq)
- Supplementary File 7: Enrichr Gene Ontology analysis related to Figure 3H
- Supplementary File 8: DESeq2 results related to Figure S2J-M (striatal synaptosome RiboTag IP)
- Supplementary File 9: DESeq2 results related to Figure 5 (Midbrain Regional RiboTag IPs)
- Supplementary File 10: DESeq2 results related to Figure 6 (Midbrain synaptosome RiboTag IP)
- Supplementary File 11: Enrichr and SynGO Gene Ontology analysis related to Figure 6G

## Contributions

BDH conceived the overall project with input from PAS and DS. BDH executed all dissection, immunoprecipitation, western blotting, qRT-PCR, and RNA-Seq experiments. MFA and BDH performed FASS experiments with input and supervision from EH. BDH performed histology and immunofluorescence experiments with assistance from OJL and EVM. BDH performed FISH experiments with assistance from LK. BDH conducted all image analysis. BDH conducted RNA-Seq analysis with input from PAS. PAS, DS, and EH supervised the research. BDH wrote the manuscript with input from PAS, DS, and EH. All authors edited, read, and approved the final manuscript.

## Acknowledgements

This work was conducted in collaboration with the JP Sulzberger Columbia Genome Center. This research was funded in part by Aligning Science Across Parkinson’s [ASAP-000375] (DS and PAS) through the Michael J. Fox Foundation for Parkinson’s Research (MJFF). For the purpose of open access, the author has applied a CC BY public copyright license to all Author Accepted Manuscripts arising from this submission. This work was supported by the JPB Foundation (DS). This work was supported by NIH grants F30 DA047775-03 (BDH), R01 NS095435 (DS), R01 DA07418 (DS), and R01 MH122470 (DS). This study benefited from the Agence Nationale de la Recherche consortium fundings (IDEX Bordeaux ANR-10-IDEX-03-02; Labex BRAIN ANR-10-LABX-43 BRAIN; France Bio Imaging ANR-10-INBS-04). EH received salary funding for MFA from Fondation pour la Recherche Médicale (ING20150532192). We thank Vanessa Morales for assistance with animal colony management and Ellen Kanter for assistance with dopamine neuron cultures. Our work benefited from the excellent technical support from the core facilities at Bordeaux university: Bordeaux Imaging Center (CNRS UMS 3420, INSERM US4); Biochemistry and Biophysics of Proteins; Flow cytometry UB’FACSility (CNRS UMS 3427, INSERM US5); animal care & breeding; genotyping.

## References

Adams, J.C. (1992). Biotin amplification of biotin and horseradish peroxidase signals in histochemical stains. J. Histochem. Cytochem. 40, 1457–1463.

Ahmad, M., Polepalli, J.S., Goswami, D., Yang, X., Kaeser-Woo, Y.J., Südhof, T.C., and Malenka, R.C. (2012). Postsynaptic Complexin Controls AMPA Receptor Exocytosis during LTP. Neuron 73, 260–267.

Aschrafi, A., Kar, A.N., Gale, J., Elkahloun, A.G., Vargas, J.-N., Sales, N., Wilson, G., Tompkins, M., Gioio, A.E., and Kaplan, B.B. (2016). A Heterogeneous Population of Nuclear-Encoded Mitochondrial mRNAs Is Present in the Axons of Primary Sympathetic Neurons. Mitochondrion 30, 18–23.

Aschrafi, A., Gioio, A.E., Dong, L., and Kaplan, B.B. (2017). Disruption of the Axonal Trafficking of Tyrosine Hydroxylase mRNA Impairs Catecholamine Biosynthesis in the Axons of Sympathetic Neurons. ENeuro 4.

Aschrafi, A., Berndt, A., Kowalak, J.A., Gale, J.R., Gioio, A.E., and Kaplan, B.B. (2019). Angiotensin II mediates the axonal trafficking of tyrosine hydroxylase and dopamine β-hydroxylase mRNAs and enhances norepinephrine synthesis in primary sympathetic neurons. J. Neurochem. 150, 666–677.

Ashburner, M., Ball, C.A., Blake, J.A., Botstein, D., Butler, H., Cherry, J.M., Davis, A.P., Dolinski, K., Dwight, S.S., Eppig, J.T., et al. (2000). Gene Ontology: tool for the unification of biology. Nat Genet 25, 25–29.

Baba-Aïssa, F., Raeymaekers, L., Wuytack, F., Callewaert, G., Dode, L., Missiaen, L., and Casteels, R. (1996). Purkinje neurons express the SERCA3 isoform of the organellar type Ca2+-transport ATPase. Molecular Brain Research 41, 169–174.

Bäckman, C.M., Malik, N., Zhang, Y., Shan, L., Grinberg, A., Hoffer, B.J., Westphal, H., and Tomac, A.C. (2006). Characterization of a mouse strain expressing Cre recombinase from the 3’ untranslated region of the dopamine transporter locus. Genesis 44, 383–390.

Banerjee, A., Lee, J., Nemcova, P., Liu, C., and Kaeser, P.S. (2020). Synaptotagmin-1 is the Ca2+ sensor for fast striatal dopamine release. Elife 9.

Bhavsar, R.B., Makley, L.N., and Tsonis, P.A. (2010). The other lives of ribosomal proteins. Hum Genomics 4, 327–344.

Biesemann, C., Grønborg, M., Luquet, E., Wichert, S.P., Bernard, V., Bungers, S.R., Cooper, B., Varoqueaux, F., Li, L., Byrne, J.A., et al. (2014). Proteomic screening of glutamatergic mouse brain synaptosomes isolated by fluorescence activated sorting. EMBO J. 33, 157–170.

Biever, A., Glock, C., Tushev, G., Ciirdaeva, E., Dalmay, T., Langer, J.D., and Schuman, E.M. (2020). Monosomes actively translate synaptic mRNAs in neuronal processes. Science 367.

Björklund, A., and Dunnett, S.B. (2007). Dopamine neuron systems in the brain: an update. Trends in Neurosciences 30, 194–202.

Bobe, R., Bredoux, R., Wuytack, F., Quarck, R., Kovàcs, T., Papp, B., Corvazier, E., Magnier, C., and Enouf, J. (1994). The rat platelet 97-kDa Ca2+ATPase isoform is the sarcoendoplasmic reticulum Ca2+ATPase 3 protein. Journal of Biological Chemistry 269, 1417–1424.

Bobrow, M.N., Litt, G.J., Shaughnessy, K.J., Mayer, P.C., and Conlon, J. (1992). The use of catalyzed reporter deposition as a means of signal amplification in a variety of formats. J. Immunol. Methods 150, 145–149.

Bolam, J.P., and Pissadaki, E.K. (2012). Living on the edge with too many mouths to feed: why dopamine neurons die. Mov. Disord. 27, 1478–1483.

Bradshaw, K.D., Emptage, N.J., and Bliss, T.V.P. (2003). A role for dendritic protein synthesis in hippocampal late LTP. Eur. J. Neurosci. 18, 3150–3152.

Bredt, D.S., and Nicoll, R.A. (2003). AMPA receptor trafficking at excitatory synapses. Neuron 40, 361–379.

Brichta, L., Shin, W., Jackson-Lewis, V., Blesa, J., Yap, E.-L., Walker, Z., Zhang, J., Roussarie, J.-P., Alvarez, M.J., Califano, A., et al. (2015). Identification of neurodegenerative factors using translatome-regulatory network analysis. Nat. Neurosci. 18, 1325–1333.

Briese, M., Saal, L., Appenzeller, S., Moradi, M., Baluapuri, A., and Sendtner, M. (2016). Whole transcriptome profiling reveals the RNA content of motor axons. Nucleic Acids Res 44, e33.

Brown, M.T.C., Henny, P., Bolam, J.P., and Magill, P.J. (2009). Activity of Neurochemically Heterogeneous Dopaminergic Neurons in the Substantia Nigra during Spontaneous and Driven Changes in Brain State. J. Neurosci. 29, 2915–2925.

Brunk, I., Blex, C., Speidel, D., Brose, N., and Ahnert-Hilger, G. (2009). Ca2+-dependent activator proteins of secretion promote vesicular monoamine uptake. J Biol Chem 284, 1050– 1056.

Burk, S.E., Lytton, J., MacLennan, D.H., and Shull, G.E. (1989). cDNA Cloning, functional expression, and mRNA Tissue Distribution of a Third Organellar Ca2+ Pump*. Journal of Biological Chemistry 264, 18561–18568.

Cai, H., Liu, G., Sun, L., and Ding, J. (2014). Aldehyde Dehydrogenase 1 making molecular inroads into the differential vulnerability of nigrostriatal dopaminergic neuron subtypes in Parkinson’s disease. Translational Neurodegeneration 3, 27.

Cajigas, I.J., Tushev, G., Will, T.J., tom Dieck, S., Fuerst, N., and Schuman, E.M. (2012). The Local Transcriptome in the Synaptic Neuropil Revealed by Deep Sequencing and High-Resolution Imaging. Neuron 74, 453–466.

Chen, B.T., and Rice, M.E. (2001). Novel Ca2+ dependence and time course of somatodendritic dopamine release: substantia nigra versus striatum. J. Neurosci. 21, 7841–7847.

Cheramy, A., Leviel, V., and Glowinski, J. (1981). Dendritic release of dopamine in the substantia nigra. Nature 289, 537–542.

Chicurel, M.E., Terrian, D.M., and Potter, H. (1993). mRNA at the synapse: analysis of a synaptosomal preparation enriched in hippocampal dendritic spines. J. Neurosci. 13, 4054– 4063.

Chu, Y., Morfini, G.A., Langhamer, L.B., He, Y., Brady, S.T., and Kordower, J.H. (2012). Alterations in axonal transport motor proteins in sporadic and experimental Parkinson’s disease. Brain 135, 2058–2073.

Costa, R.O., Martins, H., Martins, L.F., Cwetsch, A.W., Mele, M., Pedro, J.R., Tomé, D., Jeon, N.L., Cancedda, L., Jaffrey, S.R., et al. (2019). Synaptogenesis Stimulates a Proteasome-Mediated Ribosome Reduction in Axons. Cell Reports 28, 864–876.e6.

Cracco, J.B., Serrano, P., Moskowitz, S.I., Bergold, P.J., and Sacktor, T.C. (2005). Protein synthesis-dependent LTP in isolated dendrites of CA1 pyramidal cells. Hippocampus 15, 551– 556.

Crispino, M., Chun, J.T., Cefaliello, C., Perrone Capano, C., and Giuditta, A. (2014). Local gene expression in nerve endings. Dev Neurobiol 74, 279–291.

Dobin, A., Davis, C.A., Schlesinger, F., Drenkow, J., Zaleski, C., Jha, S., Batut, P., Chaisson, M., and Gingeras, T.R. (2013). STAR: ultrafast universal RNA-seq aligner. Bioinformatics 29, 15–21.

Dunn, A.R., Stout, K.A., Ozawa, M., Lohr, K.M., Hoffman, C.A., Bernstein, A.I., Li, Y., Wang, M., Sgobio, C., Sastry, N., et al. (2017). Synaptic vesicle glycoprotein 2C (SV2C) modulates dopamine release and is disrupted in Parkinson disease. Proc Natl Acad Sci U S A 114, E2253– E2262.

Ekstrand, M.I., Terzioglu, M., Galter, D., Zhu, S., Hofstetter, C., Lindqvist, E., Thams, S., Bergstrand, A., Hansson, F.S., Trifunovic, A., et al. (2007). Progressive parkinsonism in mice with respiratory-chain-deficient dopamine neurons. PNAS 104, 1325–1330.

Fusco, C.M., Desch, K., Dörrbaum, A.R., Wang, M., Staab, A., Chan, I.C.W., Vail, E., Villeri, V., Langer, J.D., and Schuman, E.M. (2021). Neuronal ribosomes dynamically exchange ribosomal proteins in a context-dependent manner. BioRxiv 2021.03.25.437026.

Geffen, L.B., Jessell, T.M., Cuello, A.C., and Iversen, L.L. (1976). Release of dopamine from dendrites in rat substantia nigra. Nature 260, 258–260.

Gene Ontology Consortium (2021). The Gene Ontology resource: enriching a GOld mine. Nucleic Acids Res 49, D325–D334.

Gervasi, N.M., Scott, S.S., Aschrafi, A., Gale, J., Vohra, S.N., MacGibeny, M.A., Kar, A.N., Gioio, A.E., and Kaplan, B.B. (2016). The local expression and trafficking of tyrosine hydroxylase mRNA in the axons of sympathetic neurons. RNA 22, 883–895.

Grammatopoulos, T.N., Jones, S.M., Ahmadi, F.A., Hoover, B.R., Snell, L.D., Skoch, J., Jhaveri, V.V., Poczobutt, A.M., Weyhenmeyer, J.A., and Zawada, W.M. (2007). Angiotensin type 1 receptor antagonist losartan, reduces MPTP-induced degeneration of dopaminergic neurons in substantia nigra. Mol Neurodegeneration 2, 1.

Gumy, L.F., Yeo, G.S.H., Tung, Y.-C.L., Zivraj, K.H., Willis, D., Coppola, G., Lam, B.Y.H., Twiss, J.L., Holt, C.E., and Fawcett, J.W. (2011). Transcriptome analysis of embryonic and adult sensory axons reveals changes in mRNA repertoire localization. RNA 17, 85–98.

Guyenet, P.G., and Crane, J.K. (1981). Non-dopaminergic nigrostriatal pathway. Brain Res 213, 291–305.

Hafner, A.-S., Donlin-Asp, P.G., Leitch, B., Herzog, E., and Schuman, E.M. (2019). Local protein synthesis is a ubiquitous feature of neuronal pre- and postsynaptic compartments. Science 364.

Hage, T.A., and Khaliq, Z.M. (2015). Tonic firing rate controls dendritic Ca2+ signaling and synaptic gain in substantia nigra dopamine neurons. J. Neurosci. 35, 5823–5836.

Hefti, F., and Lichtensteiger, W. (1978a). Dendritic dopamine: studies on the release of endogenous dopamine from subcellular particles derived from dendrites of rat nigro-striatal neurons. Neurosci. Lett. 10, 65–70.

Hefti, F., and Lichtensteiger, W. (1978b). SUBCELLULAR DISTRIBUTION OF DOPAMINE IN SUBSTANTIA NIGRA OF THE RAT BRAIN: EFFECTS OF γ-BUTYROLACTONE AND DESTRUCTION OF NORADRENERGIC AFFERENTS SUGGEST FORMATION OF PARTICLES FROM DENDRITES1. Journal of Neurochemistry 30, 1217–1230.

Hersch, S.M., Yi, H., Heilman, C.J., Edwards, R.H., and Levey, A.I. (1997). Subcellular localization and molecular topology of the dopamine transporter in the striatum and substantia nigra. Journal of Comparative Neurology 388, 211–227.

Herzog, E., Nadrigny, F., Silm, K., Biesemann, C., Helling, I., Bersot, T., Steffens, H., Schwartzmann, R., Nägerl, U.V., El Mestikawy, S., et al. (2011). In vivo imaging of intersynaptic vesicle exchange using VGLUT1 Venus knock-in mice. J. Neurosci. 31, 15544–15559.

Hobson, B.D., and Sims, P.A. (2019). Critical Analysis of Particle Detection Artifacts in Synaptosome Flow Cytometry. ENeuro 6.

Hobson, B.D., Choi, S.J., Soni, R.K., Sulzer, D., and Sims, P.A. (2021). Subcellular proteomics of dopamine neurons in the mouse brain reveals axonal enrichment of proteins encoded by Parkinson’s disease-linked genes. BioRxiv 2021.06.01.446584.

Hook, P.W., McClymont, S.A., Cannon, G.H., Law, W.D., Morton, A.J., Goff, L.A., and McCallion, A.S. (2018). Single-Cell RNA-Seq of Mouse Dopaminergic Neurons Informs Candidate Gene Selection for Sporadic Parkinson Disease. The American Journal of Human Genetics 102, 427–446.

Hoops, D., and Flores, C. (2017). Making Dopamine Connections in Adolescence. Trends Neurosci. 40, 709–719.

Houmani, J.L., Davis, C.I., and Ruf, I.K. (2009). Growth-Promoting Properties of Epstein-Barr Virus EBER-1 RNA Correlate with Ribosomal Protein L22 Binding. Journal of Virology 83, 9844–9853.

Huber, K.M., Kayser, M.S., and Bear, M.F. (2000). Role for rapid dendritic protein synthesis in hippocampal mGluR-dependent long-term depression. Science 288, 1254–1257.

Jang, M., Um, K.B., Jang, J., Kim, H.J., Cho, H., Chung, S., and Park, M.K. (2015). Coexistence of glutamatergic spine synapses and shaft synapses in substantia nigra dopamine neurons. Sci Rep 5, 14773.

Jung, H., Yoon, B.C., and Holt, C.E. (2012). Axonal mRNA localization and local protein synthesis in nervous system assembly, maintenance and repair. Nat. Rev. Neurosci. 13, 308– 324.

Juraska, J.M., Wilson, C.J., and Groves, P.M. (1977). The substantia nigra of the rat: a Golgi study. J. Comp. Neurol. 172, 585–600.

Kang, H., and Schuman, E.M. (1996). A requirement for local protein synthesis in neurotrophin-induced hippocampal synaptic plasticity. Science 273, 1402–1406.

Koopmans, F., Nierop, P. van Andres-Alonso, M., Byrnes, A., Cijsouw, T., Coba, M.P., Cornelisse, L.N., Farrell, R.J., Goldschmidt, H.L., Howrigan, D.P., et al. (2019). SynGO: An Evidence-Based, Expert-Curated Knowledge Base for the Synapse. Neuron 103, 217–234.e4.

van der Kooy, D. (1979). The organization of the thalamic, nigral and raphe cells projecting to the medial vs lateral caudate-putamen in rat. A fluorescent retrograde double labeling study. Brain Res 169, 381–387.

Korf, J., Zieleman, M., and Westerink, B.H.C. (1976). Dopamine release in substantia nigra? Nature 260, 257–258.

Kramer, D.J., Risso, D., Kosillo, P., Ngai, J., and Bateup, H.S. (2018). Combinatorial Expression of Grp and Neurod6 Defines Dopamine Neuron Populations with Distinct Projection Patterns and Disease Vulnerability. ENeuro 5.

La Manno, G., Gyllborg, D., Codeluppi, S., Nishimura, K., Salto, C., Zeisel, A., Borm, L.E., Stott, S.R.W., Toledo, E.M., Villaescusa, J.C., et al. (2016). Molecular Diversity of Midbrain Development in Mouse, Human, and Stem Cells. Cell 167, 566–580.e19.

Lammel, S., Steinberg, E.E., Földy, C., Wall, N.R., Beier, K., Luo, L., and Malenka, R.C. (2015). Diversity of Transgenic Mouse Models for Selective Targeting of Midbrain Dopamine Neurons. Neuron 85, 429–438.

Langmead, B., and Salzberg, S.L. (2012). Fast gapped-read alignment with Bowtie 2. Nat Methods 9, 357–359.

Li, H., Waites, C.L., Staal, R.G., Dobryy, Y., Park, J., Sulzer, D.L., and Edwards, R.H. (2005). Sorting of vesicular monoamine transporter 2 to the regulated secretory pathway confers the somatodendritic exocytosis of monoamines. Neuron 48, 619–633.

Liao, Y., Smyth, G.K., and Shi, W. (2014). featureCounts: an efficient general purpose program for assigning sequence reads to genomic features. Bioinformatics 30, 923–930.

Lieberman, O.J., McGuirt, A.F., Mosharov, E.V., Pigulevskiy, I., Hobson, B.D., Choi, S., Frier, M.D., Santini, E., Borgkvist, A., and Sulzer, D. (2018). Dopamine Triggers the Maturation of Striatal Spiny Projection Neuron Excitability during a Critical Period. Neuron 99, 540–554.e4.

Liu, C., Kershberg, L., Wang, J., Schneeberger, S., and Kaeser, P.S. (2018). Dopamine Secretion Is Mediated by Sparse Active Zone-like Release Sites. Cell 172, 706–718.e15.

Liu, G., Yu, J., Ding, J., Xie, C., Sun, L., Rudenko, I., Zheng, W., Sastry, N., Luo, J., Rudow, G., et al. (2014). Aldehyde dehydrogenase 1 defines and protects a nigrostriatal dopaminergic neuron subpopulation. J Clin Invest 124, 3032–3046.

Love, M.I., Huber, W., and Anders, S. (2014). Moderated estimation of fold change and dispersion for RNA-seq data with DESeq2. Genome Biology 15, 550.

Luquet, E., Biesemann, C., Munier, A., and Herzog, E. (2017). Purification of Synaptosome Populations Using Fluorescence-Activated Synaptosome Sorting. Methods Mol. Biol. 1538, 121–134.

Lüscher, C., and Huber, K.M. (2010). Group 1 mGluR-dependent synaptic long-term depression (mGluR-LTD): mechanisms and implications for circuitry & disease. Neuron 65, 445–459.

Lytton, J., Westlin, M., Burk, S.E., Shull, G.E., and MacLennan, D.H. (1992). Functional comparisons between isoforms of the sarcoplasmic or endoplasmic reticulum family of calcium pumps. Journal of Biological Chemistry 267, 14483–14489.

Maday, S., Twelvetrees, A.E., Moughamian, A.J., and Holzbaur, E.L.F. (2014). Axonal Transport: Cargo-Specific Mechanisms of Motility and Regulation. Neuron 84, 292–309.

Madisen, L., Zwingman, T.A., Sunkin, S.M., Oh, S.W., Zariwala, H.A., Gu, H., Ng, L.L., Palmiter, R.D., Hawrylycz, M.J., Jones, A.R., et al. (2010). A robust and high-throughput Cre reporting and characterization system for the whole mouse brain. Nat Neurosci 13, 133–140.

Manitt, C., Mimee, A., Eng, C., Pokinko, M., Stroh, T., Cooper, H.M., Kolb, B., and Flores, C. (2011). The netrin receptor DCC is required in the pubertal organization of mesocortical dopamine circuitry. J. Neurosci. 31, 8381–8394.

Matsuda, W., Furuta, T., Nakamura, K.C., Hioki, H., Fujiyama, F., Arai, R., and Kaneko, T. (2009). Single nigrostriatal dopaminergic neurons form widely spread and highly dense axonal arborizations in the neostriatum. J. Neurosci. 29, 444–453.

Mazaré, N., Oudart, M., Moulard, J., Cheung, G., Tortuyaux, R., Mailly, P., Mazaud, D., Bemelmans, A.-P., Boulay, A.-C., Blugeon, C., et al. (2020). Local Translation in Perisynaptic Astrocytic Processes Is Specific and Changes after Fear Conditioning. Cell Rep 32, 108076.

Melia, K.R., Trembleau, A., Oddi, R., Sanna, P.P., and Bloom, F.E. (1994). Detection and Regulation of Tyrosine Hydroxylase mRNA in Catecholaminergic Terminal Fields: Possible Axonal Compartmentalization. Experimental Neurology 130, 394–406.

Mendez, J.A., Bourque, M.-J., Fasano, C., Kortleven, C., and Trudeau, L.-E. (2011). Somatodendritic dopamine release requires synaptotagmin 4 and 7 and the participation of voltage-gated calcium channels. J. Biol. Chem. 286, 23928–23937.

Mingote, S., Chuhma, N., Kalmbach, A., Thomsen, G.M., Wang, Y., Mihali, A., Sferrazza, C., Zucker-Scharff, I., Siena, A.-C., Welch, M.G., et al. (2017). Dopamine neuron dependent behaviors mediated by glutamate cotransmission. ELife 6, e27566.

Nelson, E.L., Liang, C.L., Sinton, C.M., and German, D.C. (1996). Midbrain dopaminergic neurons in the mouse: computer-assisted mapping. J. Comp. Neurol. 369, 361–371.

Ni, J.-Q., Liu, L.-P., Hess, D., Rietdorf, J., and Sun, F.-L. (2006). Drosophila ribosomal proteins are associated with linker histone H1 and suppress gene transcription. Genes Dev. 20, 1959– 1973.

Nirenberg, M.J., Vaughan, R.A., Uhl, G.R., Kuhar, M.J., and Pickel, V.M. (1996a). The dopamine transporter is localized to dendritic and axonal plasma membranes of nigrostriatal dopaminergic neurons. J Neurosci 16, 436–447.

Nirenberg, M.J., Chan, J., Liu, Y., Edwards, R.H., and Pickel, V.M. (1996b). Ultrastructural localization of the vesicular monoamine transporter-2 in midbrain dopaminergic neurons: potential sites for somatodendritic storage and release of dopamine. J Neurosci 16, 4135–4145.

Nunes, I., Tovmasian, L.T., Silva, R.M., Burke, R.E., and Goff, S.P. (2003). Pitx3 is required for development of substantia nigra dopaminergic neurons. PNAS 100, 4245–4250.

Oh, S.W., Harris, J.A., Ng, L., Winslow, B., Cain, N., Mihalas, S., Wang, Q., Lau, C., Kuan, L., Henry, A.M., et al. (2014). A mesoscale connectome of the mouse brain. Nature 508, 207–214.

Omelchenko, N., and Sesack, S.R. (2009). Ultrastructural analysis of local collaterals of rat ventral tegmental area neurons: GABA phenotype and synapses onto dopamine and GABA cells. Synapse 63, 895–906.

Ostroff, L.E., Santini, E., Sears, R., Deane, Z., Kanadia, R.N., LeDoux, J.E., Lhakhang, T., Tsirigos, A., Heguy, A., and Klann, E. (2019). Axon TRAP reveals learning-associated alterations in cortical axonal mRNAs in the lateral amgydala. Elife 8.

Ouwenga, R., Lake, A.M., O’Brien, D., Mogha, A., Dani, A., and Dougherty, J.D. (2017). Transcriptomic Analysis of Ribosome-Bound mRNA in Cortical Neurites In Vivo. J Neurosci 37, 8688–8705.

Ouwenga, R., Lake, A.M., Aryal, S., Lagunas, T., and Dougherty, J.D. (2018). The Differences in Local Translatome across Distinct Neuron Types Is Mediated by Both Baseline Cellular Differences and Post-transcriptional Mechanisms. ENeuro 5.

Papathanou, M., Dumas, S., Pettersson, H., Olson, L., and Wallén-Mackenzie, Å. (2019). Off-Target Effects in Transgenic Mice: Characterization of Dopamine Transporter (DAT)-Cre Transgenic Mouse Lines Exposes Multiple Non-Dopaminergic Neuronal Clusters Available for Selective Targeting within Limbic Neurocircuitry. ENeuro 6.

Pereira, D.B., Schmitz, Y., Mészáros, J., Merchant, P., Hu, G., Li, S., Henke, A., Lizardi-Ortiz, J.E., Karpowicz, R.J., Morgenstern, T.J., et al. (2016). Fluorescent false neurotransmitter reveals functionally silent dopamine vesicle clusters in the striatum. Nat Neurosci 19, 578–586.

Perez, J.D., tom Dieck, S., Alvarez-Castelao, B., Tushev, G., Chan, I.C., and Schuman, E.M. (2021). Subcellular sequencing of single neurons reveals the dendritic transcriptome of GABAergic interneurons. ELife 10, e63092.

Pfeffer, M.E., Pronot, M., Angelo, M.-F., Walle, R., Cordelières, F.P., Levet, F., Claverol, S., Lacomme, S., Petrel, M., Martin, C., et al. (2020). Synaptic and supra-synaptic organisation of the dopaminergic projection to the striatum. BioRxiv 2020.02.18.952978.

Pissadaki, E.K., and Bolam, J.P. (2013). The energy cost of action potential propagation in dopamine neurons: clues to susceptibility in Parkinson’s disease. Front Comput Neurosci 7, 13.

Poulin, J.-F., Zou, J., Drouin-Ouellet, J., Kim, K.-Y.A., Cicchetti, F., and Awatramani, R.B. (2014). Defining midbrain dopaminergic neuron diversity by single-cell gene profiling. Cell Rep 9, 930–943.

Poulin, J.-F., Gaertner, Z., Moreno-Ramos, O.A., and Awatramani, R. (2020). Classification of Midbrain Dopamine Neurons Using Single-Cell Gene Expression Profiling Approaches. Trends Neurosci 43, 155–169.

Prensa, L., and Parent, A. (2001). The Nigrostriatal Pathway in the Rat: A Single-Axon Study of the Relationship between Dorsal and Ventral Tier Nigral Neurons and the Striosome/Matrix Striatal Compartments. J Neurosci 21, 7247–7260.

Raman, I.M., and Bean, B.P. (1999). Ionic currents underlying spontaneous action potentials in isolated cerebellar Purkinje neurons. J Neurosci 19, 1663–1674.

Ratai, O., Schirra, C., Rajabov, E., Brunk, I., Ahnert-Hilger, G., Chitirala, P., Becherer, U., Stevens, D.R., and Rettig, J. (2019). An Alternative Exon of CAPS2 Influences Catecholamine Loading into LDCVs of Chromaffin Cells. J Neurosci 39, 18–27.

Rayport, S., Sulzer, D., Shi, W.X., Sawasdikosol, S., Monaco, J., Batson, D., and Rajendran, G. (1992). Identified postnatal mesolimbic dopamine neurons in culture: morphology and electrophysiology. J. Neurosci. 12, 4264–4280.

Rice, M.E., and Patel, J.C. (2015). Somatodendritic dopamine release: recent mechanistic insights. Philos. Trans. R. Soc. Lond., B, Biol. Sci. 370.

Richards, C.D., Shiroyama, T., and Kitai, S.T. (1997). Electrophysiological and immunocytochemical characterization of GABA and dopamine neurons in the substantia nigra of the rat. Neuroscience 80, 545–557.

Robinson, B.G., Cai, X., Wang, J., Bunzow, J.R., Williams, J.T., and Kaeser, P.S. (2019). RIM is essential for stimulated but not spontaneous somatodendritic dopamine release in the midbrain. Elife 8.

Roy, S. (2014). Seeing the unseen: the hidden world of slow axonal transport. Neuroscientist 20, 71–81.

Sadakata, T., Mizoguchi, A., Sato, Y., Katoh-Semba, R., Fukuda, M., Mikoshiba, K., and Furuichi, T. (2004). The Secretory Granule-Associated Protein CAPS2 Regulates Neurotrophin Release and Cell Survival. J. Neurosci. 24, 43–52.

Sakers, K., Lake, A.M., Khazanchi, R., Ouwenga, R., Vasek, M.J., Dani, A., and Dougherty, J.D. (2017). Astrocytes locally translate transcripts in their peripheral processes. Proc. Natl. Acad. Sci. U.S.A. 114, E3830–E3838.

Sanz, E., Yang, L., Su, T., Morris, D.R., McKnight, G.S., and Amieux, P.S. (2009). Cell-type-specific isolation of ribosome-associated mRNA from complex tissues. Proc. Natl. Acad. Sci. U.S.A. 106, 13939–13944.

Saunders, A., Macosko, E., Wysoker, A., Goldman, M., Krienen, F., de Rivera, H., Bien, E., Baum, M., Wang, S., Goeva, A., et al. (2018). Molecular Diversity and Specializations among the Cells of the Adult Mouse Brain. Cell 174, 1015–1030.e16.

Scarnati, M.S., Kataria, R., Biswas, M., and Paradiso, K.G. (2018). Active presynaptic ribosomes in the mammalian brain, and altered transmitter release after protein synthesis inhibition. Elife 7.

Shigeoka, T., Jung, H., Jung, J., Turner-Bridger, B., Ohk, J., Lin, J.Q., Amieux, P.S., and Holt, C.E. (2016). Dynamic Axonal Translation in Developing and Mature Visual Circuits. Cell 166, 181–192.

Shigeoka, T., Koppers, M., Wong, H.H.-W., Lin, J.Q., Cagnetta, R., Dwivedy, A., de Freitas Nascimento, J., van Tartwijk, F.W., Ströhl, F., Cioni, J.-M., et al. (2019). On-Site Ribosome Remodeling by Locally Synthesized Ribosomal Proteins in Axons. Cell Rep 29, 3605–3619.e10.

Silbergeld, E.K., and Walters, J.R. (1979). Synaptosomal uptake and release of dopamine in substantia nigra: Effects of γ-aminobutyric acid and substance P. Neuroscience Letters 12, 119–126.

Simsek, D., Tiu, G.C., Flynn, R.A., Byeon, G.W., Leppek, K., Xu, A.F., Chang, H.Y., and Barna, M. (2017). The mammalian ribo-interactome reveals ribosome functional diversity and heterogeneity. Cell 169, 1051–1065.e18.

Snyder, M.E., Finlayson, M.O., Connors, T.J., Dogra, P., Senda, T., Bush, E., Carpenter, D., Marboe, C., Benvenuto, L., Shah, L., et al. (2019). Generation and persistence of human tissue-resident memory T cells in lung transplantation. Sci Immunol 4.

Subramanian, A., Tamayo, P., Mootha, V.K., Mukherjee, S., Ebert, B.L., Gillette, M.A., Paulovich, A., Pomeroy, S.L., Golub, T.R., Lander, E.S., et al. (2005). Gene set enrichment analysis: A knowledge-based approach for interpreting genome-wide expression profiles. PNAS 102, 15545–15550.

Sulzer, D. (2007). Multiple hit hypotheses for dopamine neuron loss in Parkinson’s disease. Trends Neurosci 30, 244–250.

Taylor, A.M., Berchtold, N.C., Perreau, V.M., Tu, C.H., Li Jeon, N., and Cotman, C.W. (2009). Axonal mRNA in Uninjured and Regenerating Cortical Mammalian Axons. J. Neurosci. 29, 4697–4707.

Tepper, J.M., Sawyer, S.F., and Groves, P.M. (1987). Electrophysiologically identified nigral dopaminergic neurons intracellularly labeled with HRP: light-microscopic analysis. J. Neurosci. 7, 2794–2806.

Tiklová, K., Björklund, Å.K., Lahti, L., Fiorenzano, A., Nolbrant, S., Gillberg, L., Volakakis, N., Yokota, C., Hilscher, M.M., Hauling, T., et al. (2019). Single-cell RNA sequencing reveals midbrain dopamine neuron diversity emerging during mouse brain development. Nature Communications 10, 581.

Tinevez, J.-Y., Perry, N., Schindelin, J., Hoopes, G.M., Reynolds, G.D., Laplantine, E., Bednarek, S.Y., Shorte, S.L., and Eliceiri, K.W. (2017). TrackMate: An open and extensible platform for single-particle tracking. Methods 115, 80–90.

Turiault, M., Parnaudeau, S., Milet, A., Parlato, R., Rouzeau, J.-D., Lazar, M., and Tronche, F. (2007). Analysis of dopamine transporter gene expression pattern -- generation of DAT-iCre transgenic mice. FEBS J. 274, 3568–3577.

Wang, F., Flanagan, J., Su, N., Wang, L.-C., Bui, S., Nielson, A., Wu, X., Vo, H.-T., Ma, X.-J., and Luo, Y. (2012). RNAscope: A Novel in Situ RNA Analysis Platform for Formalin-Fixed, Paraffin-Embedded Tissues. The Journal of Molecular Diagnostics 14, 22–29.

Witkovsky, P., Patel, J.C., Lee, C.R., and Rice, M.E. (2009). Immunocytochemical identification of proteins involved in dopamine release from the somatodendritic compartment of nigral dopaminergic neurons. Neuroscience 164, 488–496.

Wuytack, F., Papp, B., Verboomen, H., Raeymaekers, L., Dode, L., Bobe, R., Enouf, J., Bokkala, S., Authi, K.S., and Casteels, R. (1994). A sarco/endoplasmic reticulum Ca(2+)-ATPase 3-type Ca2+ pump is expressed in platelets, in lymphoid cells, and in mast cells. Journal of Biological Chemistry 269, 1410–1416.

Xenias, H.S., Ibáñez-Sandoval, O., Koós, T., and Tepper, J.M. (2015). Are Striatal Tyrosine Hydroxylase Interneurons Dopaminergic? J Neurosci 35, 6584–6599.

Xie, Z., Bailey, A., Kuleshov, M.V., Clarke, D.J.B., Evangelista, J.E., Jenkins, S.L., Lachmann, A., Wojciechowicz, M.L., Kropiwnicki, E., Jagodnik, K.M., et al. (2021). Gene Set Knowledge Discovery with Enrichr. Current Protocols 1, e90.

Yamada, T., McGeer, P.L., Baimbridge, K.G., and McGeer, E.G. (1990). Relative sparing in Parkinson’s disease of substantia nigra dopamine neurons containing calbindin-D28K. Brain Res 526, 303–307.

Younts, T.J., Monday, H.R., Dudok, B., Klein, M.E., Jordan, B.A., Katona, I., and Castillo, P.E. (2016). Presynaptic Protein Synthesis Is Required for Long-Term Plasticity of GABA Release. Neuron 92, 479–492.

Zampese, E., and Surmeier, D.J. (2020). Calcium, Bioenergetics, and Parkinson’s Disease. Cells 9, 2045.

