## Supplementary Figures for "Subcellular and regional localization of mRNA translation in midbrain dopamine neurons"

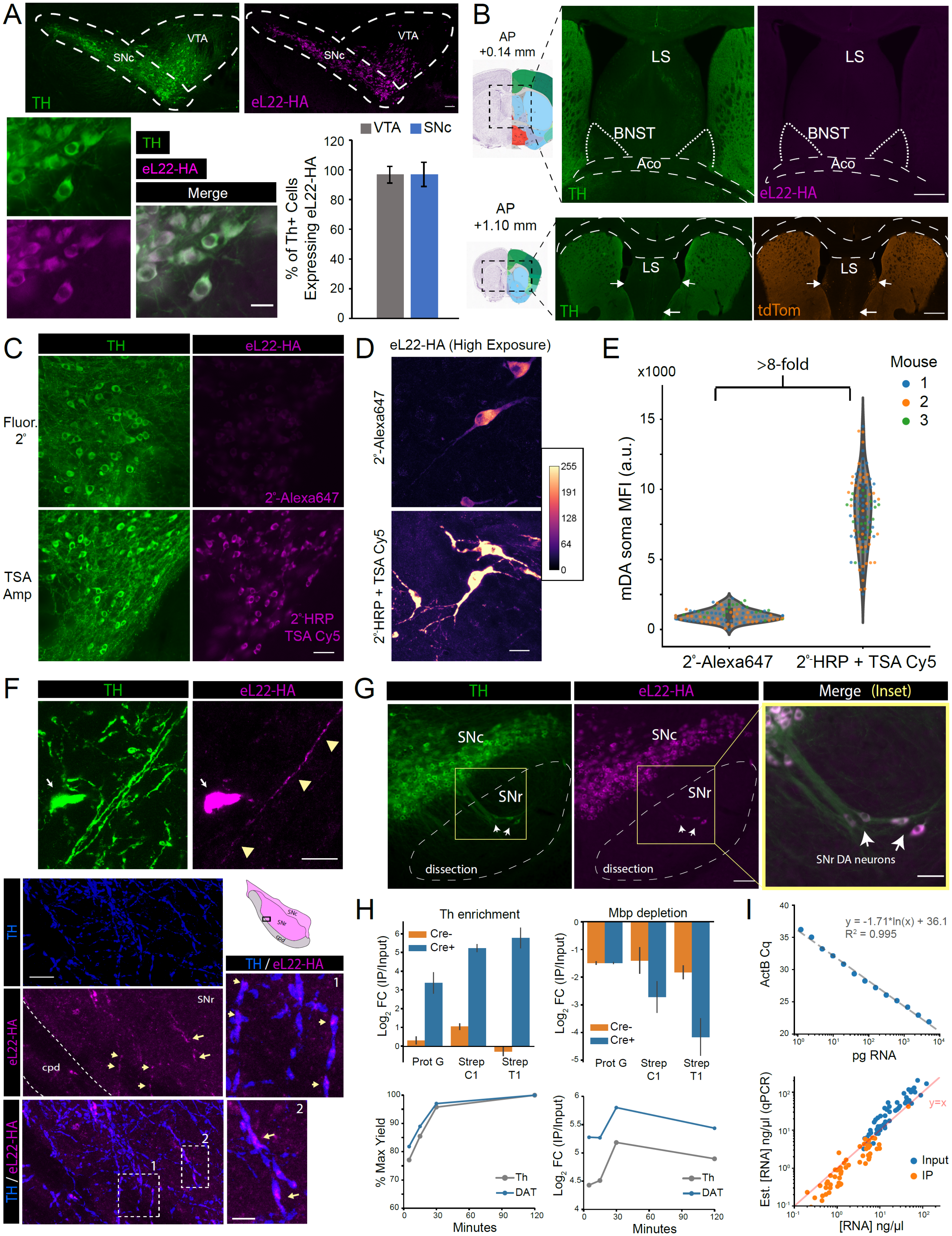

**Figure S1: Histological analysis of eL22-HA expression in DAT$^{IRES\text{-}Cre}$:RiboTag mice, eL22-HA signal amplification, eL22-HA staining in distal SNr dendrites of mDA neurons, RiboTag IP optimization, and qRT-PCR estimation of RiboTag IP yield**

**(A)** *Upper*: Epifluorescence image of DAT$^{IRES\text{-}Cre}$:RiboTag midbrain immunostained for tyrosine hydroxylase (TH) and eL22-HA. Dashed lines indicate regions used for cell counting shown below. *Lower left*: SNc of DAT$^{IRES\text{-}Cre}$:RiboTag, representative of images for cell counting. Scale bar, 20 µm. *Lower right*: % of TH$^{+}$ cells expressing eL22-HA at indicated ages. Data represent mean ± standard deviation, and are derived from n=2-3 mice and n=6 fields, (VTA) n= 717 cells; (SNc) n=451 cells. Mean is >95% in both regions. Scale bar, 100 µm.

**(B)** *Upper*: DAT$^{IRES\text{-}Cre}$:RiboTag midbrain stained for TH and eL-22HA at the indicated coordinates. *Lower*: DAT$^{IRES\text{-}Cre}$:Ai9 section stained for TH and tdTomato at indicated coordinates. Both scale bars, 500 µm.

**(C)** Comparison of AlexaFluor647-conjugated secondary antibody vs. TSA-Cy5 + HRP-conjugated secondary antibody for eL22-HA immunostaining. Exposure was optimized to avoid saturation of soma. Scale bar, 50 µm.

**(D)** Same as panel A, but using high exposure to detect dendritic labeling (somata are saturated for TSA-Cy5). Scale bar, 25 µm.

**(E)** Quantification of somata eL22-HA intensity, related to panel A. Violin plot depicts n=100 soma quantified from 2-3 fields each from 3 mice.

**(F)** *Upper*: eL22-HA labeling in descending SNr dendrites can be distinguished from soma of SNr mDA neurons. Scale bar, 20 µm. *Lower*: Amplified eL22-HA labeling is observed in distal SNr mDA neuronal dendrites near the cerebral peduncle. Scale bar, 15 µm. Inset scale bar, 5 µm.

**(G)** DAT$^{IRES\text{-}Cre}$:RiboTag midbrain immunostained for TH and eL22-HA. Scale bar, 200 µm. *Arrows*: A few scattered Th+/eL22-HA+ mDA neurons are present in the SNr. Scale bar, 50 µm.

**(H)** Optimization of eL22-HA IP conditions. *Upper*: Cre-negative or Cre-positive ventral midbrain (n=2 each genotype) lysate was split into three equal parts for capture with Rabbit anti-HA (Protein G beads) or biotinylated Rabbit anti-HA (Streptavidin C1/T1 beads). Mean and confidence intervals are plotted. *Lower*: Further optimization of capture time using four equal parts of DAT$^{IRES\text{-}Cre}$:RiboTag ventral midbrain lysate with biotinylated Rb anti-HA and Streptavidin T1 beads. Antibody was incubated with polysome lysates overnight, and beads were added for the capture time indicated.

**(I)** *Upper:* qRT-PCR of beta-Actin (ActB) with total RNA input amounts (measured by Qubit) ranging from 1 pg – 5 ng. *Lower:* qRT-PCR (ActB) estimation of RNA concentration (measured by Qubit for Input samples, or RNA Pico bioanalyzer for IP samples, n=40-48 each).

*Abbreviations*: SNc, Substantia nigra pars compacta; SNr, Substantia nigra parts reticulata; VTA, Ventral tegmental area, Aco, Anterior commissure; LS, Lateral septum; BNST, Bed nucleus of the stria terminalis, TSA; tyramide signal amplification, HRP; horseradish peroxidase, cpd; cerebral peduncle, Th; tyrosine hydroxylase, Mbp; myelin basic protein, DAT; dopamine transporter, ActB; beta-actin.

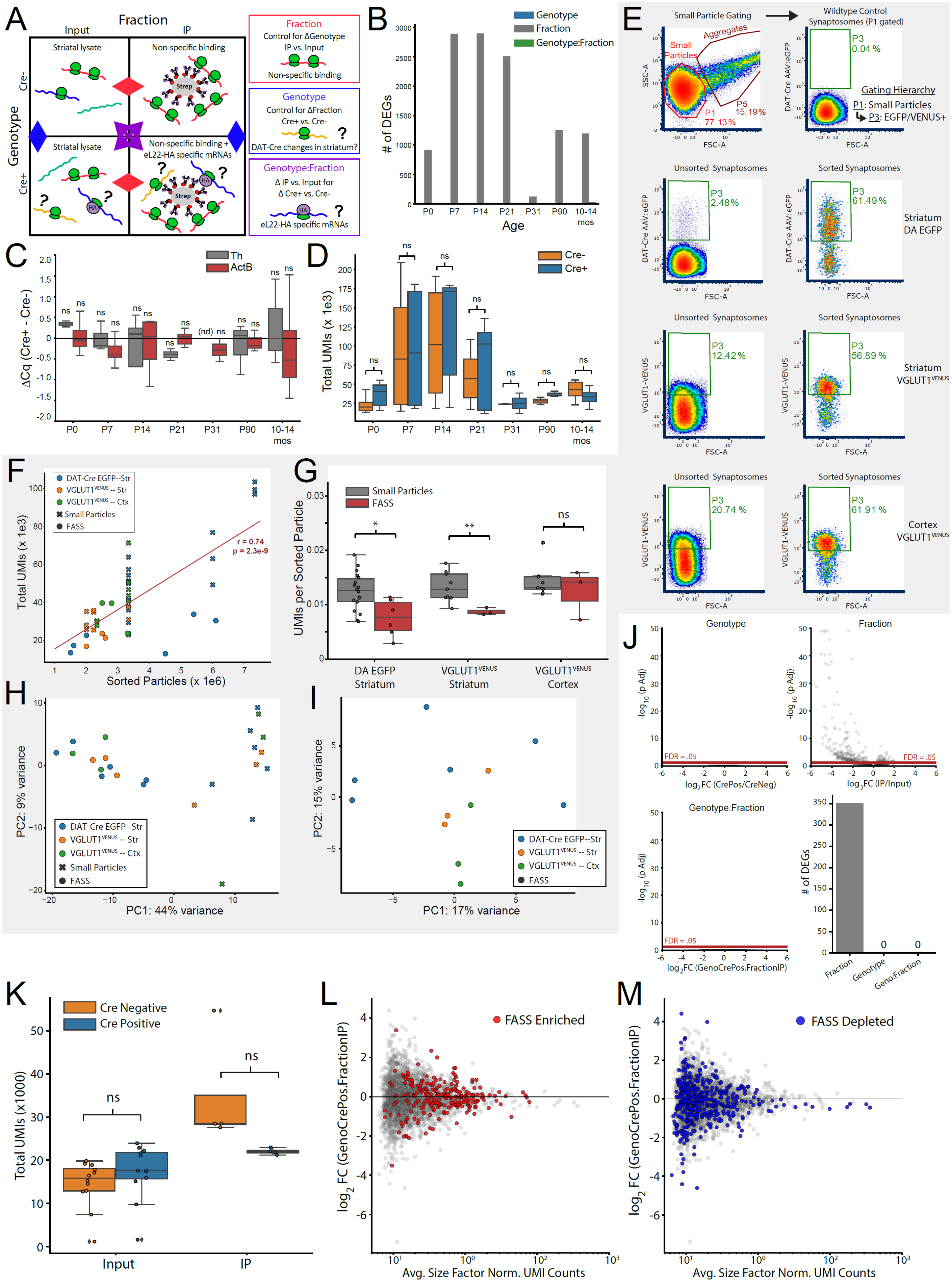
A
Fraction
Input
IP
Striatal lysate
Non-specific binding
Cre-
Genotype
Cre+
Striatal lysate
Non-specific binding +
eL22-HA specific mRNAs
Strep
HA
Fraction
Control for ΔGenotype
IP vs. Input
Non-specific binding
Genotype
Control for ΔFraction
Cre+ vs. Cre-
DAT-Cre changes in striatum?
Genotype:Fraction
Δ IP vs. Input for
Δ Cre+ vs. Cre-
eL22-HA specific mRNAs
B
Genotype
Fraction
Genotype:Fraction
### of DEGs
Age
P0
P7
P14
P21
P31
P90
10-14 mos
C
Th
ActB
ΔCq (Cre+ - Cre-)
ns
(nd)
D
Cre-
Cre+
Total UMIs (x 1e3)
E
Small Particle Gating
Wildtype Control
Synaptosomes (P1 gated)
Aggregates
Small Particles
P1 77.13%
P5 15.19%
P3 0.04 %
Gating Hierarchy
P1: Small Particles
P3: EGFP/VENUS+
SSC-A
FSC-A
DAT-Cre AAV:eGFP
Unsorted Synaptosomes
Sorted Synaptosomes
P3 2.48%
P3 61.49 %
Striatum DA EGFP
VGLUT1-VENUS
P3 12.42%
P3 56.89 %
Striatum VGLUT1 VENUS
P3 20.74 %
P3 61.91 %
Cortex VGLUT1 VENUS
F
DAT-Cre EGFP--Str
VGLUT1 VENUS -- Str
VGLUT1 VENUS -- Ctx
Small Particles
FASS
r = 0.74
p = 2.3e-9
Total UMIs (x 1e3)
Sorted Particles (x 1e6)
G
Small Particles
FASS
UMIs per Sorted Particle
*
**
ns
DA EGFP Striatum
VGLUT1 VENUS Striatum
VGLUT1 VENUS Cortex
H
PC2: 9% variance
PC1: 44% variance
I
PC2: 15% variance
PC1: 17% variance
J
Genotype
Fraction
Genotype:Fraction
-log10 (p Adj)
FDR = .05
log2FC (CrePos/CreNeg)
log2FC (IP/Input)
log2FC (GenoCrePos.FractionIP)
### of DEGs
Fraction
Genotype
Geno:Fraction
K
Cre Negative
Cre Positive
Total UMIs (x1000)
Input
IP
L
FASS Enriched
log2 FC (GenoCrePos.FractionIP)
Avg. Size Factor Norm. UMI Counts
M
FASS Depleted

**Figure S2: RNA-Seq analysis of bulk striatal and striatal synaptosome RiboTag IP samples, VGLUT1$^{VENUS}$ and DA:EGFP FASS Gating, and RNA-Seq analysis of FASS samples**

**(A)** Schematic depicting generalized linear model analysis of striatal RiboTag IP using the *DESeq2* likelihood ratio test (LRT). Analogous to a 2-way analysis of variance, the main effects of each factor and the interaction between them are each tested for statistical significance across all genes (with FDR correction). See **Methods**.

**(B)** Number of genes meeting statistical significance (FDR < 0.05) in the *DESeq2* LRT with the indicated terms omitted from the following reduced model: *~genotype + fraction + genotype:fraction*. In addition to testing each age was independently (*shown here*), the effects of *age*, *fraction*, *genotype*, and *fraction:genotype* interaction were also on the entire dataset across all ages (see **Figure 3A-B**). In contrast to the genotype-independent effect of *fraction* (non-specific binding), the effects of *genotype* and *genotype:fraction* interaction were negligible at all ages. See **Supplementary File 5** for complete summary of *DESeq2* analysis. Mice per age and genotype: (P0, Cre-) n= 6, (P0, Cre+) n=6, (P7, Cre-) n=6, (P7, Cre+) n=6, (P14, Cre-) n=6, (P14, Cre+) n=6, (P21, Cre-) n=6, (P21, Cre+) n=7, (P31, Cre-) n=2, (P31, Cre+), n=2, (P90, Cre-) n=2, (P90, Cre+), n=3, (10-14 mo., Cre-) n=6, (10-14 mo., Cre+) n=4, where each n indicates an IP and corresponding Input sample.

**(C)** qRT-PCR measurement of *Th* and *ActB* mRNA yield in striatal RiboTag IPs from Cre-negative and Cre-positive mice. The difference between the average Cq values (Cre-positive Cq – Cre-negative Cq) is plotted at each age. Mice per age and genotype are the same as in panel B. nd: not detectable, ns: all comparisons are not significant (p > 0.05), Welch's unequal variance t-test.

**(D)** Total UMIs per sample for striatal RiboTag IPs from Cre-negative and Cre-positive mice. Mice per age and genotype are the same as in panel B. ns: all comparisons are not significant (p > 0.05), Welch's unequal variance t-test.

**(E)** Density plots of Fluorescence-Activated Synaptosome Sorting (FASS) gating strategy. *<u>Upper</u>*: Particles are first selected from the 'P1' gate on forward and side scatter in order to avoid aggregates. Synaptosomes from wildtype controls are used to set a fluorescence threshold on which to sort VGLUT1$^{VENUS}$ and DA:EGFP particles. *<u>Lower</u>*: Representative density plots of Unsorted (*left*) and Sorted (*right*) synaptosomes from the indicated regions and genotypes.

**(F)** Number of sorted particles vs. total UMIs for the indicated sorted samples (n=6 striatum DA:EGFP, n=3 striatum VGLUT1$^{VENUS}$, n=3 cortex VGLUT1$^{VENUS}$, where each n represents both a FASS sample and three corresponding small particle samples). Pearson's r = 0.74, p = 2.3e-09.

**(G)** Total UMIs per sorted particle for small particle and FASS samples as indicated. * indicates p < 0.05, ** indicates p < 0.01, Welch's unequal variance t-test. Samples are the same as shown in panel F.

**(H)** PCA of *DESeq2 rlog* normalized UMI counts for FASS and small particle samples. Samples are the same as shown in panel F, but small particle technical replicates (n=3 per FASS sample) were collapsed.

**(I)** Same as panel H, but only FASS samples are included in the PCA. See **Supplementary File 6** for complete summary of *DESeq2* testing.

**(J)** *<u>Upper and lower left:</u>* Volcano plots are derived from the *DESeq2* LRT, with the indicated terms removed from the following two-factor GLM: ~ *genotype* + *fraction* + *genotype:fraction*. *<u>Lower right:</u>* Number of differentially expressed genes (DEGs, FDR < 0.05) from the *DESeq2* LRT test for the indicated factors, related to panel A. See **Supplementary File 8** for complete summary of *DESeq2* testing.

**(K)** Total UMIs for Input and IP samples from striatal synaptosome RiboTag IPs (n = 12 each genotype for Input, n = 4 each genotype for IP). ns indicates p > 0.05, Welch's unequal variance t-test.

**(L-M)** log2 fold change vs. abundance (MA) plot for FASS-enriched or FASS-depleted genes shown in **Figure 3G.** $Log_2$(GenoCrePos.FractionIP) represents the difference in the *fraction* effect between genotypes: { Cre-positive $log_2$FC(IP/Input) – Cre-negative $log_2$FC(IP/Input) }.

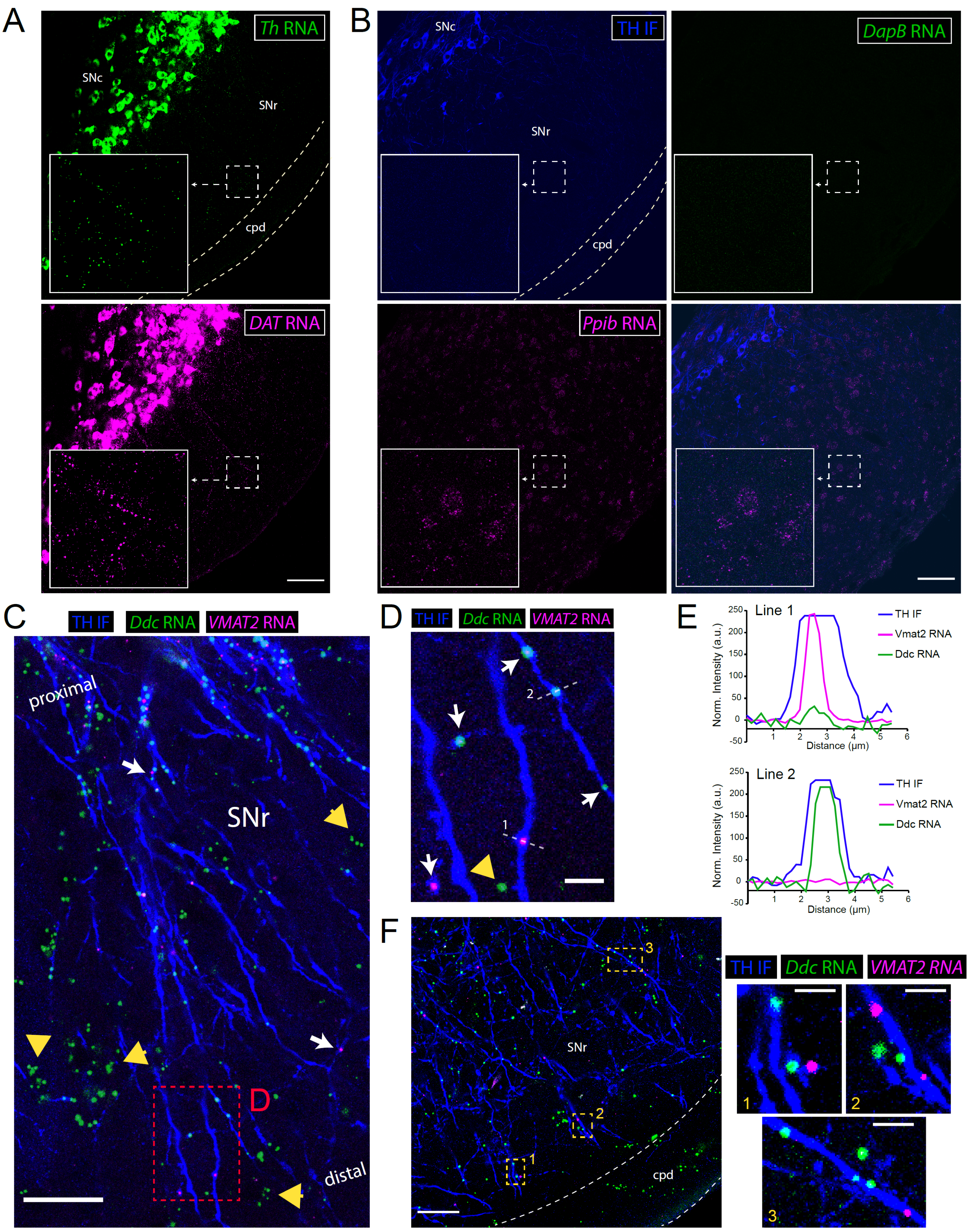

A
Th RNA
SNc
SNr
cpd
DAT RNA
B
TH IF
SNc
SNr
cpd
DapB RNA
Ppib RNA
C
TH IF
Ddc RNA
VMAT2 RNA
proximal
SNr
D
distal
D
TH IF
Ddc RNA
VMAT2 RNA
2
1
E
Line 1
TH IF
Vmat2 RNA
Ddc RNA
Norm. Intensity (a.u.)
Distance (μm)
Line 2
TH IF
Vmat2 RNA
Ddc RNA
Norm. Intensity (a.u.)
Distance (μm)
F
3
SNr
2
1
cpd
TH IF
Ddc RNA
VMAT2 RNA
1
2
3

**Figure S3: Midbrain FISH for dopaminergic and control mRNAs, *Ddc* and *Slc18a2/Vmat2* mRNA in TH$^+$ SNr dendrites**

**(A)** Multicolor fluorescence *in situ* hybridization (FISH, RNAScope assay) for *Th* and *Slc6a3/DAT* mRNA in the substantia nigra. Scale bar, 100 μm.

**(B)** TH immunostaining combined with multicolor FISH for the negative control (bacterial) mRNA *DapB* and positive control mRNA *Ppib*. Scale bar, 100 μm.

**(C)** TH immunostaining combined with multicolor FISH for *Ddc* and *Slc18a2/Vmat2* mRNA in the SNr. Yellow large arrowheads indicate clusters of *Ddc* mRNA outside of TH+ dendrites. White arrows indicate *Slc18a2/Vmat2* mRNA within TH$^+$ dendrites. Red dashed lines indicate the inset in panel D. Scale bar, 25 μm.

**(D)** Inset corresponding to red dashed lines in panel C. Yellow large arrowhead indicates a *Ddc* mRNA puncta outside of TH+ dendrites. White arrows indicate *Ddc* and *Slc18a2/Vmat2* mRNA within TH$^+$ dendrites. White dashed lines correspond to the intensity profiles shown below in panel E. Scale bar, 5 μm.

**(E)** Intensity profiles for all three fluorescent channels, corresponding to the lines indicated above in panel D.

**(F)** Left: TH immunostaining combined with multicolor FISH for *Ddc* and *Slc18a2/Vmat2* mRNA in the distal SNr near the cerebral peduncle (cpd). Yellow dashed lines indicate the insets shown on the right. Scale bar, 25 μm. *Right:* Insets of *Ddc* and *Slc18a2/Vmat2* mRNA in TH$^+$ dendrites. Scale bars, 5 μm.

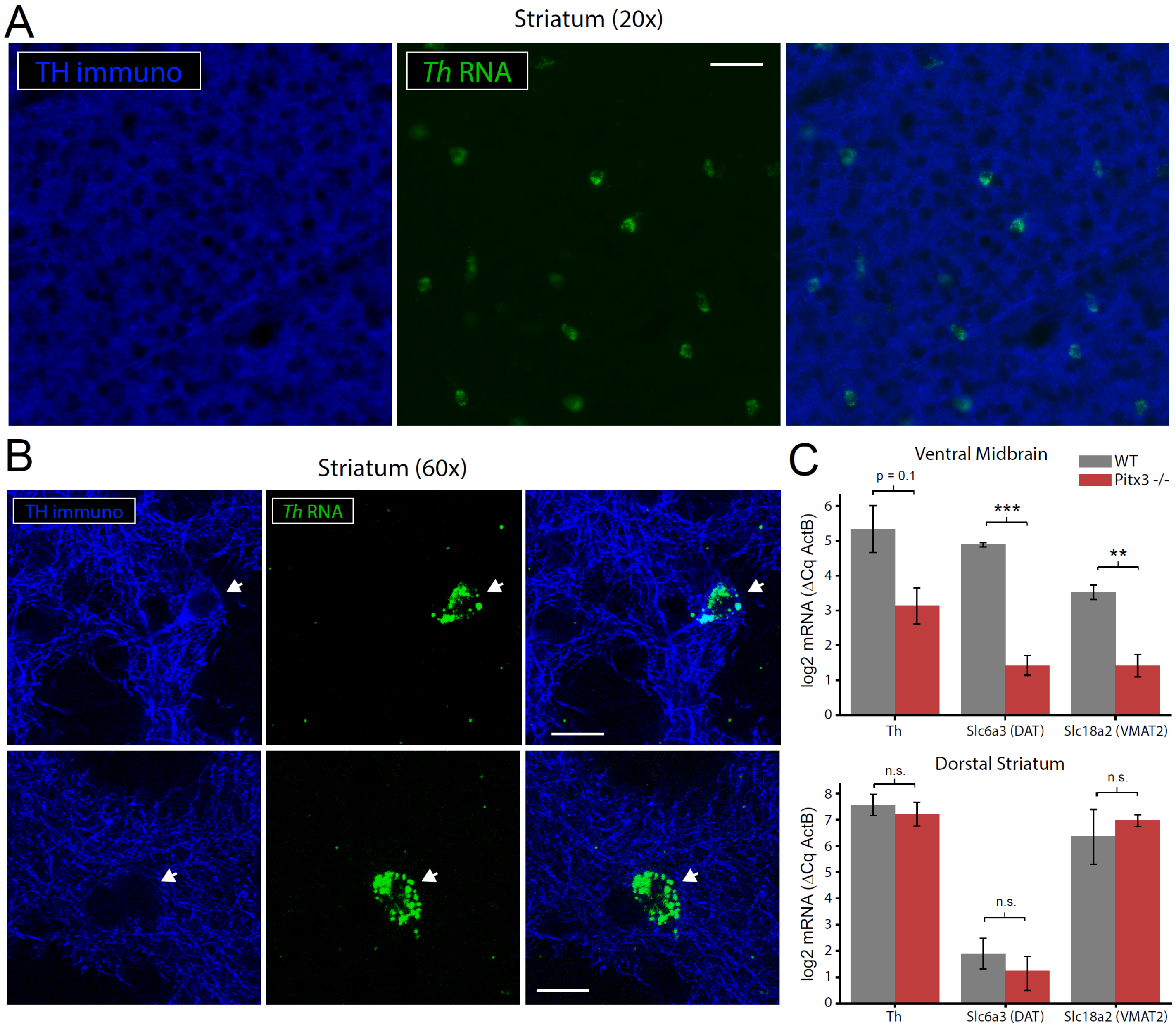


**Figure S4: *Th* mRNA⁺ striatal neurons, not dopaminergic axons, are the source of *Th* mRNA in the striatum**

**(A)** TH immunostaining combined with FISH for *Th* mRNA in the striatum. Scale bar, 50 µm.

**(B)** TH immunostaining combined with FISH for *Th* mRNA in the striatum. Occasionally, some *Th* mRNA$^+$ neurons also display TH immunoreactivity (*upper*). Typically, they do not (*lower*). Scale bars, 15 µm.

**(C)** qRT-PCR of the indicated dopaminergic mRNAs in the Ventral Midbrain (*upper*) or dorsal striatum (*lower*) of Wildtype (n=3) and Pitx3-/- mice (n=4). ** indicates $p < 0.01$, *** indicates $p < 0.001$, Welch's unequal variance t-test.

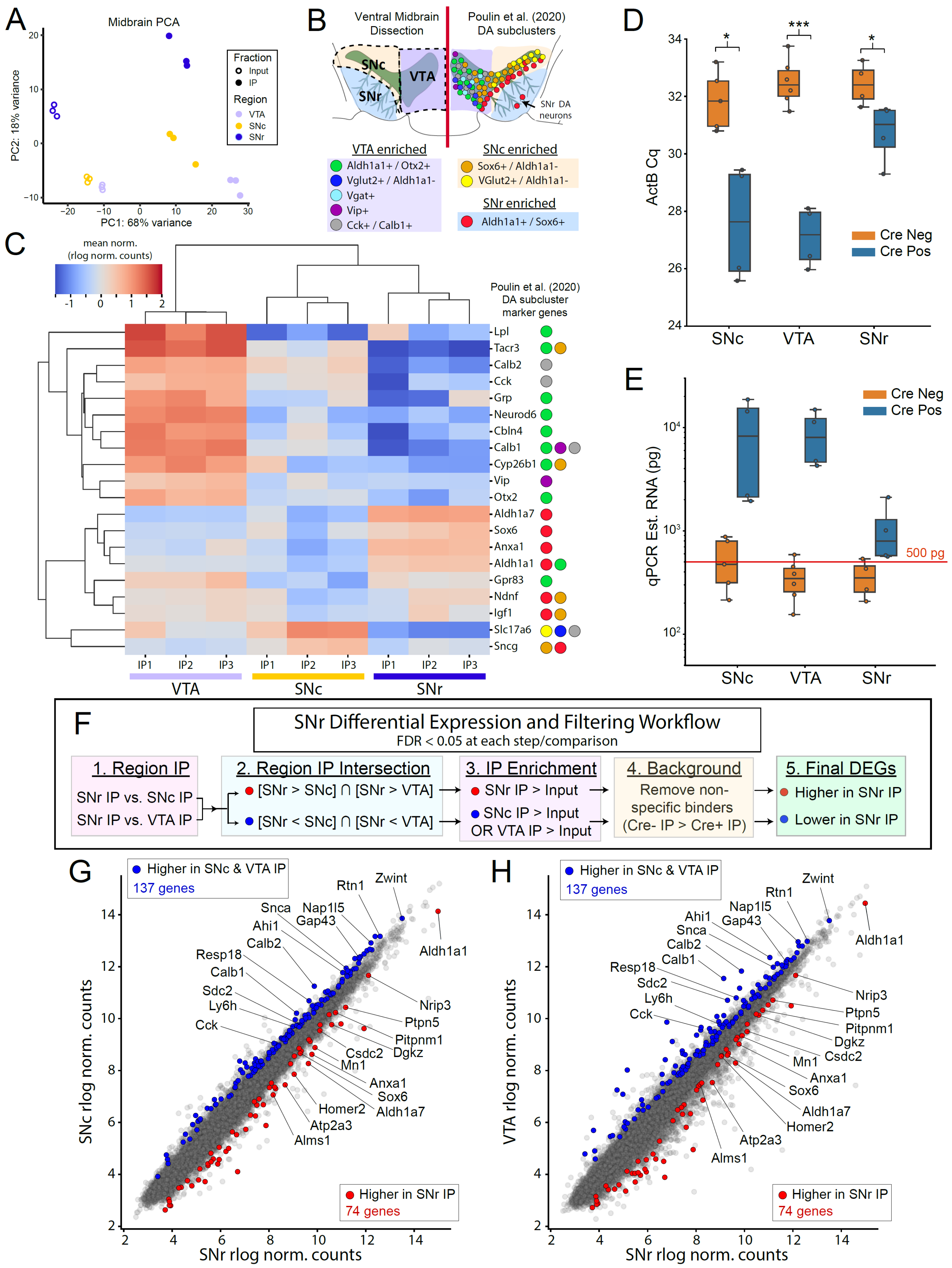
A
Midbrain PCA
PC2: 18% variance
PC1: 68% variance
Fraction
Input
IP
Region
VTA
SNc
SNr
B
Ventral Midbrain Dissection
Poulin et al. (2020) DA subclusters
SNc
VTA
SNr
SNr DA neurons
VTA enriched
Aldh1a1+ / Otx2+
Vglut2+ / Aldh1a1-
Vgat+
Vip+
Cck+ / Calb1+
SNc enriched
Sox6+ / Aldh1a1-
VGlut2+ / Aldh1a1-
SNr enriched
Aldh1a1+ / Sox6+
C
mean norm. (rlog norm. counts)
Poulin et al. (2020) DA subcluster marker genes
Lpl
Tacr3
Calb2
Cck
Grp
Neurod6
Cbln4
Calb1
Cyp26b1
Vip
Otx2
Aldh1a7
Sox6
Anxa1
Aldh1a1
Gpr83
Ndnf
Igf1
Slc17a6
Sncg
IP1 IP2 IP3
VTA
SNc
SNr
D
ActB Cq
Cre Neg
Cre Pos
SNc
VTA
SNr
E
qPCR Est. RNA (pg)
500 pg
Cre Neg
Cre Pos
SNc
VTA
SNr
F
SNr Differential Expression and Filtering Workflow
FDR < 0.05 at each step/comparison
1. Region IP
SNr IP vs. SNc IP
SNr IP vs. VTA IP
2. Region IP Intersection
[SNr > SNc] ∩ [SNr > VTA]
[SNr < SNc] ∩ [SNr < VTA]
3. IP Enrichment
SNr IP > Input
SNc IP > Input OR VTA IP > Input
4. Background
Remove non-specific binders (Cre- IP > Cre+ IP)
5. Final DEGs
Higher in SNr IP
Lower in SNr IP
G
Higher in SNc & VTA IP
137 genes
Higher in SNr IP
74 genes
SNc rlog norm. counts
SNr rlog norm. counts
Rtn1
Zwint
Snca
Nap1l5
Gap43
Ahi1
Calb2
Resp18
Calb1
Sdc2
Ly6h
Cck
Aldh1a1
Nrip3
Ptpn5
Pitpnm1
Dgkz
Csdc2
Mn1
Anxa1
Sox6
Aldh1a7
Homer2
Atp2a3
Alms1
H
Higher in SNc & VTA IP
137 genes
Higher in SNr IP
74 genes
VTA rlog norm. counts
SNr rlog norm. counts
Zwint
Rtn1
Nap1l5
Gap43
Ahi1
Snca
Calb2
Calb1
Resp18
Sdc2
Ly6h
Cck
Aldh1a1
Nrip3
Ptpn5
Pitpnm1
Dgkz
Csdc2
Mn1
Anxa1
Sox6
Aldh1a7
Homer2
Atp2a3
Alms1

**Figure S5: RiboTag IP from midbrain dissections recapitulates mDA neuronal heterogeneity, qRT-PCR estimated yield of midbrain RiboTag IPs, SNr RiboTag IP Filtering, and Enrichment of *Aldh1a1*⁺/*Sox6*⁺ mDA neuronal markers**

**(A)** PCA of RiboTag IP and Input samples from VTA, SNc, and SNr dissections.

**(B)** Schematic depicting anatomical dissections and the anatomical distribution of mDA neuronal clusters described by Poulin et al. (2020).

**(C)** Clustered heatmap of *DESeq2 rlog* normalized counts, mean-normalized within each gene, for the indicated VTA, SNc, and SNr RiboTag IPs (n=3 each). The twenty genes shown are strong markers for specific mDA neuronal clusters as shown in panel B and in Poulin et al. (2020).

**(D)** beta-Actin (ActB) Cq values for Cre-negative and Cre-positive RiboTag IPs in the indicated regions (n=4 each region/genotype). * indicates $p < 0.05$, *** indicates $p < 0.001$, Welch's unequal variance t-test.

**(E)** Estimated total RNA (picograms) based on ActB Cq values. Red line indicates 500 picograms.

**(F)** Schematic depicting filtering and comparison of SNr RiboTag IP to VTA and SNc RiboTag IPs (FDR < 0.05 at each step). First, DEGs in SNr vs. SNc and in SNr vs. VTA comparisons are identified. Second, the intersection of SNr-enriched or SNr-depleted genes (relative to SNc/VTA) from these two DEG lists is retained. Third, only genes enriched in SNr IP vs. Input or SNc/VTA IP vs. Input comparisons are retained. Fourth, genes that are significantly higher in Cre-negative IP samples compared to Cre-positive IP samples are removed (non-specific binders). The final list of DEGs includes genes enriched (red) or depleted (blue) in SNr RiboTag IPs relative to SNc and VTA RiboTag IPs. See **Supplementary File 9** for complete summary of DEGs and filtering.

**(G-H)** Log-log plots depicting the average *DESeq2 rlog* normalized counts for SNr RiboTag IPs (x-axis) and SNc (*left*) or VTA (*right*) RiboTag IPs (y-axis). DEGs corresponding to panel F are labeled in blue or red, with select genes labeled.

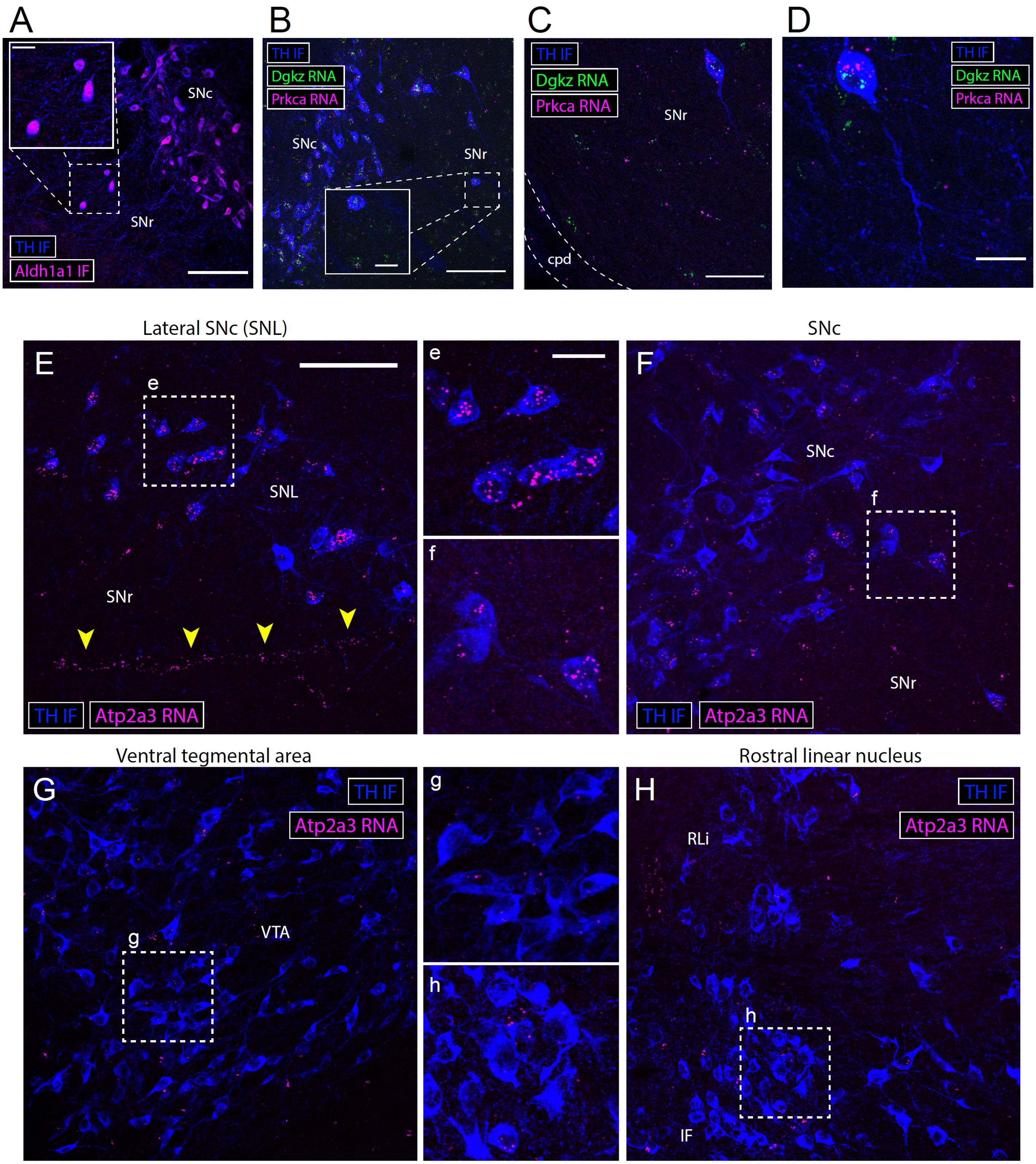


**Figure S6: SNr RiboTag IP-enriched mRNAs localized within SNr mDA somata and heterogeneous expression of *Atp2a3* (SERCA3) in mDA neurons**

**(A)** Immunostaining for TH and Aldh1a1 reveals Aldh1a1$^{+}$ mDA neurons within the SNr. Scale bar, 100 µm. Inset scale bar, 20 µm.

**(B-D)** TH immunostaining combined with multicolor FISH for *Dgkz* and *Prkca* mRNA in the proximal and distal SNr. Both of these SNr RiboTag IP-enriched mRNAs are localized within the soma of mDA neurons in the SNr, and are not distributed in dopaminergic dendrites. **(B)** Scale bar, 100 µm. Inset scale bar, 20 µm. **(C)** Scale bar, 50 µm. **(D)** Scale bar, 20 µm.

**(E-H)** TH immunostaining combined with FISH for *Atp2a3/SERCA3* mRNA in the indicated regions. Dashed white lines indicate the insets shown in the center (e-h). Yellow arrowheads in **(E)** indicate prominent labeling of blood vessels, likely within endothelial cells which are known to express SERCA3. Scale bar, 100 µm. Inset scale bar, 25 µm. See **Figure 5G** for quantification within each region.

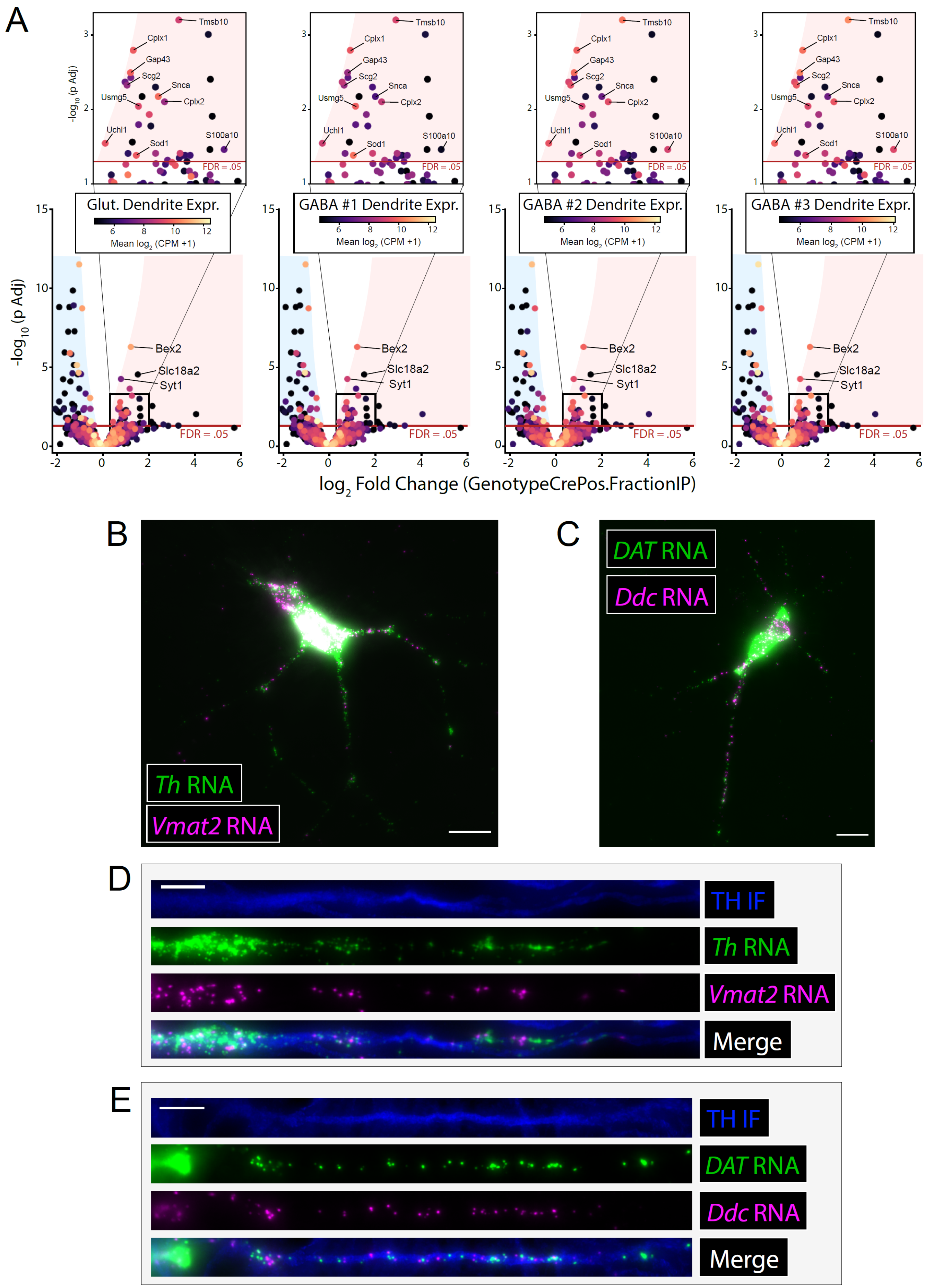
A
Glut. Dendrite Expr.
GABA #1 Dendrite Expr.
GABA #2 Dendrite Expr.
GABA #3 Dendrite Expr.
Mean log2 (CPM +1)
-log10 (p Adj)
Tmsb10
Cplx1
Gap43
Scg2
Snca
Usmg5
Cplx2
Uchl1
Sod1
S100a10
FDR = .05
Bex2
Slc18a2
Syt1
log2 Fold Change (GenotypeCrePos.FractionIP)
B
Th RNA
Vmat2 RNA
C
DAT RNA
Ddc RNA
D
TH IF
Th RNA
Vmat2 RNA
Merge
E
TH IF
DAT RNA
Ddc RNA
Merge

**Figure S7: mRNAs encoding presynaptic proteins are also present in hippocampal dendrites, and dendritic localization of dopaminergic mRNAs in cultured mDA neurons**

**(A)** Volcano plots showing the Cre-dependent log2 fold change in IP vs. Input (Cre-positive – Cre-negative) for midbrain synaptosomal RiboTag IPs (same as **Figure 6E**) with genes colored by their expression level in the dendrites of four subsets of hippocampal neurons (Perez et al., 2021).

**(B-C)** Multicolor FISH for the indicated dopaminergic mRNAs in cultured mDA neurons. Scale bars, 20 µm.

**(D-E)** TH immunostaining combined with multicolor FISH for the indicated dopaminergic mRNAs in cultured mDA neurons. Straightened dendritic segments are displayed. Scale bars, 10 µm.
